# An unexpected mode of whole-body regeneration from reaggregated cell suspension in *Hydractinia* (Cnidaria, Hydrozoa)

**DOI:** 10.1101/2025.01.06.631080

**Authors:** Camille Curantz, Gabriel Krasovec, Helen R Horkan, Laura Ryan, Áine Varley, Uri Frank

## Abstract

Hydrozoan cnidarians are among the few animals that can regenerate whole bodies from reaggregated cell dissociations but the cellular and molecular mechanisms that control this ability and how it is related to embryonic development are not well understood. Furthermore, the evolution of this type of regeneration is enigmatic since it does not occur naturally. Here, we show that aggregate regeneration in *Hydractinia symbiolongicarpus* proceeds through several, consistent stages that include the formation of an epidermal layer, followed by migration, proliferation, and differentiation of adult pluripotent stem cells, known as i-cells. Migration of i-cells is controlled by sphingosine-1-phosphate signaling. Single-cell transcriptomics revealed, surprisingly, that the newly regenerated individual derives nearly exclusively from i-cell progeny rather than from recycled somatic cells, as seen in other hydrozoans. Given the similarity of this phenomenon to embryogenesis, we propose that the ability of *Hydractinia* cell aggregates to regenerate is a side effect of the animal’s i-cell-mediated development.

## INTRODUCTION

Animals exhibit a wide range of regenerative abilities and processes ^1–3^, which differ even between closely related taxa ^4,5^. This probably reflects a complex evolutionary history that included multiple gains and losses of regeneration in distinct clades. Nevertheless, with some notable exceptions, a general trend exists in metazoans such that animals with simpler body plans tend to enjoy a better regenerative ability than those with a more complex morphology. This is probably because intricate morphologies require a hierarchical, stepwise assembly of structures that are more difficult to repair if partially lost. Furthermore, complex animals require greater cell fate stability to maintain ‘law-and-order’ in their elaborate organs, while simpler animals can afford to maintain uncommitted cells in their tissues.

It is therefore not surprising that hydrozoan cnidarians, a group of animals that are characterized by a simple body structure, are among the best regenerators in the animal kingdom. Studied members of the Hydrozoa can not only regrow any lost body part, but also possess the rare ability to reassemble an intact, functional adult from unstructured, mixed aggregates of cells obtained from dissociated tissues ^6,7^. This remarkable capability has been most intensively studied in the freshwater, solitary hydrozoan *Hydra* ^8^. It has been shown that *Hydra* cell aggregates self-sort the two epithelial body layers ^9^, followed by *de novo* establishment of a WNT-driven organizer region that drive the development of the new body axis ^10^. A model of selective epithelialization was proposed to explain how external vs. internal cells sort ^11^. Individual neurons, which survived the dissociation/reaggregation procedure, progressively re-synchronize their activities into functional ensembles ^12^. However, the molecular pathways that control aggregate regeneration remains poorly understood. Furthermore, given hydrozoan diversity, it would be interesting to comparatively study the process in other members of the group.

We have studied whole-body regeneration of dissociated, reaggregated cells in *Hydractinia symbiolongicarpus*. *Hydractinia* is a colonial, clonal marine hydrozoan. Studying regeneration in this animal is attractive because its mode of head regeneration differs from the one of *Hydra* ^13^. Moreover, *Hydractinia* possesses a population of adult pluripotent cells, known as i-cells, that contribute to all somatic lineages and to gametes ^14^; this is also different from *Hydra* i-cells that are only multipotent, unable to form epithelia ^15^. We find that aggregate regeneration in *Hydractinia* is indeed fundamentally different from its *Hydra* counterpart. *Hydractinia* aggregates generate a temporary epithelium, enclosing a mass of cell debris. Guided by sphingosine signaling, i-cells migrate to foci where they coalesce to form buds from which the new animal develops with no or only little contribution from recycled somatic cells.

## RESULTS

### Cell aggregates regenerate a fully functional individual through distinctive stages

To uncover the mechanisms involved in whole body regeneration in *Hydractinia*, we dissociated 12 feeding and 12 sexual polyps into a cell suspension (see methods). The cells were pelleted by centrifugation at 1300 g for 30 minutes, leading to an amorphous cell aggregate composed of randomly arranged cell types. Aggregates were placed in small glass Petri dishes in artificial seawater at room temperature under gentle agitation. Similar to a previous report ^14^, we found that aggregates could regenerate to fully functional feeding polyps within 5 to 9 days. These *de novo* polyps were morphologically and functionally normal (**Figure 1A**). Once fed, they grew stolonal tissue from which new polyps budded. Sexual competence was reached within 2 months (**Figure S1**), showing that regeneration from reaggregated cells can generate a fully functional animal.

**Figure 1.**
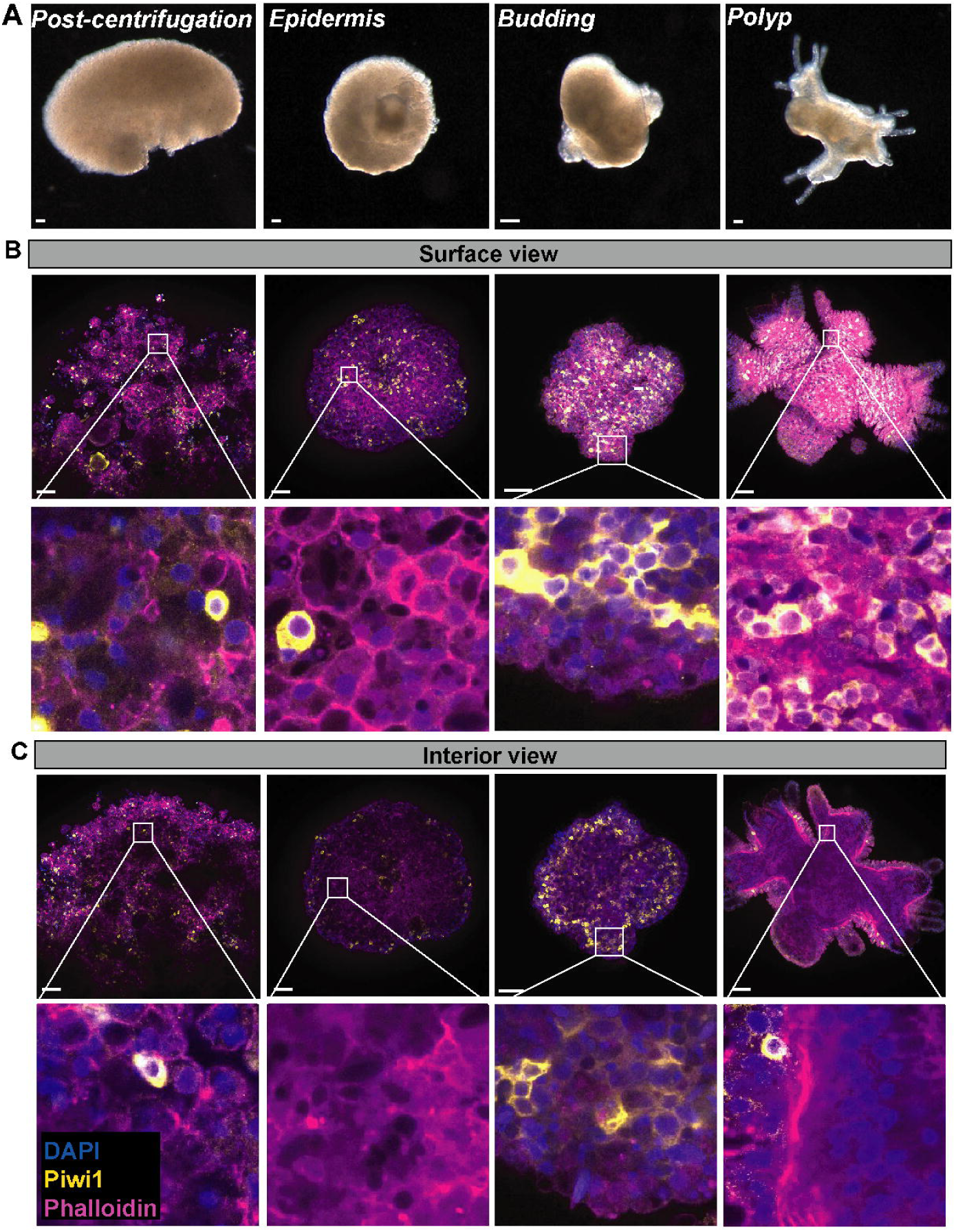
Morphological and cellular regeneration stages of *Hydractinia* aggregates. (**A**) Brightfield images of *Hydractinia* aggregates at different stages of regeneration. (**B**) Maximum projection of aggregate surface, stained with phalloidin (cell membranes), Piwi1 (i-cells), and DAPI (nuclei). (**C**) Aggregate interior. White rectangles correspond to close-up views. Scale bar 100 μm.

While the time and morphological details of regeneration varied between aggregates, regeneration followed a pattern of successive, consistent stages. In the first stage, termed *post-centrifugation* stage, aggregates appeared devoid of structured tissues, forming an amorphic shape with an irregular surface (**Figure 1A**). Within 24 hours, the aggregates became spherical and acquired a smooth outer layer, resembling a normal epidermis; we termed this the *epidermis* stage (**Figure 1A**). Within 2-3 days, one to several small buds emerged from the spherical aggregates; this stage was termed *budding* stage (**Figure 1A**). Finally, heads and tentacles developed at the buds’ distal end, marking the complete regeneration of a polyp that was able to feed; this stage was termed *polyp* stage (**Figure 1A**).

### The cellular events accompanying regeneration

To study the cellular events accompanying the regeneration stages, we stained aggregates with DAPI, phalloidin, FITC-coupled DSA lectin, and antibodies against Piwi1, RFamide, GLWamide, and NCol3 to visualize cell nuclei, filamentous actin, nematocyst capsules, i-cells, neurons, and nematoblasts, respectively. At the *post-centrifugation* stage, phalloidin staining showed no recognizable structure. The distribution of i-cells, nematoblasts, and nematocyst capsules appeared random as expected (**Figure 1B** and **Figure S2**), but no intact neurons were seen.

By the *epidermis* stage, an organized epidermal epithelial layer formed around the aggregates. i-cells appeared in the interstitial space between epithelial cells (**Figure 1B,C** and **Figure S3**), which is their normal localization in intact animals ^16^. However, the interior of the aggregates remained unstructured. Moreover, in the *epidermis* stage, internal cells appeared to be dead, being either devoid of, or having fragmented, nuclei. Piwi1^+^ i-cells were absent in the internal mass, either because they had migrated to the forming epidermis or died. To track i-cells during the first hours following aggregation, we generated cell aggregates from a *Piwi1::GFP* reporter animal ^13^. This animal contains high levels of GFP in i-cells and early germ cells. *In vivo* time-lapse imaging of aggregates from the reporter animal showed migration of i-cells towards the forming epidermis (**Movie S1**). Hence, the absence of i-cells in the internal mass of aggregates was, at least in part, due to their active migration. BrdU incorporation assays showed that no S-phase cells were present during this stage (**Figure S4**). This was unexpected given that i-cells are known to be continuously cycling. Bulk RNA-seq followed by KEGG analysis of differentially expressed genes between aggregate stages (**Figure S5** and **File S1**) revealed that genes implicated in phagocytosis and biological material recycling were upregulated in the *post-centrifugation* and *epidermis* stages, consistent with numerous dead cells being cleared and their content recycled during the first steps of regeneration.

The *budding* stage was characterized morphologically by the appearance of buds at the aggregate surface. Concomitant with the morphological bud development, i-cells coalesced into buds (**Figure 1B,C** and **Figure S3**). It was the earliest stage where S-phase cells were present in aggregates (**Figure S4**). KEGG analysis of differentially expressed genes between aggregate stages confirmed that genes implicated in cell cycling and DNA replication were downregulated until the *budding* stage and started to be upregulated from the *budding* to *polyp* stages (**Figure S5**).

Finally, at the *polyp* stage, heads and tentacles developed. At the cellular level, this stage marked the appearance of a well-established gastrodermal tissue, with a visible mesoglea layer, which is the cnidarian extracellular matrix that divides the two body layers—epidermis and gastrodermis (**Figure 1B,C**). The latter finding is consistent with a recent study, showing that mesoglea components are primarily produced by gastrodermal cells in the sea anemone *Nematostella vectensis* ^17^.

### i-cells are required for aggregate regeneration into polyps

Since i-cells migrated to budding areas, we sought to probe their necessity for whole body regeneration. To this end, we compared the ability of aggregates made from upper body parts of feeding polyps (which contain few or no i-cells ^16^) to regenerate, compared to those sourced from lower body parts (which contain i-cells). We found that aggregates made from lower body parts were able to regenerate normally. However, aggregates made of the upper part of polyps, reached the *epidermis* stage but could not progress to the *budding* stage (**Figure 2A**). These results indicated that i-cells are required for bud formation and subsequent whole-body regeneration but not for the establishment of the initial aggregate epidermis.

**Figure 2.**
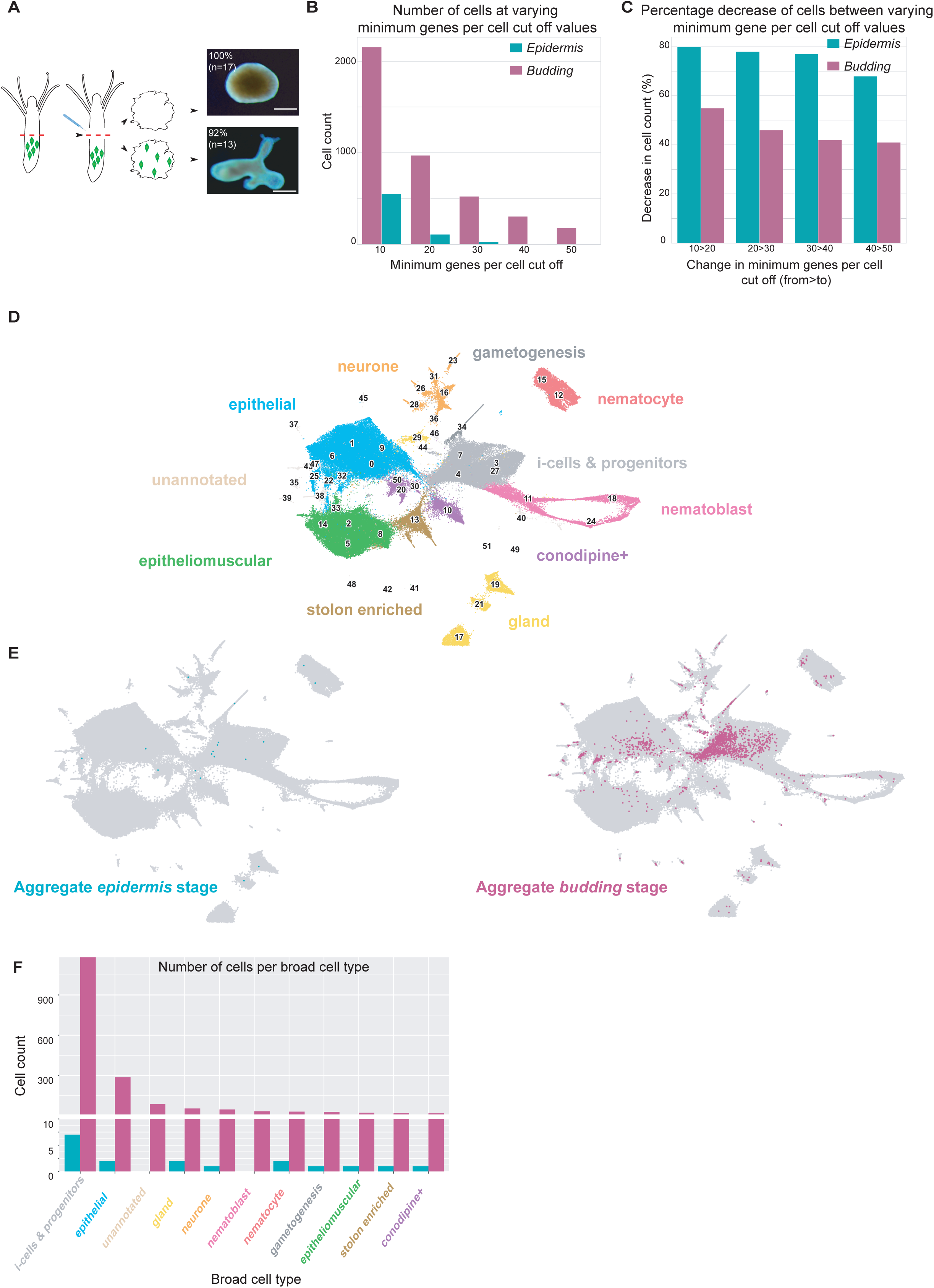
Aggregate regeneration relies mostly on i-cell differentiation. **(A)** Cartoon of the experimental procedure, showing i-cells in green. Brightfield images of *Hydractinia* aggregates made from upper (devoid of i-cells) or lower part (containing i-cell) of feeding polyps showing an absence of complete regeneration when i-cells are not initially present. Scale bar 300 μm (**B**) Total number of cells at varying minimum Genes Per Cell (GPC) thresholds in *epidermis* (teal) and *budding* stage (magenta) aggregate single cell samples. **(C)** Percentage decrease in cells between minimum GPC thresholds in *epidermis* (teal) and *budding* stage (magenta) aggregate single cell samples. **(D)** UMAP of control *Hydractinia* cell atlas integrated with *epidermis* and *budding stage* single cell samples, colored by broad cell type annotation. **(E)** UMAPs split by *epidermis* (teal) and *budding* stage (magenta) single cell samples. **(F)** Number of cells per broad cell type annotation from *epidermis* (teal) and *budding* stage (magenta) single cell samples.

### Aggregate regeneration relies mostly on i-cell differentiation

To gain a better understanding of the cellular composition of aggregates at different stages of regeneration, we coupled the morphological description with single cell transcriptomic analysis. We employed ACME SPLiT-Seq, a robust method for simultaneous fixation and dissociation of tissue, followed by combinatorial indexing for single cell RNA sequencing analysis ^18^. Cell suspensions were prepared from aggregates at the *epidermis* and *budding* stages and processed by SPLiT Seq with a starting input of 120,000 cells per stage. Following barcoding, cells were sorted by FACS to retain only intact, nucleated, single cells, and processed for deep Illumina sequencing. We mapped ∼76 million reads from two sub-libraries of 25,000 cells each to the chromosome level genome assembly of *Hydractinia symbiolongicarpus* ^19^. We assigned reads to cells based on barcode and identified cells belonging to each aggregate stage based on differential Round 1 barcoding. A minimum genes per cell (GPC) cut off was set to 50 GPC in accordance with the *Hydractinia* cell atlas dataset ^20^. Based on 50 GPC, we retained 18 and 1813 cells from *epidermis* and *budding* stages, respectively, showing large cell loss. Typical SPLiT Seq libraries retain 50,000-100,000 cells during post FACS barcoding, and a library of 25,000 post FACS cells usually recapitulates ∼15,000 cells at a 50 GPC threshold after deep sequencing ^18,20–22^. To understand the cell loss observed in the *epidermis* stage, we assessed the total number of cells retained at varying GPC thresholds (10, 20, 30 40 and 50 GPC). With decreasing GPC, we retained more cells from both aggregate stages (Figure 2B), with *budding* stage aggregates showing less loss than *epidermis* stage ones. To further examine the differential cell loss between *epidermis* and *budding* stage aggregates when the GPC threshold was increased, we looked at the percentage loss of cells between threshold values (**Figure 2C**). At every GPC cut-off change, *epidermis* stage aggregate cells were lost with a higher percentage than *budding* stage cells. Therefore, the mRNA level in most *epidermis*-stage cells is either low or none. This is consistent with the observations made by microscopy (**Figure 1B,C**), showing that the interior of the *epidermis*-stage aggregate is made up mostly of dead cells.

To determine the contribution of different cell types to the regenerating aggregate, we merged the pre-processed, 50 GPC cut-off aggregate sub-libraries with the *Hydractinia* cell atlas dataset. After clustering, we transferred broad cell type annotations from the atlas dataset to the new, combined dataset (**Figure 2D**) and confirmed the annotations by plotting the expression of known marker genes (Figure S6). The overall distribution of aggregate cells across the combined dataset (**Figure 2E,F**) showed that in the *epidermis* stage, most cells present were i-cells and progenitors (**Figure 2E,F**), and to a lower extent, epithelial cells. By the *budding* stage, most cell types re-emerged.

Overall, the above analyses reveal three key findings: first, at the *epidermis* stage, most cells are dead. Second, those that are alive are mostly i-cells and early progeny. Lastly, cell type diversity is restored in the *budding* stage. Therefore, the fully regenerated animal derives primarily from i-cells rather than from recycled somatic cells of the dissociated polyps.

### Cell proliferation is not required for bud formation

Since aggregated cells only re-entered the cell cycle during the *budding* stage (Figure S4, S5), we asked whether buds could form from i-cells directly inherited from the dissociated polyps, without generating new ones via self-renewal. For this, we inhibited cell proliferation using hydroxyurea (HU). We confirmed the action of HU using decapitated polyps in which the contribution of cell proliferation to head regeneration is established ^13^. Control decapitated polyps were able to regrow a head within 3 days but as expected, HU prevented regeneration of treated decapitated polyps (**Figure S7**). No BrdU was incorporated in HU-treated polyps as opposed to control ones (Figure S7). We found that inhibiting cell proliferation in aggregates by HU treatment had no effect on bud formation. Aggregates treated with HU were indistinguishable from control aggregates at the *epidermis* and early *budding* stages, showing that the initial two stages of aggregate regeneration are conducted by cells directly inherited from the dissociated polyps (**Figure S8**). The majority of these cells are i-cells and early progenitors (**Figure 2**). However, HU-treated aggregates were not able to give rise to a fully functional polyp. They remained arrested at the *budding* stage, unable to develop a head or tentacles to complete regeneration (Figure S8). Investigating the i-cell population by Piwi1 immunostaining revealed that only few i-cells were present in *budding*-stage aggregates treated with HU compared to the control (**Figure S8**). The reduction in i-cell numbers while buds developed is consistent with i-cell differentiation into various cell types, allowing the aggregates to generate buds. However, i-cells’ inability to self-renew under HU prevented complete polyp regeneration. Therefore, initial budding is independent of proliferation.

### Bud formation depends on i-cell migration

To shed more light on i-cell migration during aggregates regeneration, we prevented cell migration by using cytochalasin B (CB), a drug known to prevent actin polymerization, thereby inhibiting any cell movement including cell rearrangement ^23^. First, we verified the action of the drug in decapitated polyps in which the contribution of cell migration for head regeneration is established ^13^. In CB-treated decapitated polyps, we found that the wound was unable to close, demonstrating that this drug strongly inhibits all cell movements. Piwi1 immunostaining showed that in control polyps, Piwi1^+^ cells migrated to the injury site within 24 hours post decapitation, as expected ^13^. In CB-treated polyps, Piwi1+ cells did not migrate toward the injury site, staying at the lower part of the body column (**Figure S9**). Treating regenerating aggregates with CB post-centrifugation led to their death without forming an epidermis (**Figure S10**). These results suggested that aggregate epithelization is achieved by morphallactic cell rearrangement, as also found in the related hydrozoan *Hydra* ^11,24^. To prevent i-cell migration post epidermis formation, we treated aggregates at the *epidermis* stage. This treatment prevented aggregates from budding (Figure S10). From these results we hypothesized that an absence of cell migration prevents i-cell clustering, and therefore bud formation, in aggregate and polyp regeneration.

### Sphingosine signaling controls i-cell migration

To identify candidate regulators of i-cell migration in *Hydractinia* regeneration, we compared gene expression between successive stages of aggregate regeneration by bulk RNA sequencing (**Figure S5** and **File S1**). We found that genes of the sphingosine pathway were upregulated during early aggregate regeneration (**File S1** and **Figure S5**). The sphingosine pathway is involved in development ^25^ and many cellular processes, including cell proliferation ^26^, differentiation ^27^, and migration and motility ^28–30^. The pathway is activated by the binding of sphingosine-1-phosphate (S1P)−produced by the phosphorylation of sphingosine by sphingosine kinase (Sphk)−to its receptors (S1PR) in an autocrine or paracrine fashion. The *Hydractinia* genome encodes one Sphk homolog and 8 S1PRs (**Figure S11**). Therefore, we chose to investigate the role of the sphingosine pathway during regeneration. First, we investigated the expression of sphingosine pathway components in aggregates by *in situ* mRNA hybridization chain reaction (HCR). *Sphk* was expressed broadly throughout the entire aggregate at *epidermis* and *budding* stages (Figure 3A,B). *S1pr1* (one of the 8 S1PR receptor homologues) was expressed by i-cells (studied in intact and decapitated polyps; Figure 3C,D), suggesting that these cells can respond to S1P.

**Figure 3:**
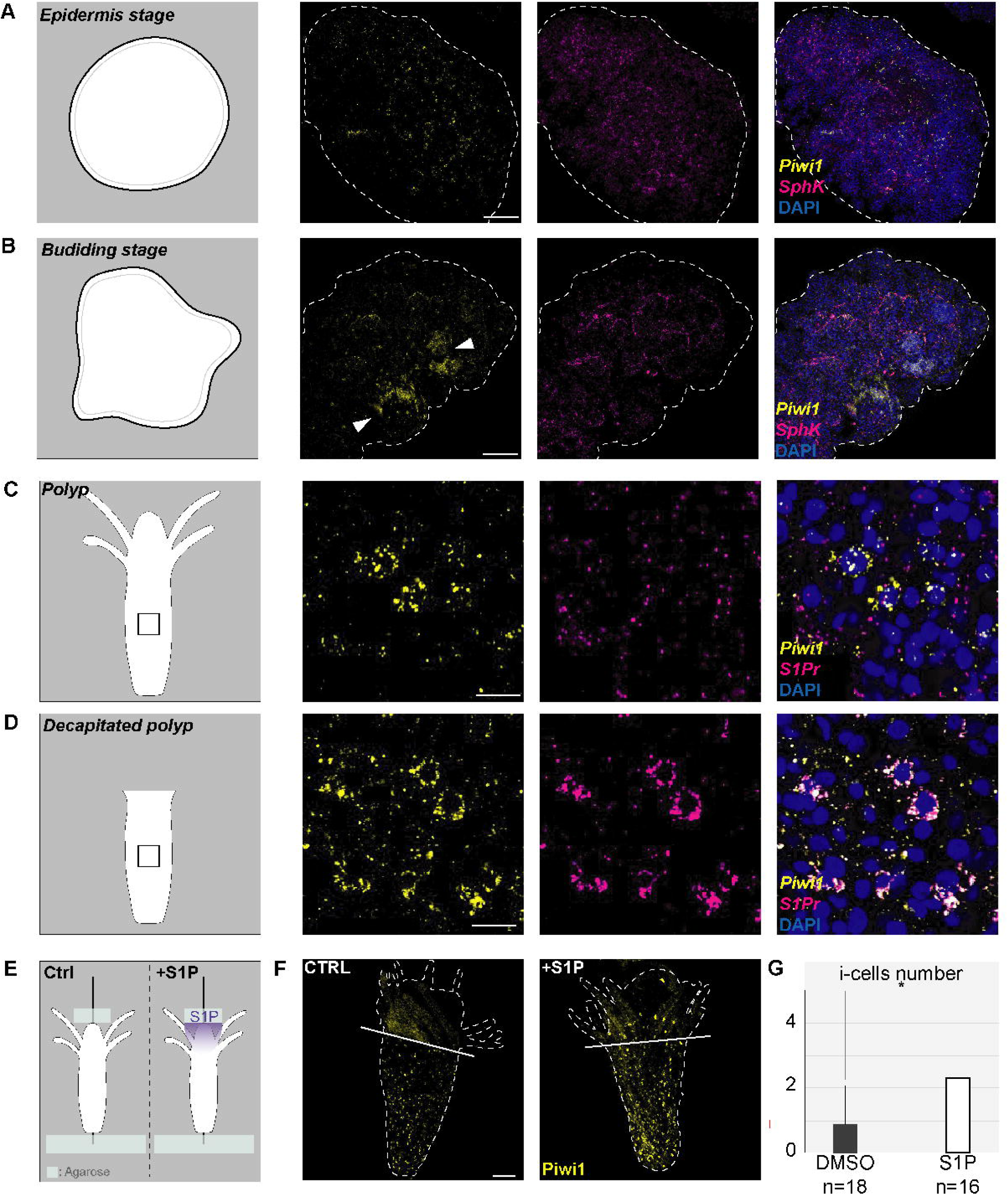
S1P pathway in regeneration. (**A-B**) Cartoon of aggregates and confocal section of mRNA HCR *in situ* hybridization of *Sphk* and *Piwi* at *epidermis* (**A**) and *budding* stage (**B**). (**C-D**) Cartoon of a *Hydractinia* polyp and confocal section of mRNA HCR *in situ* hybridization of *S1Pr* and *Piwi1* in control (**C**) and in a decapitated polyp (**D**). Black rectangles represent the position of the close-up view of the HCR *in situ* hybridization. (**E**) Experimental procedure to test i-cell attraction to S1P during regeneration. A block of agarose containing DMSO (control) or 50 μM S1P were attached to the polyp at the oral pole. (**F**) Piwi1 immunostaining indicates an increase of i-cells located in the head of S1P-treated polyp, compared to control. (**G**) The differences in the number of i-cell in the head or tentacles of polyps treated, or not, with 50 μM of S1P is statistically significant. White bar represents the position of the lowest tentacle. White dashed lines show the contour of aggregates/polyps. Scale bar 100 μm (A, B, E) and 10 μm (C, D).

To test if an S1P gradient can directly induce i-cell migration in *Hydractinia*, we performed a migration assay on live polyps. For this, we attached blocks of agarose containing 0, 50 or 500 μM S1P to the oral tip of polyps using a metal pin (**Figure 3E**). In normal polyps, most i-cells reside in the lower body column, below the lower tentacle line ^13^. We assessed the number of i-cells present in the head of polyps (i.e., above the upper tentacles) exposed to the two different concentrations of S1P and compared this to control polyps to which an agarose block with no S1P was attached. We found that the number of i-cells was significantly higher in heads of polyps exposed to 50 μM S1P compared to control (≃2,5 vs ≃0,8), (**Figure 3F,G**). In polyps exposed to 500 μM S1P, the number of i-cells present in the head was similar to the control (≃1,4 vs ≃1), (**Figure S12**). This is consistent with i-cells migrating along a S1P gradient, stopping once they have reached an area of high concentration ^28^.

To evaluate the function of the sphingosine pathway during the regeneration process, we treated decapitated polyps with FTY720, a structural analogue of sphingosine S1P which blocks the S1PRs ^31^, and with SKI-II, a Sphk inhibitor ^32^; both drugs have been demonstrated to interfere with sphingosine signaling ^29^. In decapitated polyps treated with SKI-II or FTY720, wound healing was normal, but the polyps were unable to regenerate a head (Figure S7). Importantly, both SKI-II and FTY720 did not block proliferation, indicating that the phenotype is not a side effect of cell cycle arrest (Figure S7). Anti-Piwi1 antibody staining showed that i-cells could not migrate under treatment, remaining predominately in the lower body column at one day post decapitation while i-cells accumulated in the upper blastema in the control (Figure S9). These results suggest that the sphingosine pathway regulates i-cell migration in polyps. We then performed the same inhibition experiments in aggregates, exposing them to SKI-II or FTY720. Drug treated aggregates reached the *epidermis* stage, but no further regeneration was observed (**Figure 4A,B**). The pattern of i-cell distribution in treated and control aggregates was assessed following anti-Piwi1 IF using coordinate variance, which is a measure of sample dispersion. The higher the variance the greater is the spread of the cells (see Materials & Methods). Piwi1 immunostaining showed that in SKI-II-or FTY720-treated aggregates, i-cells remained spread around the epidermal tissue and no clustering occurred, consistent with a block of active migration (**Figure 4C,D**). Lack of i-cell migration was confirmed by time-lapse imaging of aggregates made from the Piwi1::GFP reporter animal, treated with SKI-II inhibitor (**Figure S13** and **Movie S2**).

**Figure 4.**
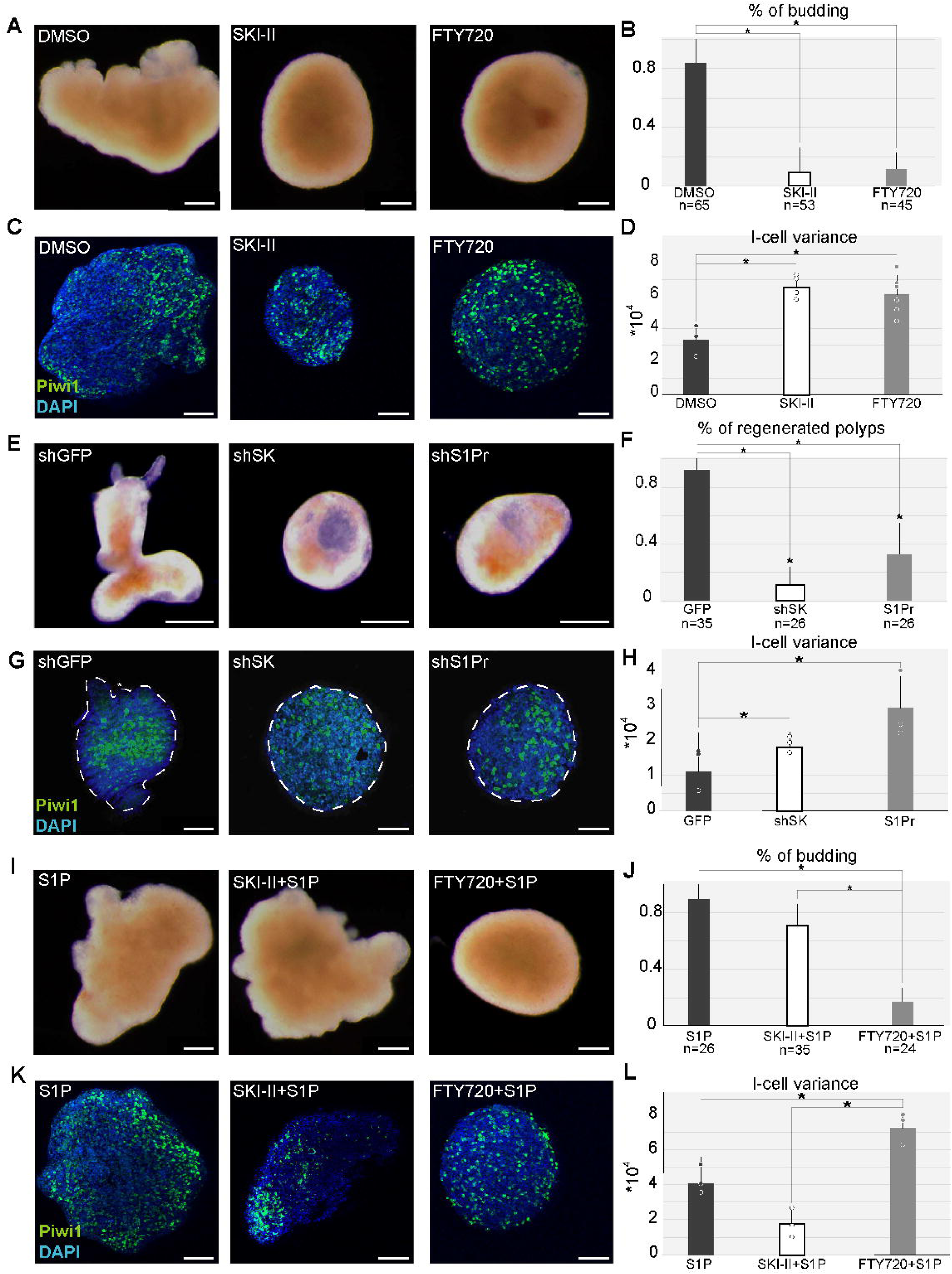
The S1P pathway is required for aggregate regeneration. (**A**) Brightfield images of control (DMSO) *budding* stage aggregates and ones treated with Sphk inhibitor (SKI-II) or S1pr inhibitor (FTY720). (**B**) Proportion of aggregates that reach *budding* stage, showing that both SKI-II and FTY720 prevent bud development. (**C**) Piwi1 immunostaining showing i-cell location in control *budding* stage aggregates and ones treated with SKI-II or FTY720. (**D**) Quantification of i-cell variance as a measure of cell dispersion (see methods section) per condition. The lower variance in control indicates i-cell clustering, opposed to treated aggregates in which the variance remains high. (**E**) Brightfield images of control aggregates electroporated with shRNA targeting *GFP* (shGFP) and aggregates electroporated with shRNA targeting *Sphk* (shSK) or *S1pr* (shS1P). shSK and shS1P disrupted aggregates are unable to reach the *budding* stage, conversely to control aggregates which fully regenerate to the *polyp* stage (**F**). Proportion of aggregates that reach *polyp* stage. *Sphk* and *S1Pr* loss of functions prevent bud development. (**G**) Piwi1 immunostaining, showing i-cell location in aggregates at *polyp* stage which were electroporated with shGFP, shSK and shS1Pr. (**H**) Quantification of the i-cell variance per condition. Significantly lower variance in control indicates i-cell grouping into clusters, opposed to shSK and shS1P electroporated aggregates. (**I**) Brightfield images of aggregates incubated with S1P, S1P plus SK, and S1P plus FTY720. (**J**) Proportion of aggregates that reach *budding* stage per condition. S1P treatment does not affect the budding. Addition of S1P to SKII-treated aggregates rescues the phenotype as buds developed normally, while FTY720 effect remains unchanged even with S1P addition. (**K**) Piwi1 immunostaining showing i-cell location in aggregates incubated with S1P, S1P and SK, S1P and FTY720. (**L**) Quantification of i-cell variance per condition. In the SKI-II+S1P condition, a decrease in variance indicates i-cell clustering conversely to FTY720+S1P treatment. White dashed lines show the contours of the aggregates. Scale bar 100 μm.

To validate the drug results genetically, we knocked down the *Hydractinia Sphk* and *S1pr1* genes, which were upregulated during regeneration in our RNA-seq experiments (**Figure S5** and **File S1**). For this, we designed short hairpin RNAs (shRNAs) against *Sphk* and *S1pr1* to *S1pr8* (**Table S1**) mRNA which we delivered to the dissociated cell suspension via electroporation, prior to reaggregation. A shRNA against *GFP* served as control. Control aggregates whose cells were electroporated with a shRNA targeting *GFP* regenerated a functional polyp with head and tentacles. However, aggregates generated from cells electroporated with shRNA targeting either *Sphk* or *S1pr1* were not able to regenerate a polyp (**Figure 4E,F**). No phenotype was observed when shRNA against *S1pr2*-*8* was electroporated. The efficiency of *Sphk* and *S1pr1* shRNAs was assessed by qPCR on decapitated polyps (**Table S2** and **Figure S14**). i-cells remained sporadically distributed in knockdown aggregates, while their distribution was clustered in control ones, electroporated with *GFP* shRNA (**Figure 4G,H**). The latter were able to regenerate an intact polyp.

We performed rescue experiments by adding S1P to the seawater in which aggregates were kept. As SKI-II inhibits the production of S1P by blocking Sphk, we hypothesized that S1P addition to SKI-II-treated aggregates should rescue the phenotype. Conversely, addition of S1P to FTY720-treated aggregates should not since this drug blocks the receptor. As expected, S1P alone had no effect on aggregate regeneration; however, S1P addition to SKI-II-treated aggregates rescued i-cell clustering and budding. Aggregates exposed both to FTY720 and S1P did not generate buds, and their i-cells remained spread throughout the aggregate (**Figure 4I-L**).

Taken together, regeneration of polyps from reaggregated cells requires S1P signaling-dependent i-cell migration, and these cells are the main contributors to the new individual.

## DISCUSSION

*Hydractinia* aggregate regeneration proceeds through several, well-defined stages, starting with the formation of an epithelial epidermal layer around a mass of dead cells; the latter may function physically to provide mechanical support and/or as a source of nutrients to fuel regeneration, as supported by upregulation of endocytosis genes in our KEGG analysis (**Figure S5**). The epidermis is colonized by migratory i-cells within 24 hours. These i-cells then migrate laterally, guided by S1P. Eventually, i-cells form buds that develop into normal polyps (**Figure 5**). Our data also show that the regenerated individual predominately derives from differentiating proliferative i-cells, rather than from recycled somatic cells. Epithelial cells that survive dissociation and aggregation form the initial epidermis but only marginally contribute to the eventual polyp. Also of interest, i-cells re-enter the cell cycle only during the *budding* stage. The reasons for this transient withdrawal from the cell cycle remain unknown.

**Figure 5.**
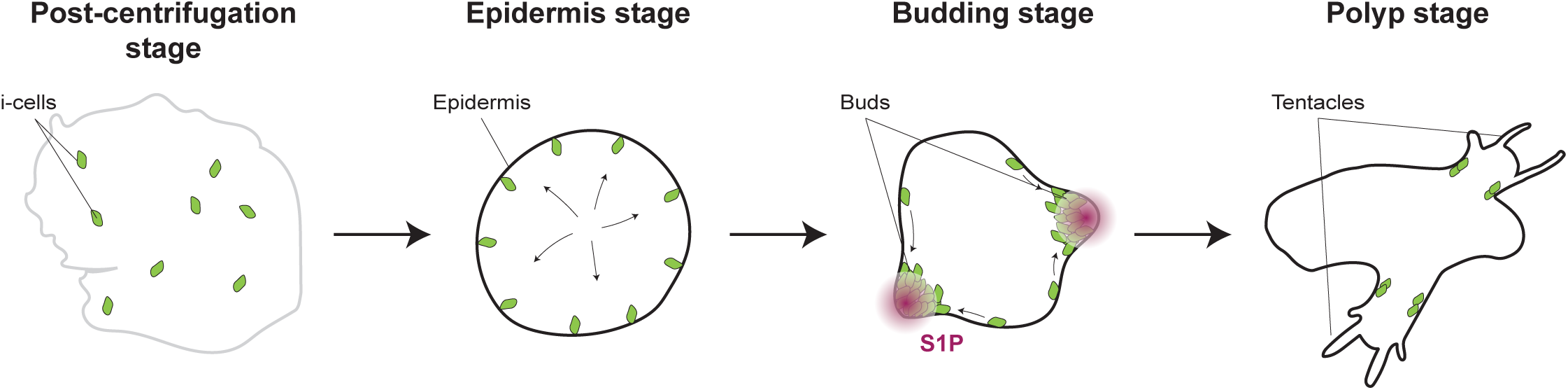
A model summarizing the major events in aggregate regeneration.

While we directly show that i-cells migrate along an S1P gradient (**Figure 4I-L**), the expression pattern of *Sphk*, not being restricted to foci, makes it difficult to explain the localized source of S1P. Furthermore, our rescue experiments were successful even when S1P was ubiquitously added to seawater, further complicating the understanding of how the gradient is formed. One possibility is that S1P phosphatase and/or lyase regionally lower S1P concentrations, resulting in the formation of a S1P gradient as shown to be the case in the tunicate *Botryllus schlisseri* ^28^.

Although the ability to regenerate a functional adult from reaggregated cells is widespread in hydrozoans ^7^, the freshwater, solitary polyp, *Hydra*, regenerates through a different mechanism that includes the spatial sorting and reuse of somatic cells. The fate and function of *Hydra* i-cells—as well as of mitosis—in the process are unclear. The difference in regeneration mode between these two hydrozoans could be explained by the distinct developmental potential of i-cells in *Hydra* vs *Hydractinia*. In *Hydra*, i-cells are not pluripotent. The animal’s body is composed of three segregated and developmentally restricted cell lineage ^33^. Therefore, representatives of all three lineages need to be reused in regeneration. *Hydractinia* i-cells, by contrast, are pluripotent, contributing to all somatic lineages and to germ cells ^14^, leaving recycled somatic cells dispensable for regeneration.

Regeneration from disorganized reaggregated cells is unlikely to be adaptive since it does not occur naturally. It is therefore interesting to speculate on how this capability had evolved and whether it shares features with embryonic development. In this context, while most studied animals (including many cnidarians) develop stereotypically, *Hydractinia* embryogenesis appears rather erratic (**Movie S3**) ^34^. An external epithelial layer is established by the 16/32-cell stage, enclosing a mass of mesenchymal cells but the process overall is variable ^34^, similar to aggregate regeneration. First 1-4 Piwi1^+^ i-cells mostly appear in the internal mesenchymal cell mass around the 32/64-cell stage (**Figure S15**). They migrate outwards to the epidermal layer during metamorphosis to generate the primary polyp ^35^, resembling i-cell migration during the establishment of the *epithelial* and *budding* stages of aggregate regeneration. The relative simplicity of *Hydractinia*’s morphology does not require a highly ordered, stepwise construction of structures, permitting considerable lenience in morphogenesis; this is also true for aggregate regeneration.

In summary, cell aggregate regeneration in *Hydractinia* resembles embryogenesis in several aspects including the initial formation of an epithelial layer and the central role i-cells play in the process. We propose that the ability of aggregates to regenerate is a byproduct of the animal’s ‘liberal’ mode of development.

## ACKNOWLEDGEMENTS

We thank members of the Frank lab for discussions and advice. Cian Lawless is kindly acknowledged for animal care. Confocal microscopy and flow cytometry were conducted at the Centre for Microscopy and Imaging, and at the Flow Cytometry Core Facilities at University of Galway, respectively. U.F. is a Wellcome Trust Investigator in Science (grant no. 210722/Z/18/Z). G.K. was an Irish Research Council post-doctoral fellow (project ID GOIPD/2020/149). H.R.H. was a doctoral student in the Science Foundation Ireland Centre for Research Training in Genomic Data Science (grant no. 18/CRT/6214).

## AUTHOR CONTRIBUTION

CC, GK, and UF conceptualized the study. CC and GK performed experiments. CC performed in situ hybridization and S1P migration assay. GK performed time-lapse imaging. LR performed experiments on cell proliferation. AV performed experiment on sexual maturity and immunofluorescence on embryos. CC and HRH generated and collected samples for single-cell transcriptomic experiments. HRH performed bioinformatic single-cell analyses. GK performed phylogeny. CC, GK, HRH and UF wrote the paper. All authors read and approved the final version of the manuscript.

## MATERIALS AND METHODS

### Hydractinia husbandry

Adult *Hydractinia symbiolongicarpus* colonies were grown on glass slides kept in artificial seawater (ASW) at room temperature. Animals were fed four times per week with *Artemia franciscana* nauplii, and once a week with pureed oysters. To induce scheduled spawning, we kept the animals in a constant 14:10 light:dark cycle, where females and males spawn 1.5 hours after exposure to light.

### Aggregates

An aggregate consists of 12 feeding and 12 sexual polyps dissociated in 200 µL of ASW. Polyps were cut and incubated for 12 minutes in Calcium and Magnesium Free Seawater (CMFSW) (for 1L: 26.24g of NaCl, 0.671g of KCl, 4.687g of Na2SO4, 4.3 mL of NAHCO_3_, 10mL of Tris-HCl 1M pH 8.0, 10mL of EGTA 0.25M pH 8.0) to facilitate their mechanical dissociation. CMFSW was then discarded and replace by ASW. Polyps were mechanically dissociated into cell suspension by drawing them in and out of an 18G needle affixed to a 1mL plastic syringe. The dissociated cell solution was filtered using a pluriStrainer Mini 100 µm filter (43-10100-40; PluriSelect). The filtered solution was centrifuged at 2000g for 30 minutes at 18°C in a swinging bucket rotor. Using a Paster pipette the cell pellets were placed into ASW containing 5 U/mL of Penicillin-Streptomycin antibiotics (P0781, Merck) in medium glass Petri dishes. Aggregates were maintained at 18°C on rocker and ASW was changed every day. 24 hours post-centrifugation, dead cells surrounding the aggregates were removed by pipetting up and down with a Paster pipette.

### Gene identifications and phylogenetic analysis

We performed Blastx searches against the genome of *Hydractinia symbiolongicarpus* ^19^ using the amino-acid sequences of human sphingosine kinase and the five sphingosine 1-phosphate receptors as queries, followed by a reciprocal Blast search. We aligned all amino-acid sequences from the open reading frame using MAFFT 7 software ^36^ and removed vacancies and blur sites with Gblocks 0.91b ^37^. Phylogenetic analysis was performed with MrBayes 3.1.2 ^38^ under mixed model. The amino acid evolution model used was Blossum for both sphingosine 1-phosphate receptors and sphingosine kinase analyses. The analyses were ran for 500 000 generations with 10 randomly started simultaneous Markov chains (1 cold, 9 heated). One fourth of the topologies were discarded (burn-in values), and the remaining were used to calculate the posterior probability representing each nodes robustness.

### Pharmacological treatments

Sphingosine kinase inhibitor SKI-II (ab141594; Abcam), sphingosine receptor antagonist FTY720 (SML0700; Merck), and Cytochalasin-B (250233; Merck) were suspended in DMSO at −20°C and used at the final concentration of 5 µM, 1 µM, and 6 µM, respectively. Hydroxyurea (H8627; Merck) was suspended in H_2_O at −20°C and used at a final concentration of 5 mM. Sphingosine 1-Phosphate (SML2709; Merck) was used at a final concentration of 5 µM. Chemicals were administrated at the *post-centrifugation* stage, excepted for the cytochalasin-B added at the *post-centrifugation* and the *epidermis* stage.

### S1P migration assay in living polyps

*H. symbiolongicarpus* colonies were starved for 3 days and anesthetized in 4% MgCl_2_ (in 50% distilled water/50% filtered seawater). Polyps were dissected from the colony. Feeding polyps were skewered with a Minutien pin (26002-10; Fine Science 315 Tools). Then, a block of 0.5% agarose gel containing either 50 µM or 500µM S1P, or DMSO (control), was pressed against the head of the polyps and left for 2 hours for the chemical to diffuse into the tissue. After 2 hours, polyps were detached and kept in sea water overnight before being fixed in in 4% formaldehyde in PBS.

### HCR *in situ* hybridization

*S1pr*, *Sphk* (HCR amplifier: B1-647) and *Piwi1* probe sets (amplifier: B3-488) were generated by Molecular Instruments Inc. Polyps and aggregates were fixed in 4% PFA overnight for 1 hour at RT. Polyps were washed 3 times with 1× PBS then dehydrated in 100% methanol. Samples were stored at –20°C overnight. Samples were rehydrated in a series of MeOH/PBSTween 0.1% dilutions (75% MeOH:25% 1× PBSTw, 50% MeOH:50% 1× PBSTw, 25% MeOH:75% 1× PBSTw, 100% 1×PBSTw). Samples were incubated in PBST 0.1% for 10 minutes (6 times), followed by 3 washes of 5 minutes in 5XSCCTw. Samples were then pre-hybridized in probe hybridization buffer (Probe hybridization buffer, Molecular Instruments Inc) for at least 4 hours at 37°C. Samples were then incubated with 3 pmol of the probe set diluted in probe hybridization buffer for 16 hr at 37°C. Samples were washed 3 times for 20 minutes each with probe wash buffer (HCR probe wash buffer, Molecular Instruments Inc) at 37°C. Samples were washed 2 times for 5 minutes with 5× SSCTw and incubated with probe amplification buffer (HCR, Molecular Instruments Inc) containing snap cooled amplifier hairpins (1: 50) at room temperature. Samples were then washed 4 times in 5xSSCtw for 30 minutes.

### Statistical analyses

Statistical significances were evaluated by Wilcoxon Mann-Whitney test using the software R 4.3.1. Effects were considered significant with a p-value < 0.05 and the notation of the p-value is as follows: *p<0.05.

### Imaging and Image analysis

Images were taken with an Olympus FV3000 confocal microscope. Time-lapse images were taken with a Leica MZ16FA fluorescence stereomicroscope and Leica DFC350FX camera. Samples for time lapse were mounted on microscope slides in 0.5% low melting agarose (BP16525; Thermo Fisher Scientific) sealed using the coverslip sealant Biotium 23005. Image analysis and processing were done using FIJI. Their brightness and contrast were adjusted as a whole. We aimed to display all polyp images in the same orientation, therefore some images were rotated, and a black rectangle was inserted behind for aesthetic reasons. i-cells distribution was evaluated using FIJI. Images were first corrected for binarized using a manual color Threshold. Once segmented the ultimate point coordinates were obtained using the FIJI cell counting tool. Then, either the number of i-cells or their coordinates were extracted. The coordinates were then exported in Excel where their variance was calculated.

### Immunostaining

Animals were first relaxed in 4% MgCl_2_ for 15 minutes and fixed in 4% formaldehyde in PBS overnight at 4°C. Animals were washed three times in PBS - 0.5% Triton (PBST) and blocked for 1 hour in 3% BSA/PBST. Primary antibodies (anti-NCol (1:500) ^39^, anti-Rfamide (1:1000) ^40^, anti-GLW (1:1000) ^40^ and anti-Piwi1 (1:2000) ^41^in BSA/PBST and incubated overnight at 4°C, followed by three washes with PBST then blocked for 30 minutes in 5% serum in BSA/PBST. Secondary antibodies (Alexa Fluor 488 (Abcam, # A32731) goat anti-rabbit IgG, Alexa Fluor 594 (Abcam, # A-11012), 647 (Abcam, # A-21245) were diluted 1:500 in BSA/PBST/serum and incubated for 1 hour at room temperature. Animals were washed three times with PBST, incubated in 1:2000 Hoescht 33258 (20 mg/ml, Sigma-Aldrich # B2883), DAPI (1mg/ml, ThermoFisher #62248) (1:2000) or preconjugated Alexa Fluor 546 - Phalloidin (1:2000, ThermoFisher; #A12381) in PBST and washed a further three times in PBST. Animals were mounted in 97% 2,2’-Thiodiethanol (TDE).

### BrdU

BrdU immunodetection was performed on aggregates after incubation in ASW containing 150μM of BrdU overnight. Post incubation aggregates were fixed, treated with 2M HCL for 30 minutes and the immunostaining protocol was carried as stated above using a primary antibody against BrdU (Abcam; #ab6326; 1:100).

### Short-hairpin RNA synthesis and electroporation

Short-hairpin RNAs (shRNA) were designed as previously described ^42^. Synthesis carried out for three days using the HiScribe™ T7 High Yield RNA Synthesis Kit (E2040S; New England Biolabs). The synthesis mix was treated for 1 hour at RT with DNase I and purified using the Monarch® RNA Clean-up Kit (T2050L; New England Biolabs) according to manufacturer protocol. Cell suspension for electroporation was obtained by dissociation of 40 polyps from colonies of male cloned 291-10 (20 feeding and 20 sexual polyps) in 20 µL, per aggregate. Electroporation mix was composed of 12 µL of 1.54M D-mannitol suspended in H_2_O (240184, Merck), 18 µL of dissociated cells, and 18 µL of shRNA solution. Final concentration of shRNA was 2000µg/ µL. Electroporation was performed in Cuvette Plus™ Electroporation Cuvettes (7321136; BTX) at 25V with a single 25 msec pulse using a homemade electroporator. shRNA used are provided in Table S1.

### Real-time PCR

Aggregates were dissolved in TRIzol (15596026; Invitrogen) and RNA was extracted as described ^42^. cDNA was synthetized using the Omniscript RT Kit (205111; Qiagen) according to manufacturer protocol. Real-time PCR was conducted on a StepOne Plus machine with in fast run mode under Quantification – Comparative Ct (ΔΔCt) experiment and using the TaqMan system (4444556; Applied Biosystems). Thermal cycler was run at: 95 °C for 15s for the holding stage; then 40 cycles of amplification at 95 °C for 1s and 64 °C for 30s with 10µL of reaction mix. Reactions were done in triplicate and GAPDH was used as reference gene as previously described ^43^. Oligos used are provided in Table S2.

### RNA libraries preparation, sequencing, and sequences analysis

RNA libraries were prepared by sampling 8 aggregates per replicate. Samples consist of three replicates taken at 5 time points corresponding to *post-centrifugation*, *epidermis*, *budding* (one at the beginning, one at the end), and *polyp* stages. Aggregates were dissolved in TRIzol (15596026; Invitrogen) and stored at −80°C. RNA was extracted from TRIzol solution as described ^44^. The eluted RNA was quantified using NanoDrop and shipped to Novogene Europe (Cambridge, UK) for further processing and sequencing. RNA integrity was assessed using the RNA Nano 6000 Assay Kit of the Bioanalyzer 2100 system (Agilent Technologies, CA, USA), and the library preparations were sequenced on an Illumina Novaseq platform and 150 bp paired-end reads were generated. An index of the reference genome was built using Hisat2 v2.0.5 and paired-end clean reads were aligned to the reference genome using Hisat2 v2.0.5. The mapped reads of each sample were assembled by StringTie (v1.3.3b). Pairwise differential expression analysis between aggregate stages (three biological replicates per stage) was performed using the DESeq2 R package. The resulting P-values were adjusted using the Benjamini and Hochberg’s Approach, and genes with an adjusted P-value <=0.05 found by DESeq2 were assigned as differentially expressed.

### ACME SPLiT Seq

Cells from 480 *epidermis stage* aggregates and 480 *budding stage* aggregates were collected by simultaneous fixation and dissociation, as previously described ^20^. Aliquots of resulting suspensions were diluted 1 in 3 in PBS and stained with 5 uM Draq5 to identify nucleated cells and 38 nM ConcanavalinA-488 to assess the presence of cytoplasm. Cells were analysed by flow cytometry, using an AcurriC6+ Sampler, to calculate the number of single (FSC-A vs. FSC-H), nucleated (Draq5+) and intact (ConcanavalinA-488+) cells per microlitre of the original suspension. Cells were loaded 5000 cells per well, based on the previous flow cytometric analysis, into Round 1 of barcoding for SPLiT Seq, with half of the Round 1 plate (1-6) containing *epidermis stage* cells and half (7-12) containing *budding stage* cells, allowing for later identification. SPLiT Seq barcoding, library preparation and sequencing then continued as previously described ^20^.

### Data Analysis

Aggregate SPLiT Seq libraries were pre-processed according to parameters used in ref ^20^, and only cells meeting a minimum Genes Per Cell threshold of 50 were retained. The aggregate libraries were then merged with the *Hydractinia* cell atlas ^20^. Sub-libraries were labelled according to type (Ep = *epidermis stage* aggregate, Bd = *budding stage* aggregate, Control = atlas cells) and the data was integrated using Harmony ^45^. Clustering was performed and broad cell types were identified based on known marker genes and previous cluster annotation from the *Hydractinia* cell atlas. The presence of cell types in each aggregate stage was assessed by analysis of the proportion of cells in each broad cell type. To assess cell loss from the aggregate dataset, pre-processing was performed at varying Genes Per Cell (GPC) cut off thresholds; 10, 20, 30, 40 and 50 GPC. These datasets were labelled according to their GPC cut off threshold and merged. Cells were then labelled based on their aggregate stage and an analysis of cell loss between stages was performed by assessing percentage reduction between threshold values.

---

**Figure S1. A sexually mature aggregate.** Brightfield image of a sexually mature male aggregate. White rectangle shows a close-up view of a sexual polyp. White arrows indicate sporosacs.

**Figure S2: Intact neurons are absent at *post-centrifugation* stage.** Single confocal slices of aggregates stained with FITC-coupled DSA lectin, and antibodies against Ncol3, GLWamide, RFamide and Piwi2. Nematoblasts (Ncol3) and nematocytes (DSA) are widely distributed in the aggregates. Neurons (GLW^+^ and RFamide^+^) are absent or fragmented. Scale bar 100 μm.

**Figure S3: i-cell distribution in aggregates.** Single confocal slices of Piwi1-immunostained i-cells in aggregates made exclusively with feeding polyps, showing i-cells clustering in buds. Scale bar 100 μm.

**Figure S4: Proliferation in *Hydractinia* aggregates. (A)** Single confocal slices of Piwi1 and BrdU immunostaining showing i-cells in S-phase in aggregates at *epidermis*, *budding*, and *polyp* stages. Scale bar 100 μm. (**B**) Magnification of a bud at the *budding* stage. Scale bar 10 μm.

**Figure S5: KEGG analysis of bulk RNA-sequencing comparing each stage of aggregate regeneration.** We sampled *epidermis* stage at two timepoints (onset of bud appearance, end of bud formation). From *post-centrifugation* to *epidermis* stages, actors of the sphingosine pathway are detected to be upregulated. Until *polyp* stage, indicators of cell death and biological material recycling are upregulated (phagosome, lysosome). Cell proliferation markers (DNA replication, cell cycle) are downregulated at both *post-centrifugation* and *epidermis* stages and start to be upregulated from the *budding* stage on.

**Figure S6: Marker gene expression plots.** UMAP expression plots of top marker genes per cluster for the integrated *Hydractinia* cell atlas with the aggregate single cell dataset.

**Figure S7. Effects of proliferation and sphingosine pathway inhibition on regeneration of decapitated polyps.** (**A**) Control decapitated polyps (DMSO) regenerate a full head within 3 days, while those treated with hydroxyurea (HU), cytochalasin-B (CB), SKI-II, and FTY720 do not. **(B)** BrdU labelling at one day post-decapitation. i-cells forming the blastema on the injury site of decapitated control polyp massively proliferate. HU-treated polyps do not proliferate. CB-, SKI-II-, and FTY720-treated polyps have some proliferative cells, but they remain in the lower part of the body column. This indicates that these treatments block i-cell migration but not proliferation. Scale bar 200 μm.

**Figure S8. Inhibition of cell proliferation in aggregates.** (**A**) Brightfield images of control (DMSO) and HU-treated aggregates at *budding* stage. (**B**) Quantification of the number of aggregates that reached *budding* stage per condition. (**C**) Brightfield images of control (DMSO) and HU treated aggregates at *polyp* stage. (**D**) Quantification of the number of aggregates that reached *polyp* stage per condition. (**E**) Piwi1 immunostaining showing i-cell location in control aggregates at *polyp* stage and in aggregates treated with HU. (**F**) Quantification of the number of i-cells per condition. Scale bar 100 μm.

**Figure S9. Inhibition of i-cell migration in decapitated polyps.** (**A**) Piwi1 immunostaining, showing i-cell location in control (DMSO), CB, SKI-II and FTY720-treated polyps 24 hours after decapitation. Images are oriented with the amputation site to the top. (**B**) Cartoon explaining the quantification method. During the first day post-decapitation, i-cells migrate from the lower part of the body to the upper part (black arrow). The number of i-cells in the lower and the upper halfs of the polyps were counted to evaluate the percentage distribution of i-cells along the body column. (**C**) Quantification of the percentage distribution of i-cells. In control regenerating polyps about 70% of the total i-cells are localized in the upper part of the body, reflecting their migration to the wound. Conversely, CB, SKI-II, and FTY720-treated polyps have a significantly lower proportion of i-cells in their upper part, resulting from defects in their migration. Scale bar 100 μm.

**Figure S10. Treating aggregates with cytochalasin-B prevents bud formation.** (**A**) Brightfield images of control (DMSO) and CB-treated aggregates. (**B**) Quantification of the number of aggregates that reach *budding* stage per condition. (**C**) Piwi1 immunostaining, showing i-cell location in control aggregates and aggregates treated with CB. Scale bar 100 μm.

**Figure S11. Phylogenies of the Sphk and S1Pr.** Phylogenies of Sphk and S1Pr made by Bayesian inference on amino acid sequences. Sampling was made on species representative of metazoan diversity. Sphk is present at the metazoan scale, with a single copy in *Hydractinia*. S1Pr phylogeny shows a diversification of *Hydractinia* receptors with 8 paralogues. Node supports represent posterior probabilities.

**Figure S12. S1P migration assay in polyps.** (**A**) Piwi1 immunostaining, showing i-cell location in polyps treated, or not, with 500 μM S1P. (**B**) Quantification of the number of i-cells in the head or tentacles of polyps treated, or not, with 500 μM of S1P. White bar represents the limit between the head and the body column. White dashed lines show the contour of polyps. Scale bar 100 μm.

**Figure S13. Time lapse imaging of i-cell migration.** Evaluation of i-cell migration from the inner toward the outer part of aggregates based on Movies S1 and S2. (**A**) the number of i-cells inside an area defined by two circles (grey area) were counted at 4h, 8h, and 12h post-centrifugation. Percentages of i-cells variance inside the area were calculated from one time point to the next one. (**B**) Localization and counting of i-cells detected in outer part of aggregates from Movies S1 (DMSO) and S2 (SKI-II). (**C**) Evolution of i-cells proportion close to the outer part of aggregates. Number of i-cells close to the epidermis increased by around 30% due to their active migration in the control, while their number remained similar in the SKI-II-treated aggregates.

**Figure S14. Real-time PCR of *S1pr1* and** *Sphk* **genes.** Relative expression of both *S1pr1* (**A**) and *Sphk* (**B**) genes compared to the *Gapdh* gene reference. Expression level significantly decreased in decapitated polyps electroporated with sh***S1pr1*** and sh*Sphk*, respectively, in comparison to the control (shGFP).

**Figure S15. Appearance of embryonic i-cells.** 64-stage *Hydractinia* embryos showing the first i-cells. Grey, DNA. Green, Piwi1.

**Movie S1.** Time lapse of aggregates made from Piwi1::GFP reporter animals showing i-cell migration from the inner toward the outer parts of aggregates between 4-12h post-centrifugation.

**Movie S2.** Time lapse of aggregates from 4-12h post-centrifugation made from Piwi1::GFP reporter animals, treated with SKI-II. i-cell do not migrate from the inner toward the outer parts of aggregates.

**Movie S3.** Time lapse of *Hydractinia* embryogenesis.

