## Supplementary figures and images for "An unexpected mode of whole-body regeneration from reaggregated cell suspension in *Hydractinia* (Cnidaria, Hydrozoa)"

### Figure S1

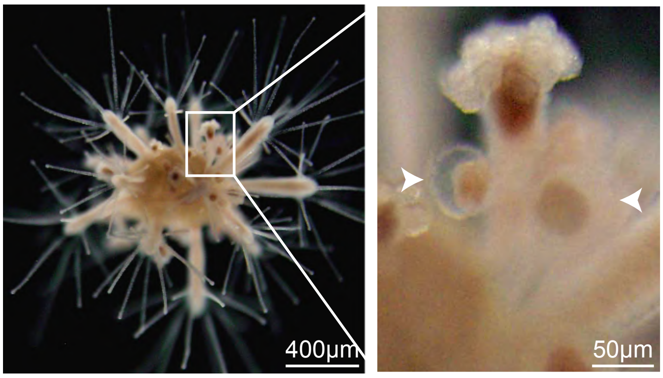

### Figure S2

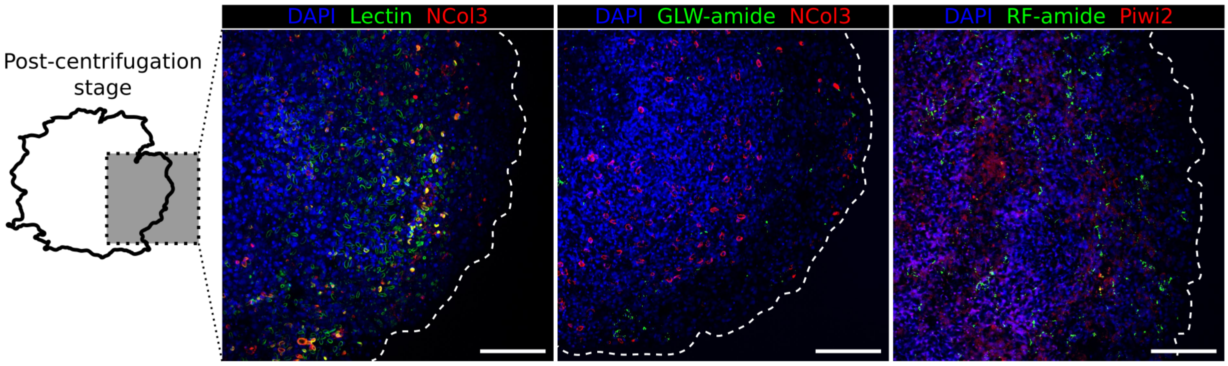

### Figure S3

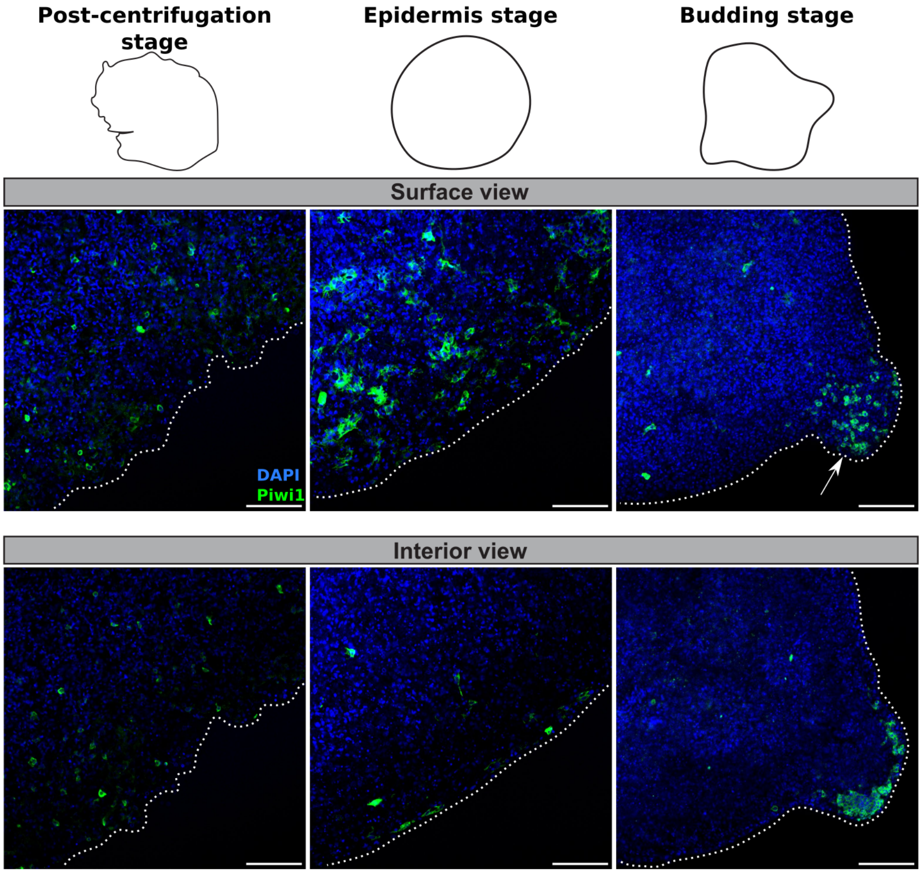

### Figure S4

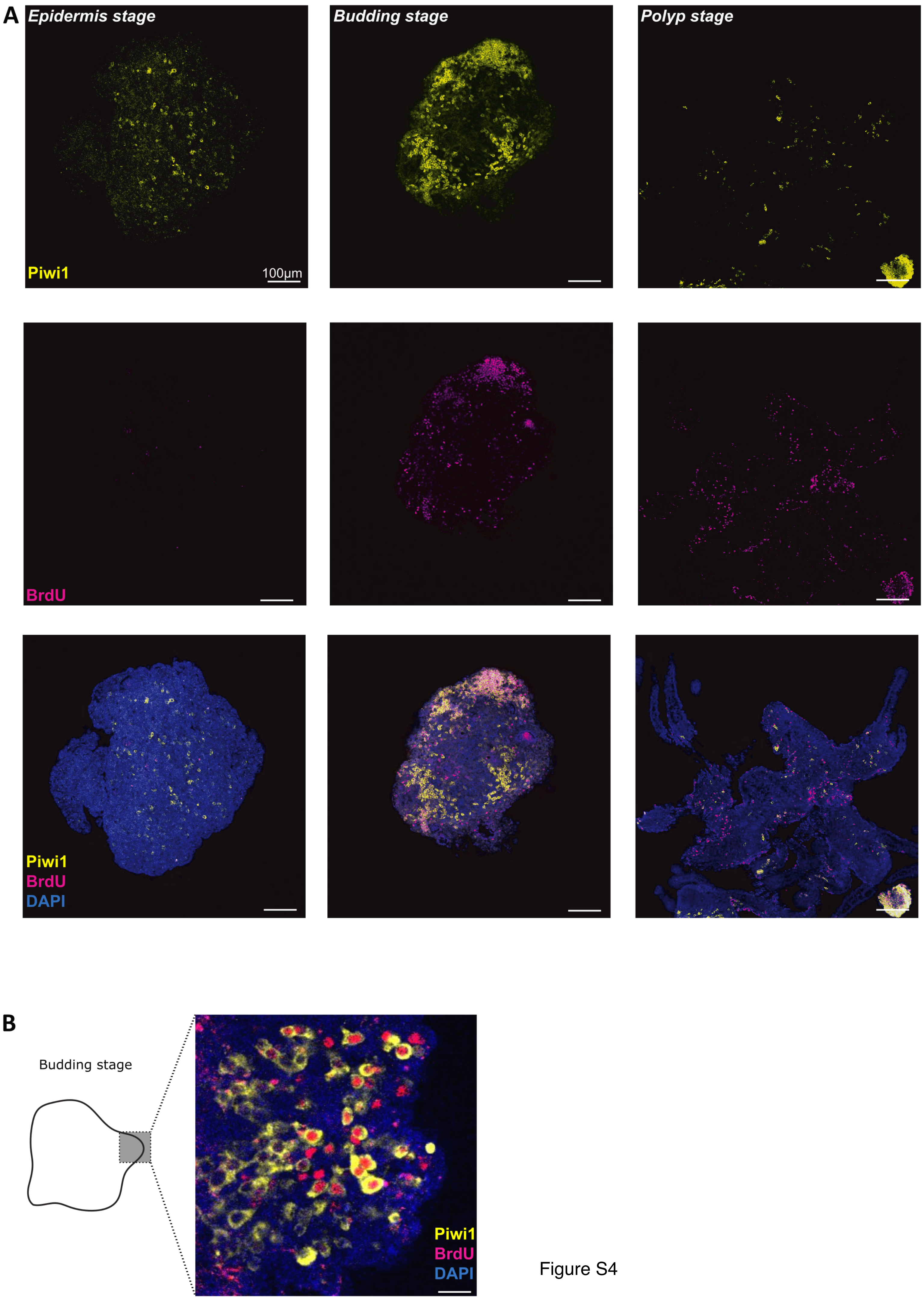

### Figure S6

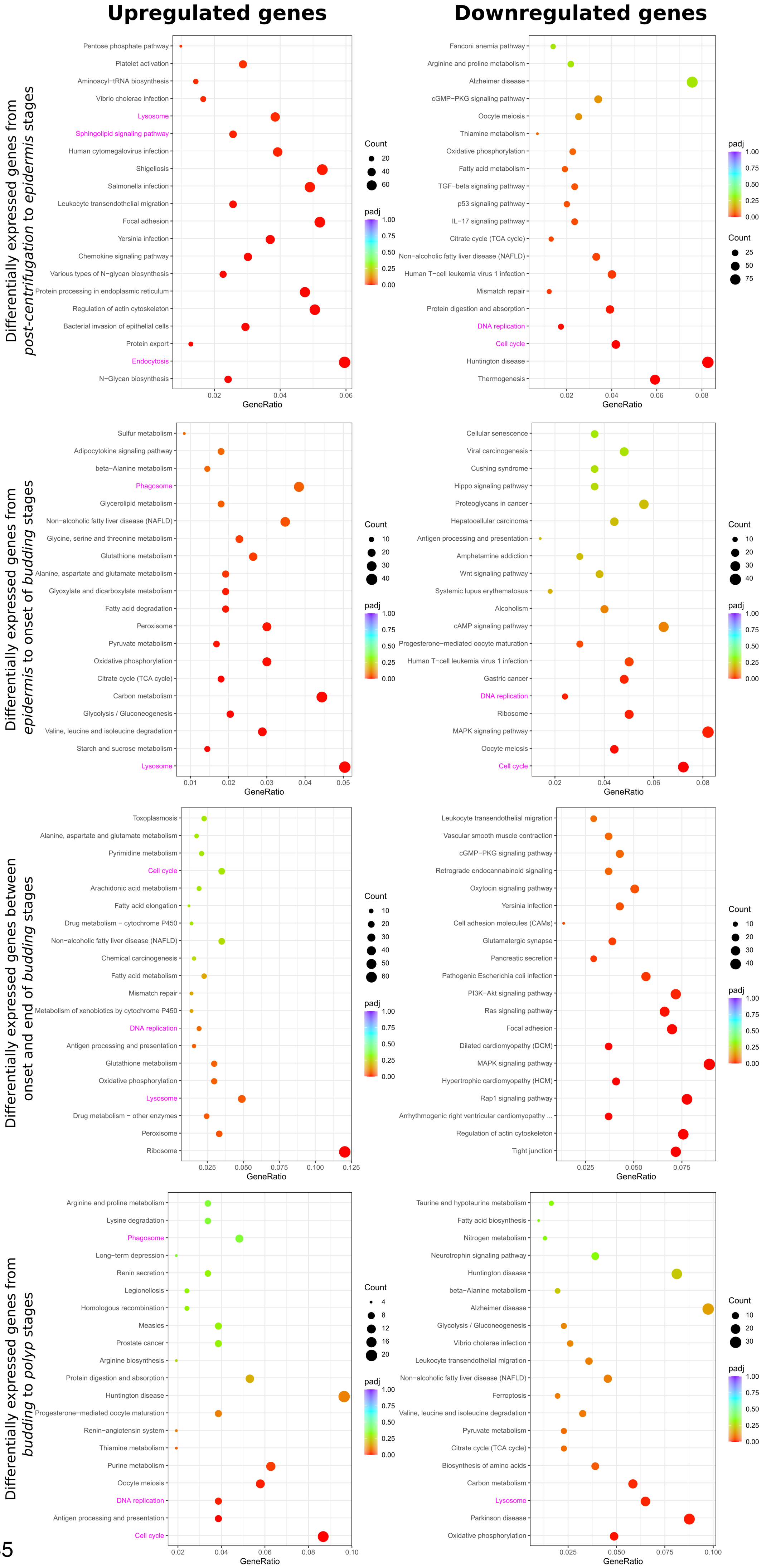

### Figure S7

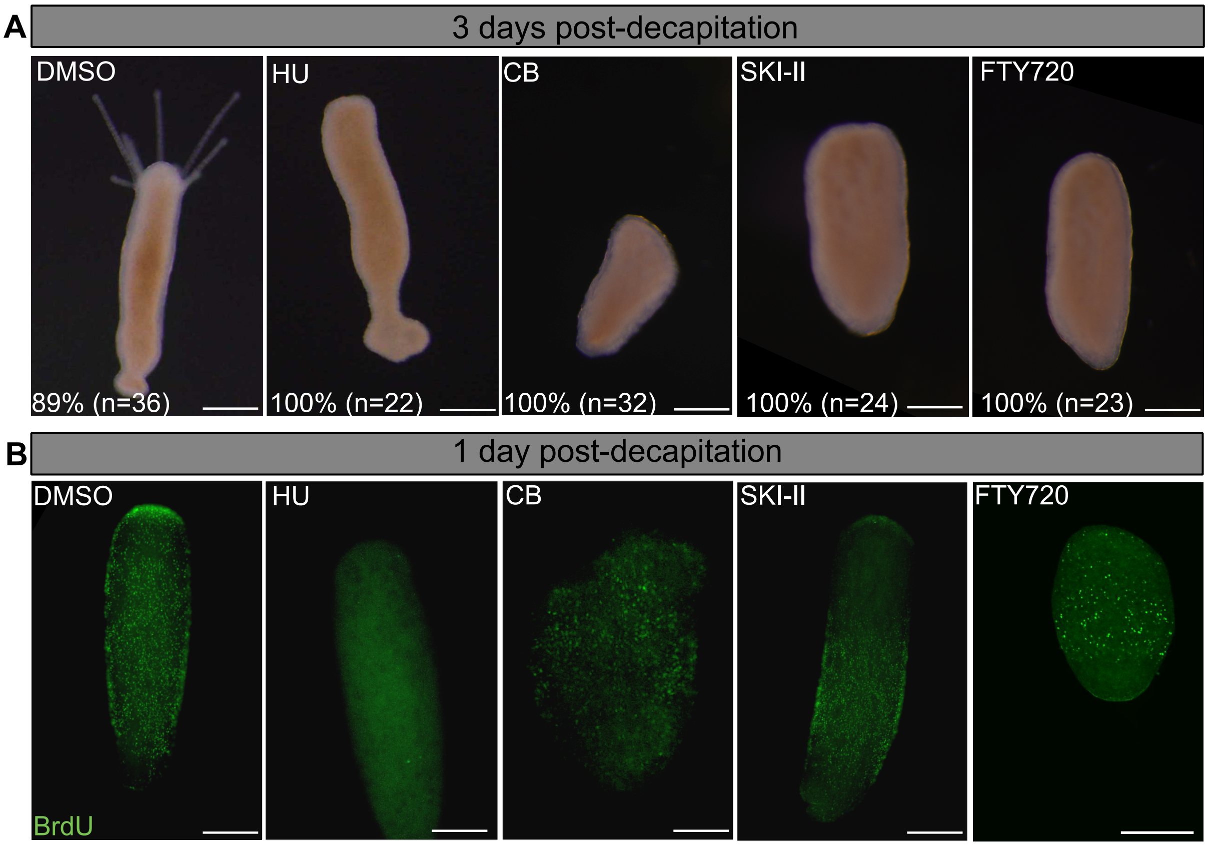

### Figure S8

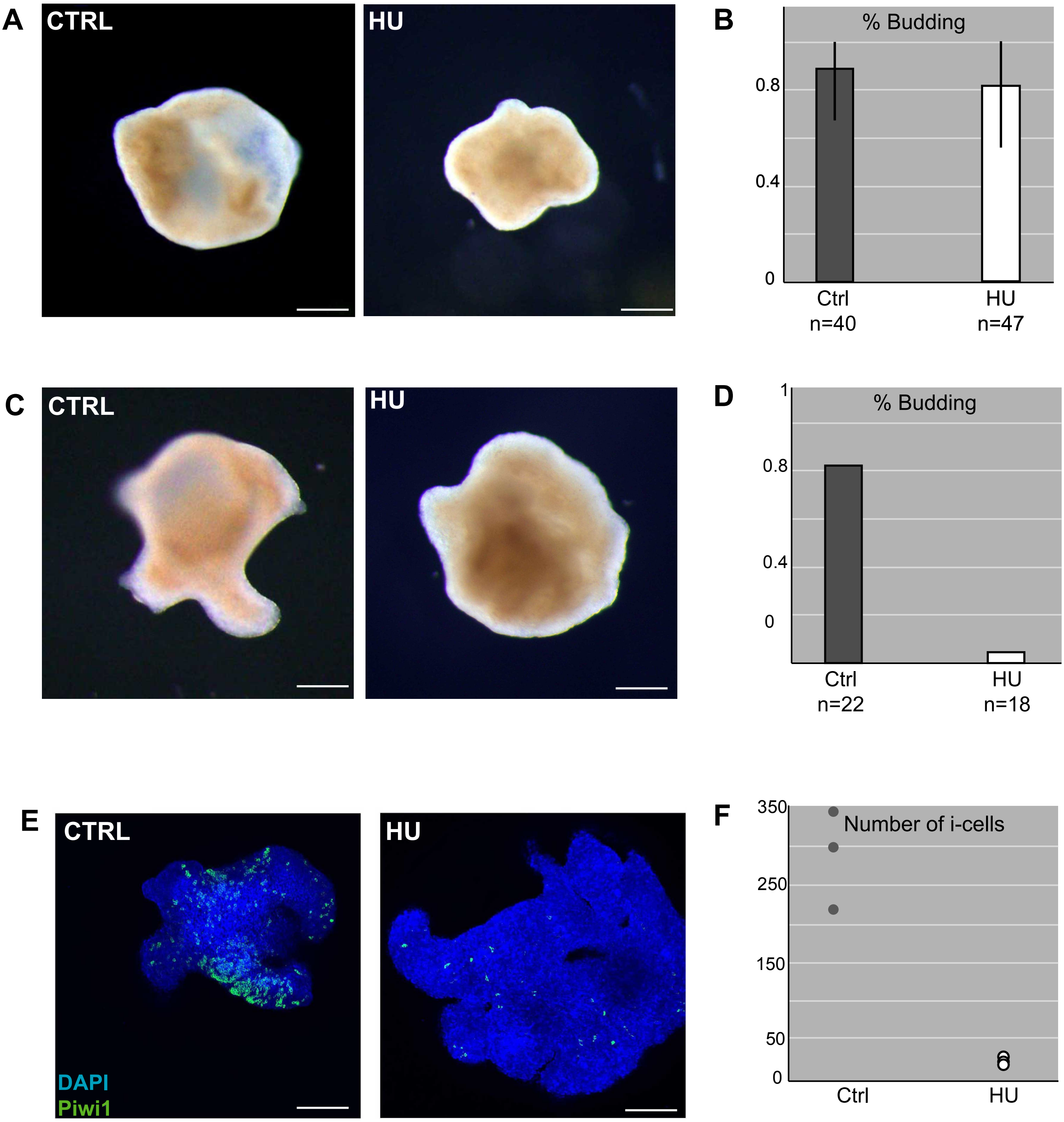

### Figure S9

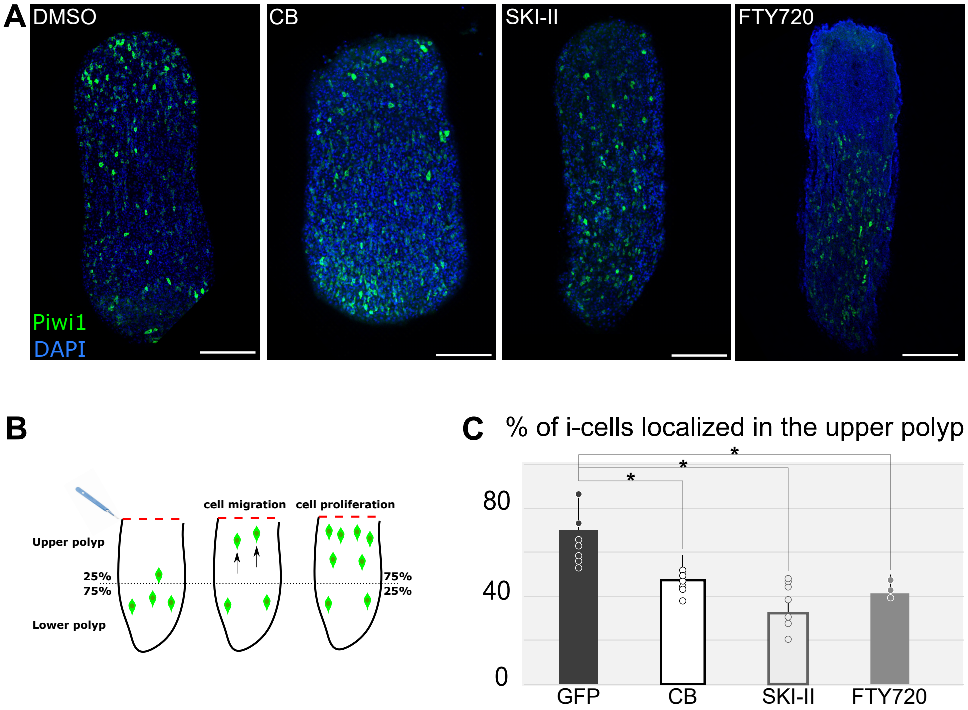

### Figure S10

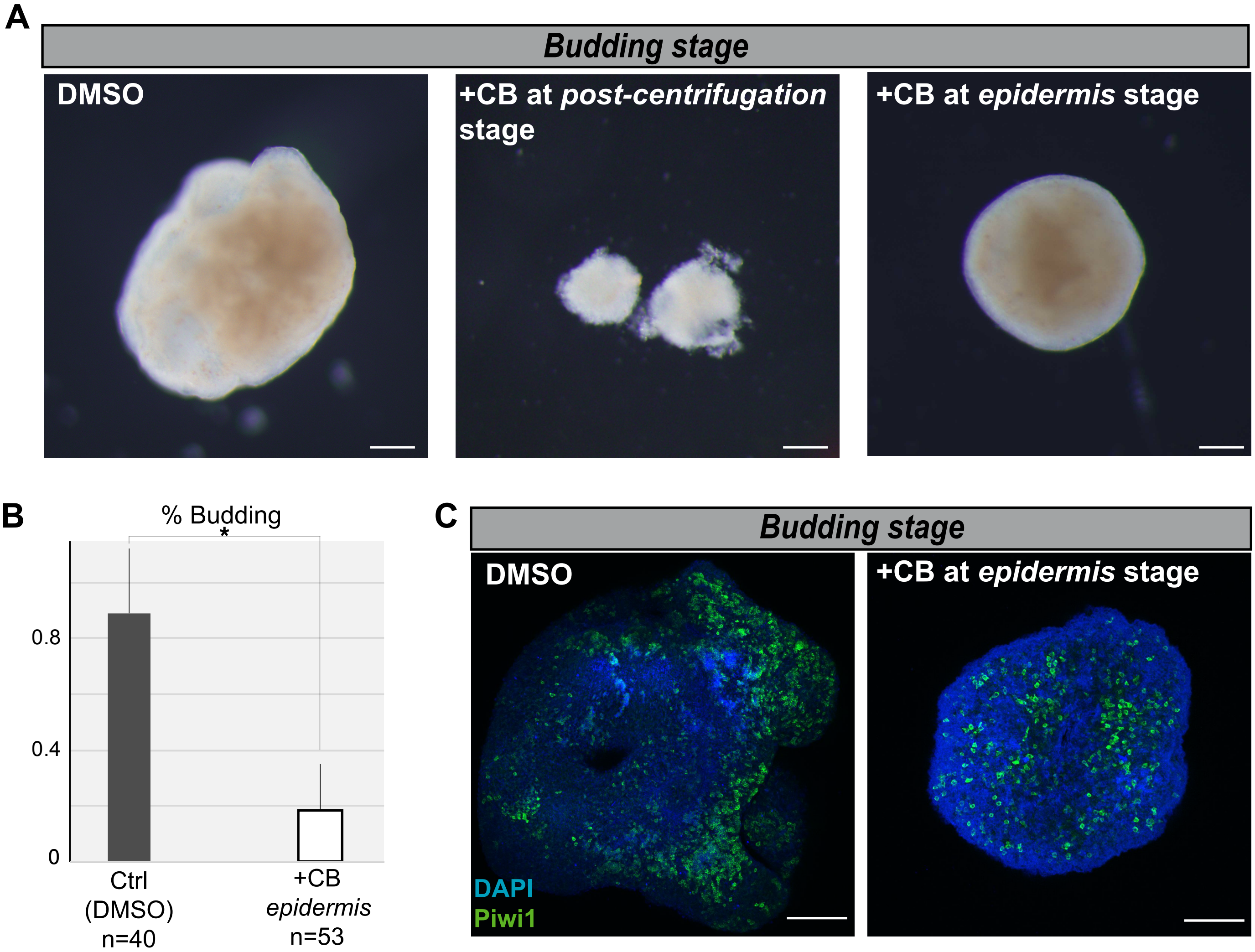

### Figure S11

#### Sphingosine kinase phylogeny

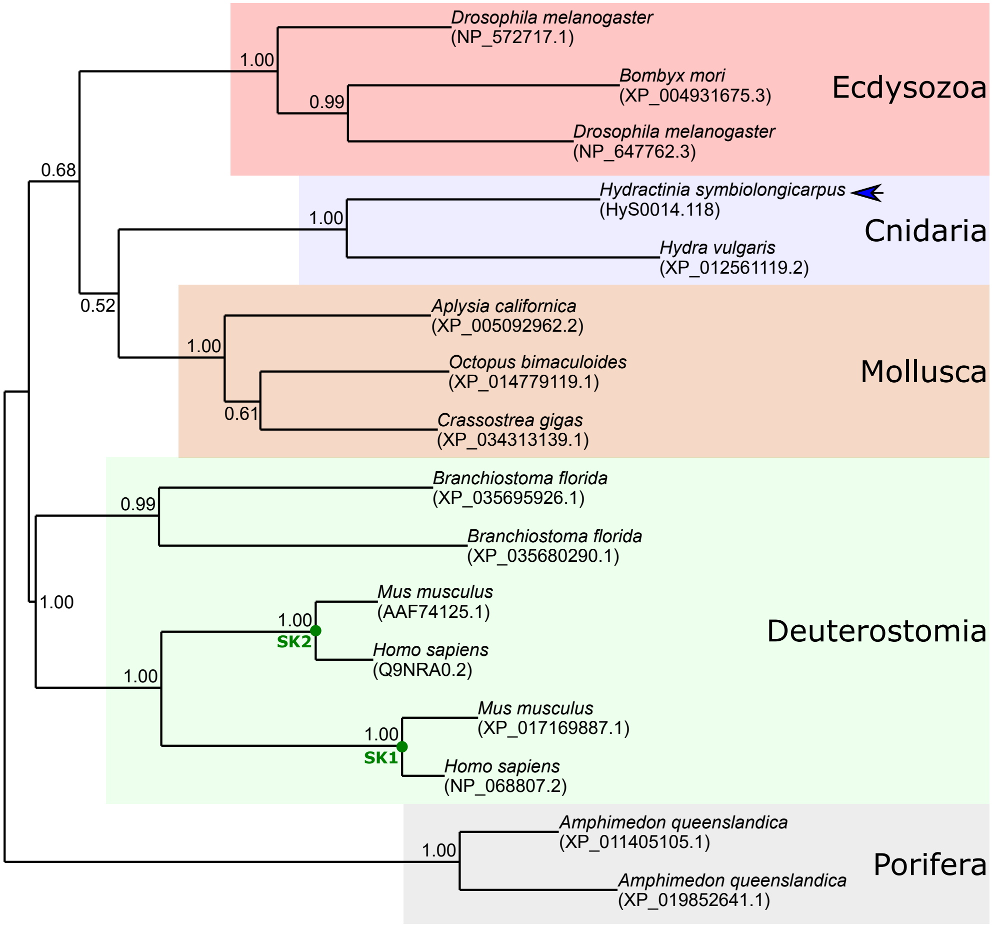


#### S1P receptors phylogeny

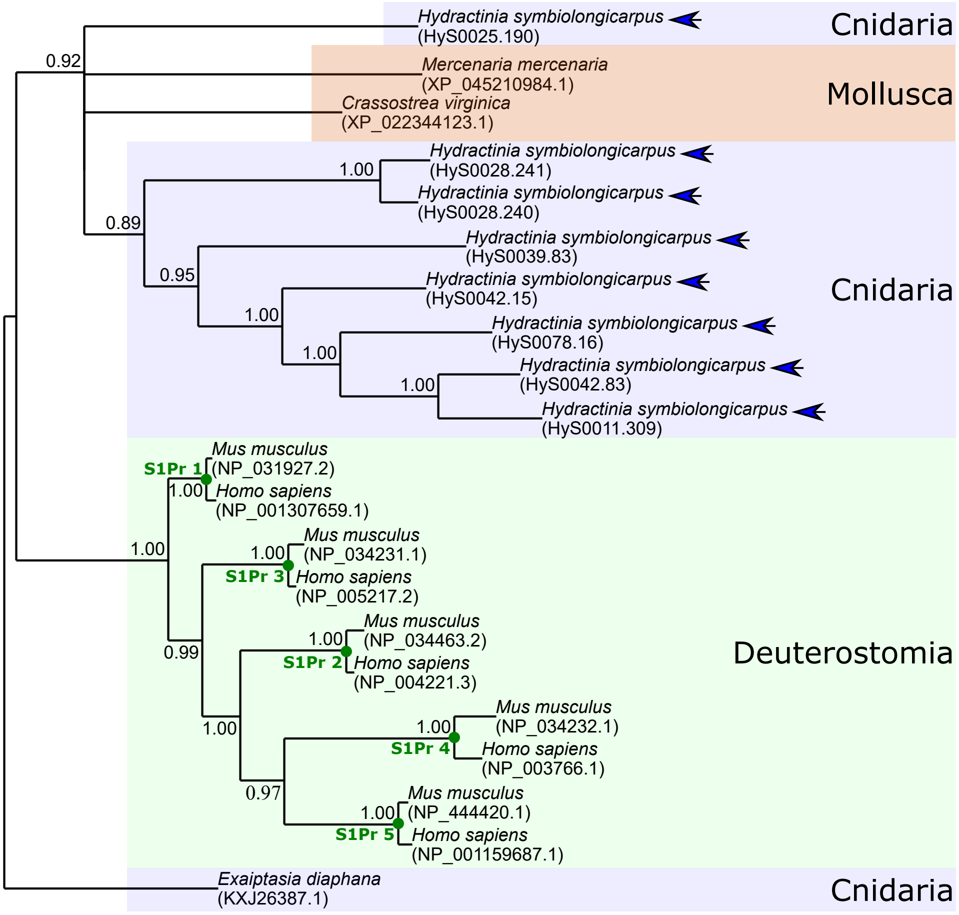


Figure S11

### Figure S12

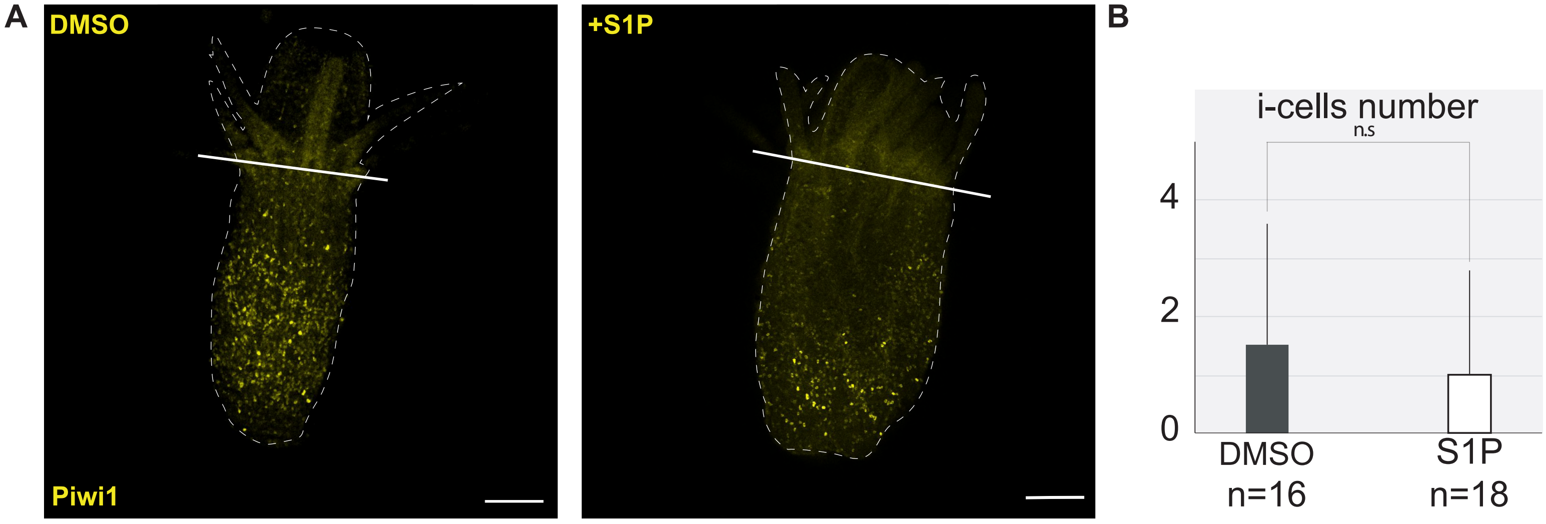


Figure S12

### Figure S13

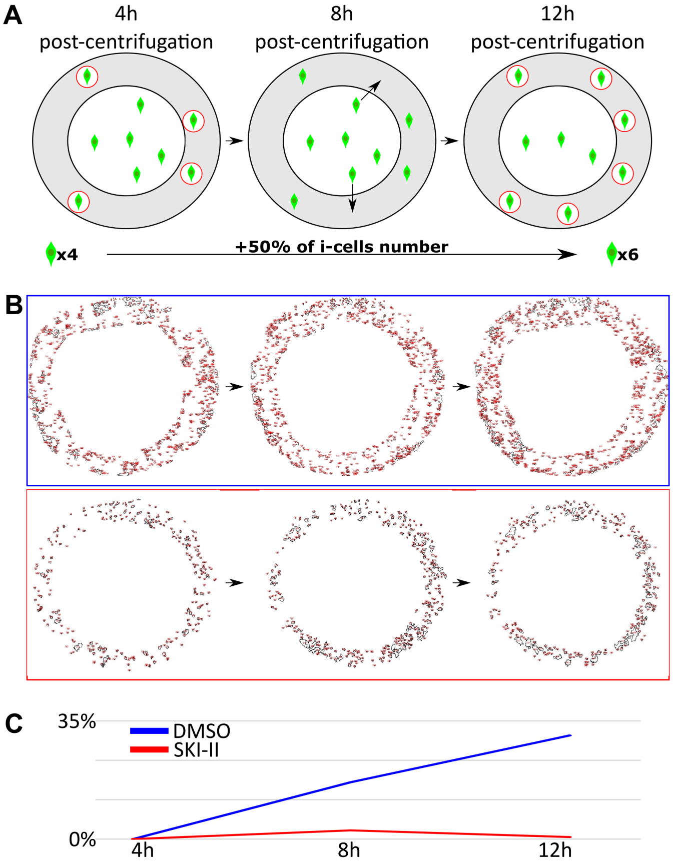

### Figure S14

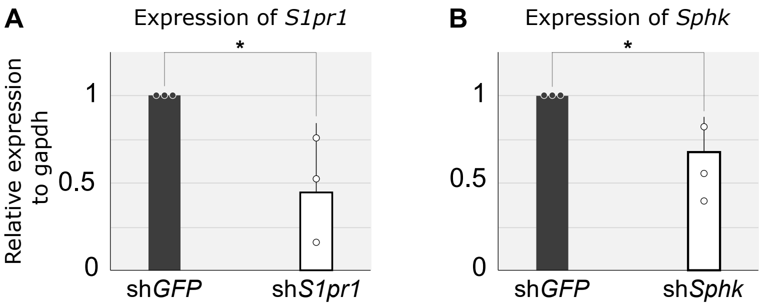

### Figure S15

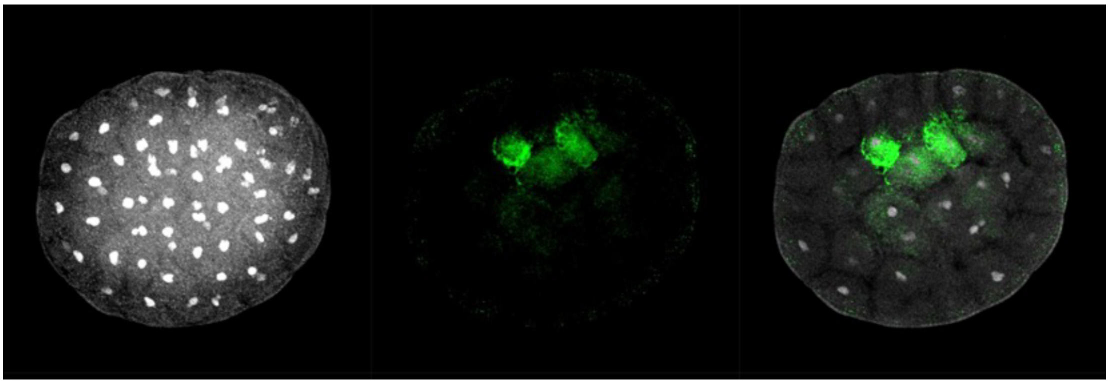

Figure S15
