## Supplementary material for "An unexpected mode of whole-body regeneration from reaggregated cell suspension in *Hydractinia* (Cnidaria, Hydrozoa)": Figure S6

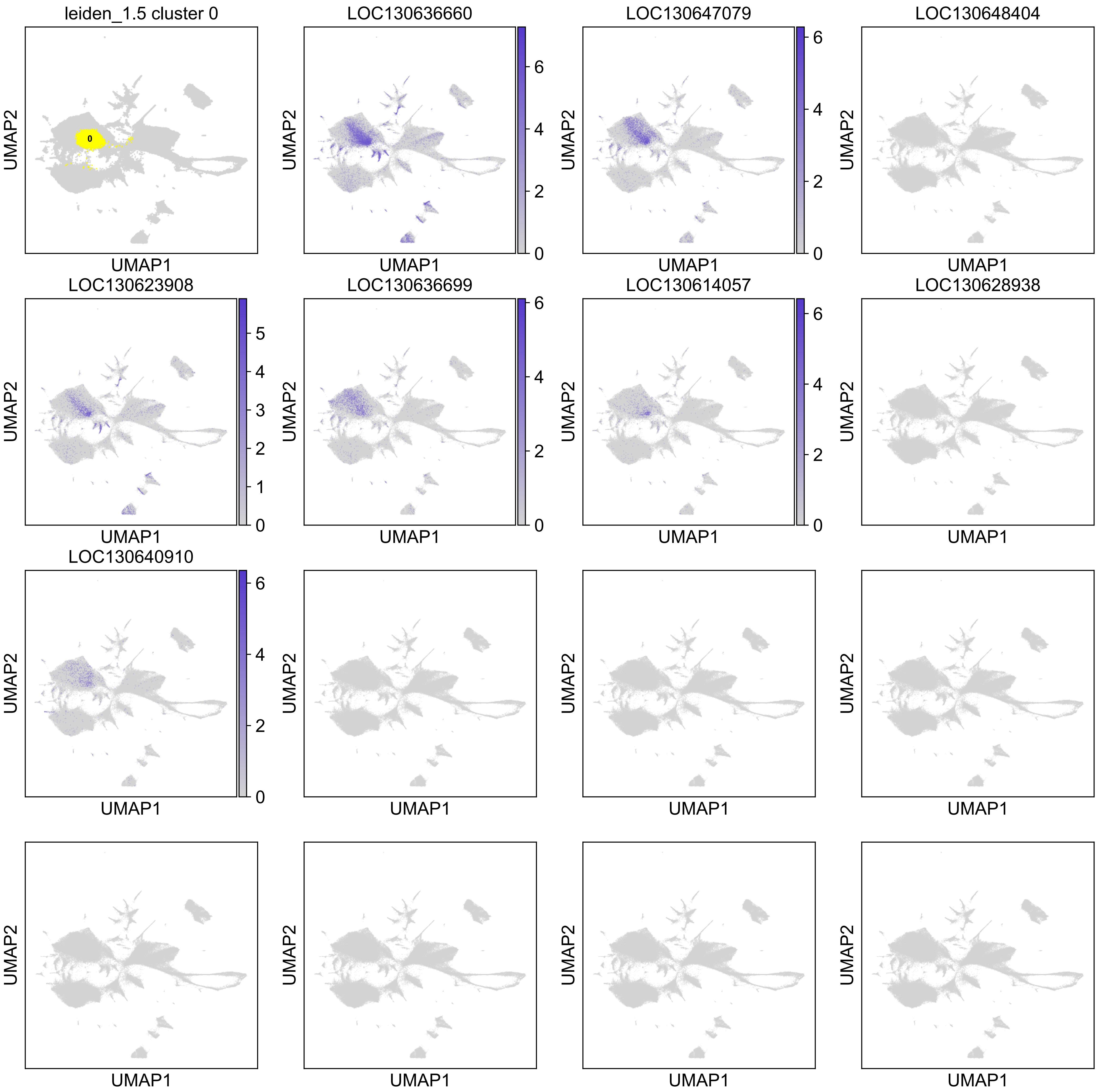

leiden_1.5 cluster 0
0
LOC130636660
LOC130647079
LOC130648404
LOC130623908
LOC130636699
LOC130614057
LOC130628938
LOC130640910
UMAP1
UMAP2

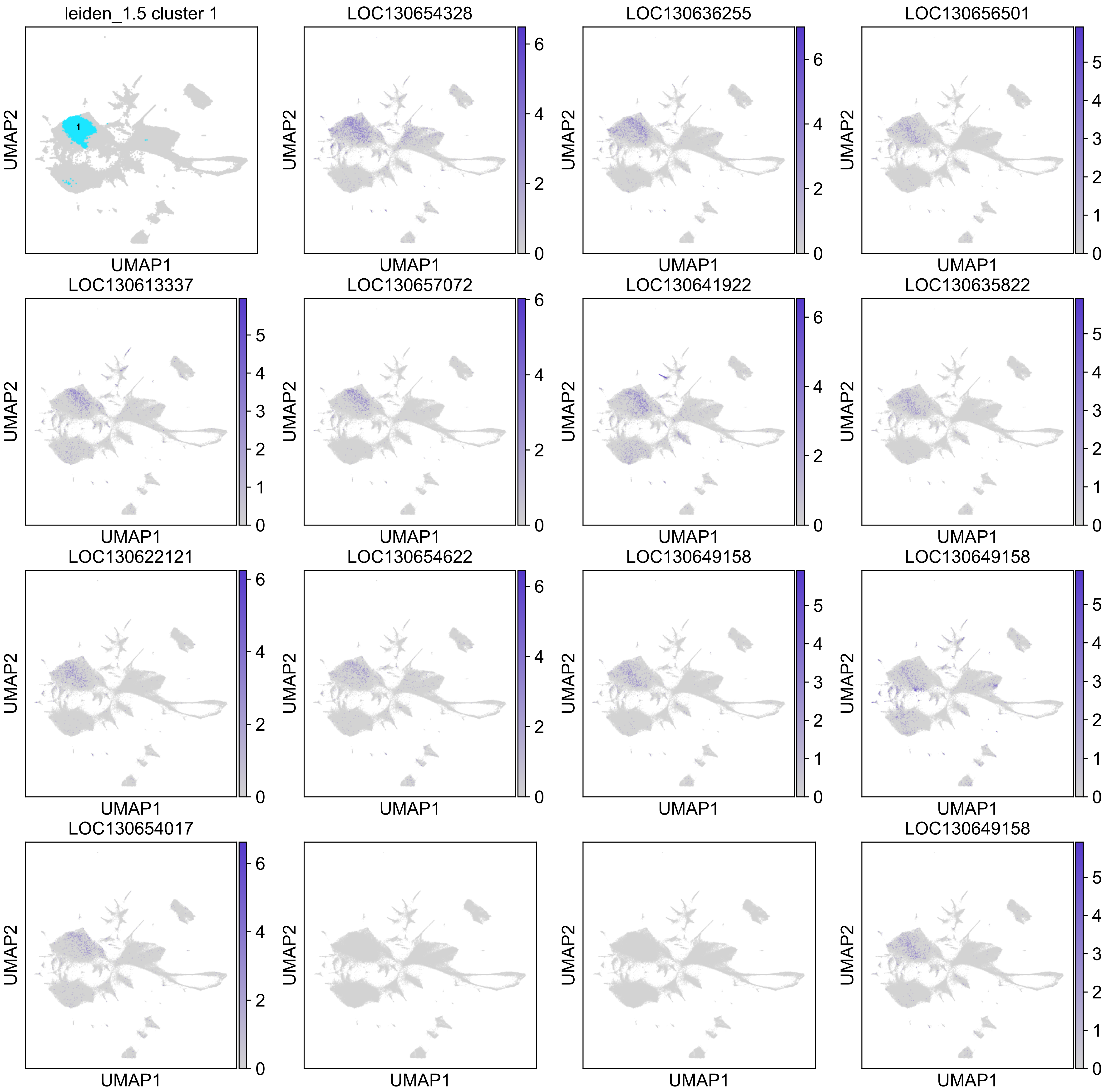

leiden_1.5 cluster 1
LOC130654328
LOC130636255
LOC130656501
LOC130613337
LOC130657072
LOC130641922
LOC130635822
LOC130622121
LOC130654622
LOC130649158
LOC130649158
LOC130654017
LOC130649158
UMAP1
UMAP2

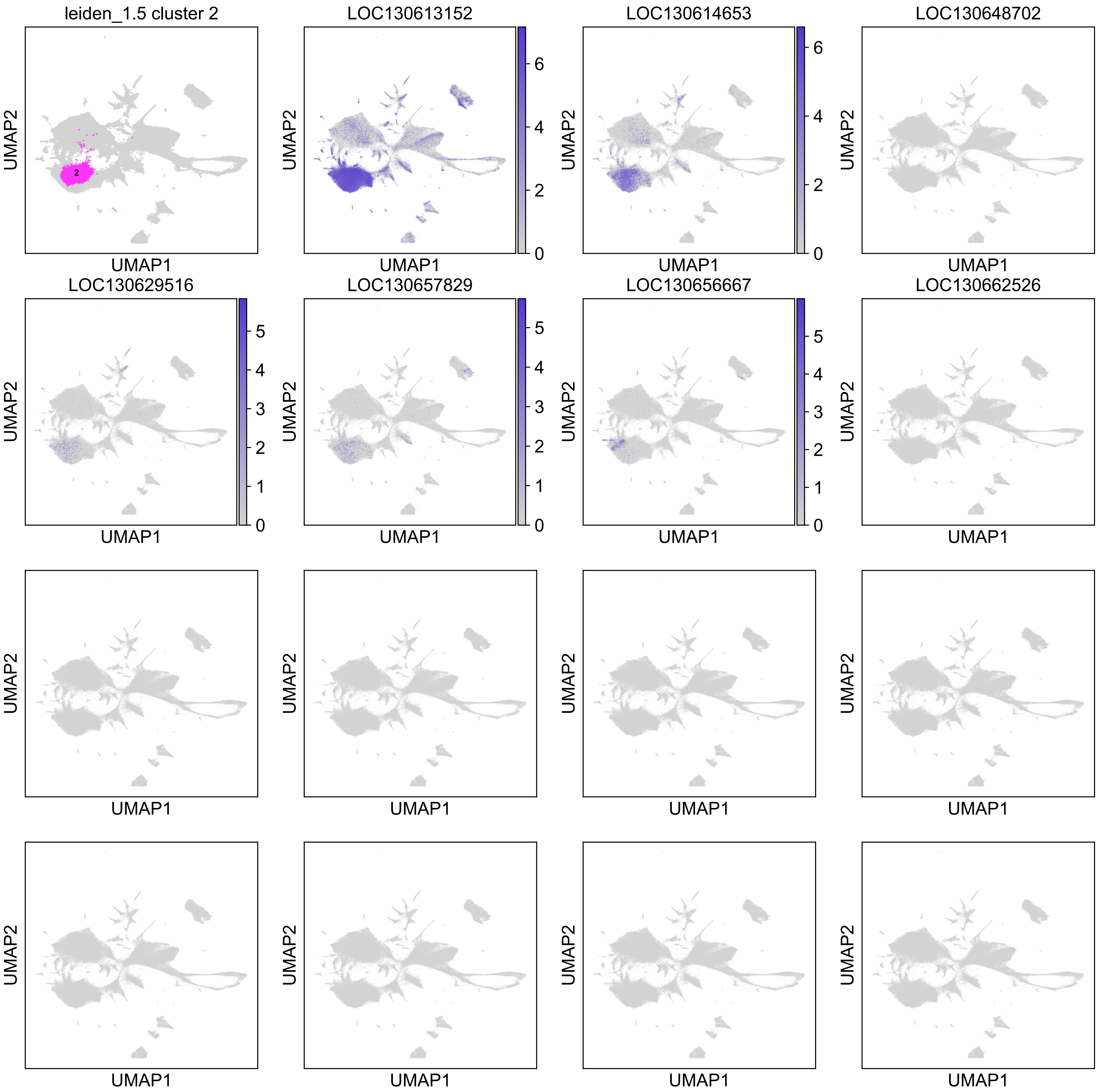
leiden_1.5 cluster 2
2
LOC130613152
LOC130614653
LOC130648702
LOC130629516
LOC130657829
LOC130656667
LOC130662526
UMAP1
UMAP2
0
1
2
3
4
5
6

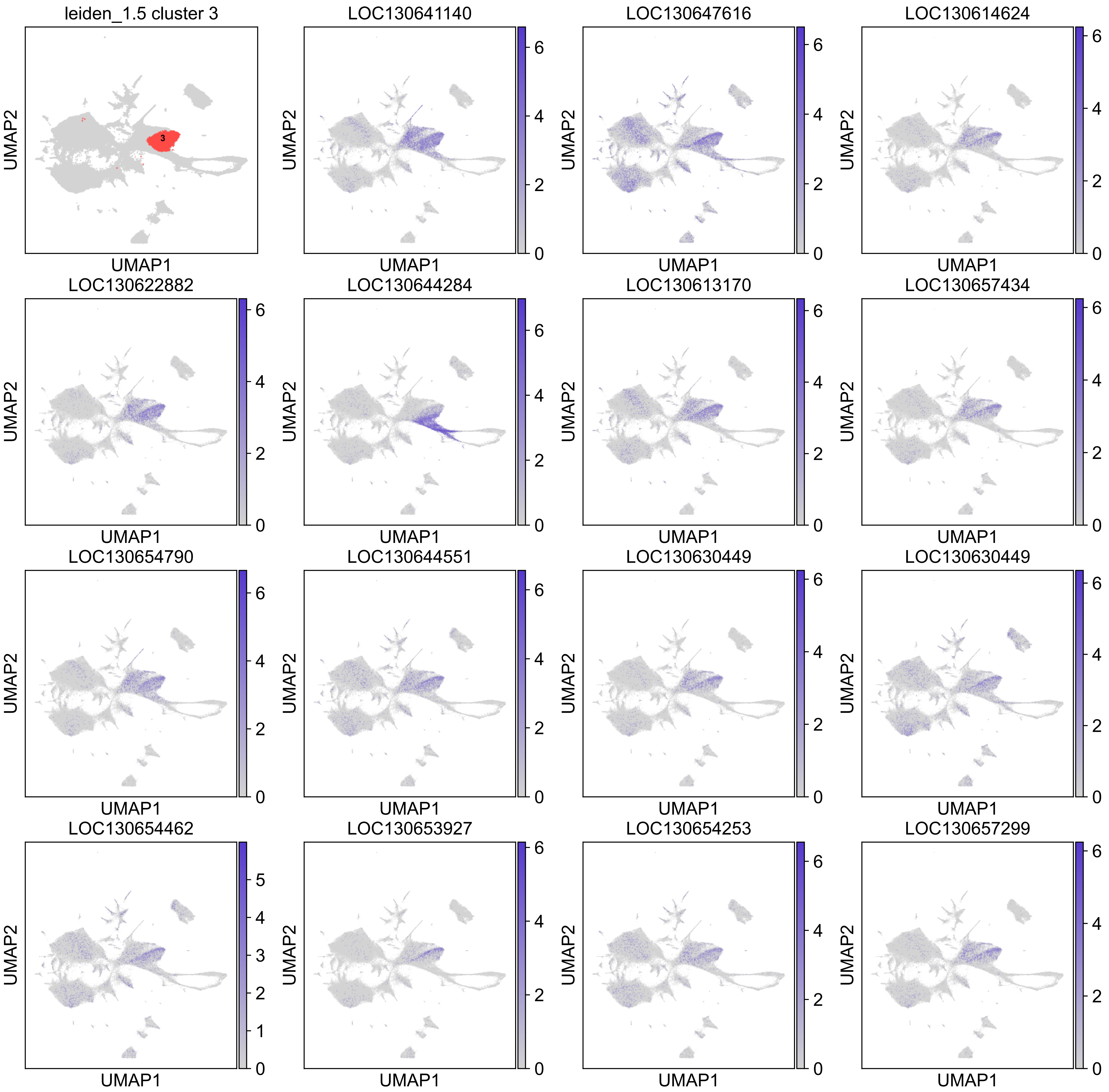
leiden_1.5 cluster 3
3
LOC130641140
LOC130647616
LOC130614624
LOC130622882
LOC130644284
LOC130613170
LOC130657434
LOC130654790
LOC130644551
LOC130630449
LOC130630449
LOC130654462
LOC130653927
LOC130654253
LOC130657299
UMAP1
UMAP2

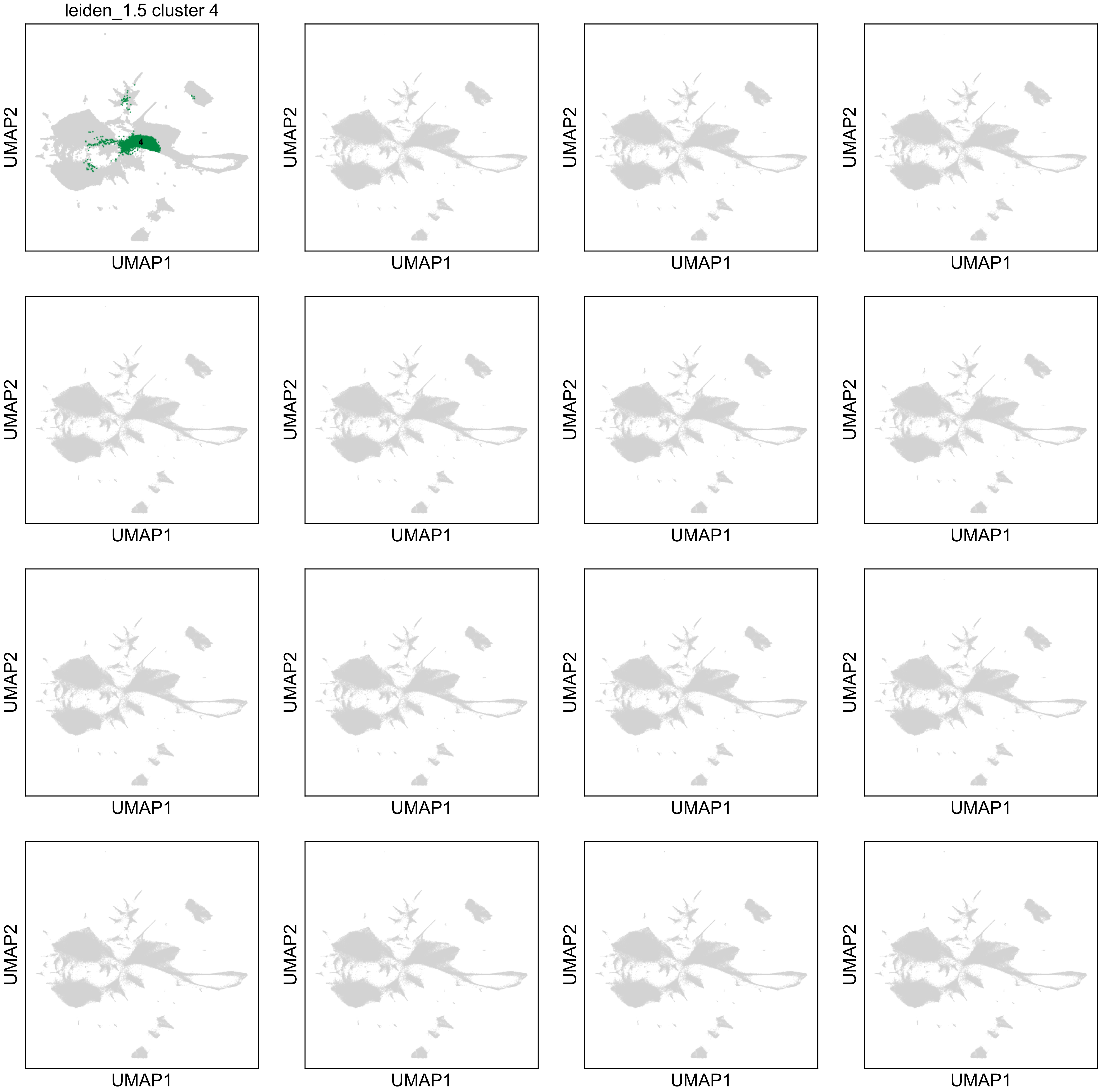
leiden_1.5 cluster 4
4
UMAP2
UMAP1

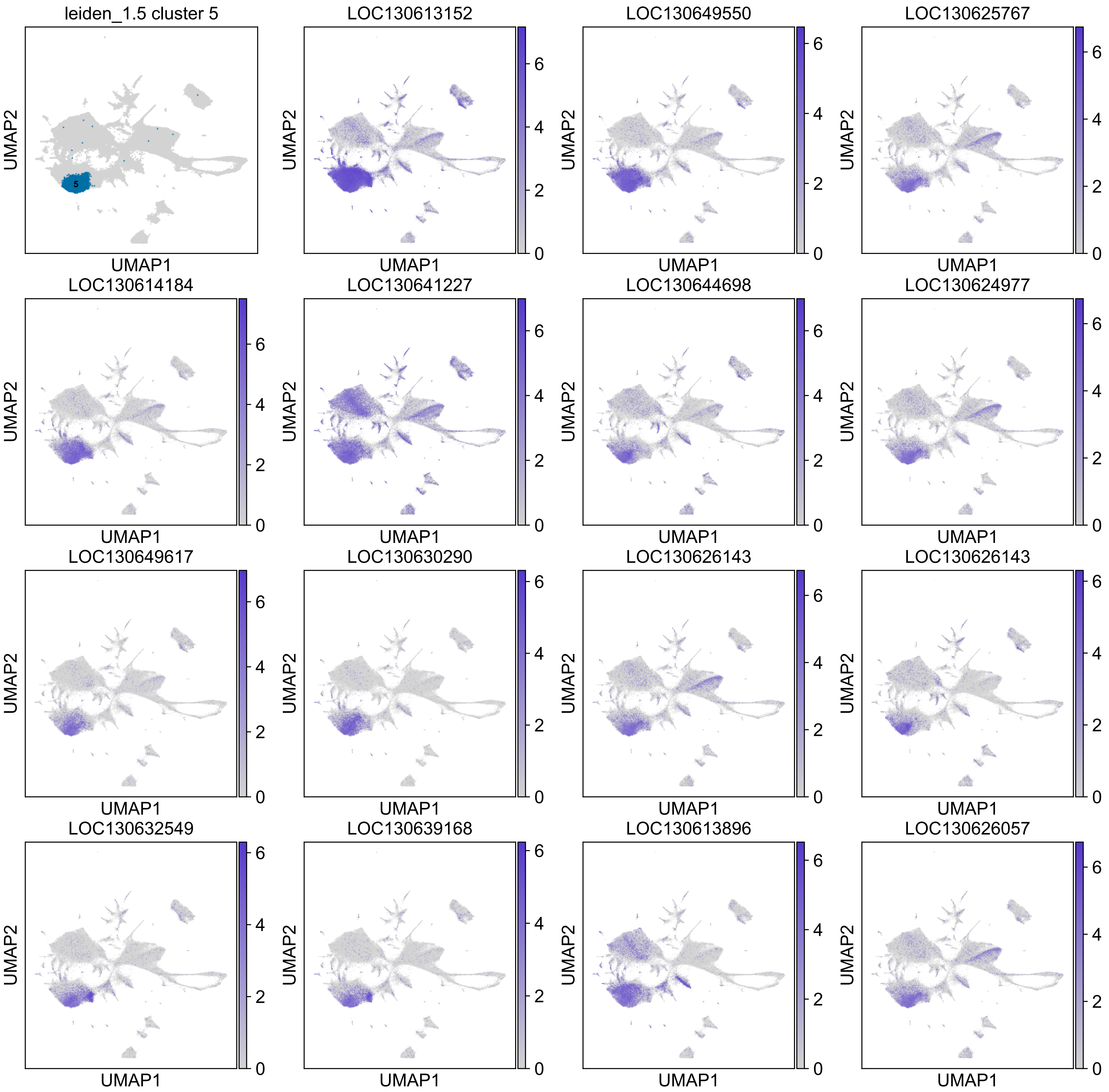

leiden_1.5 cluster 5
LOC130613152
LOC130649550
LOC130625767
LOC130614184
LOC130641227
LOC130644698
LOC130624977
LOC130649617
LOC130630290
LOC130626143
LOC130626143
LOC130632549
LOC130639168
LOC130613896
LOC130626057
UMAP1
UMAP2

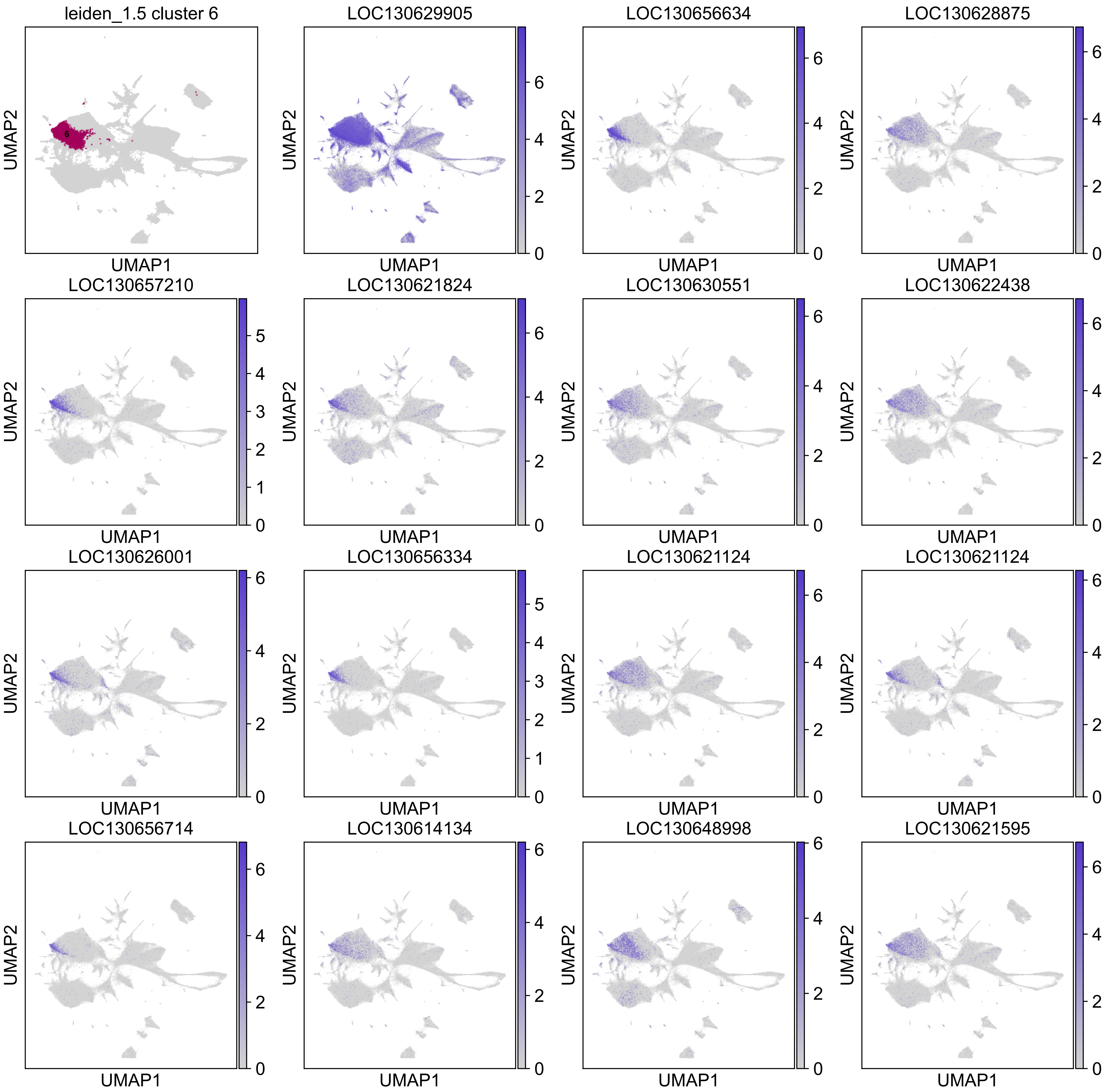

leiden_1.5 cluster 6
6
LOC130629905
LOC130656634
LOC130628875
LOC130657210
LOC130621824
LOC130630551
LOC130622438
LOC130626001
LOC130656334
LOC130621124
LOC130621124
LOC130656714
LOC130614134
LOC130648998
LOC130621595
UMAP1
UMAP2
0
1
2
3
4
5
6

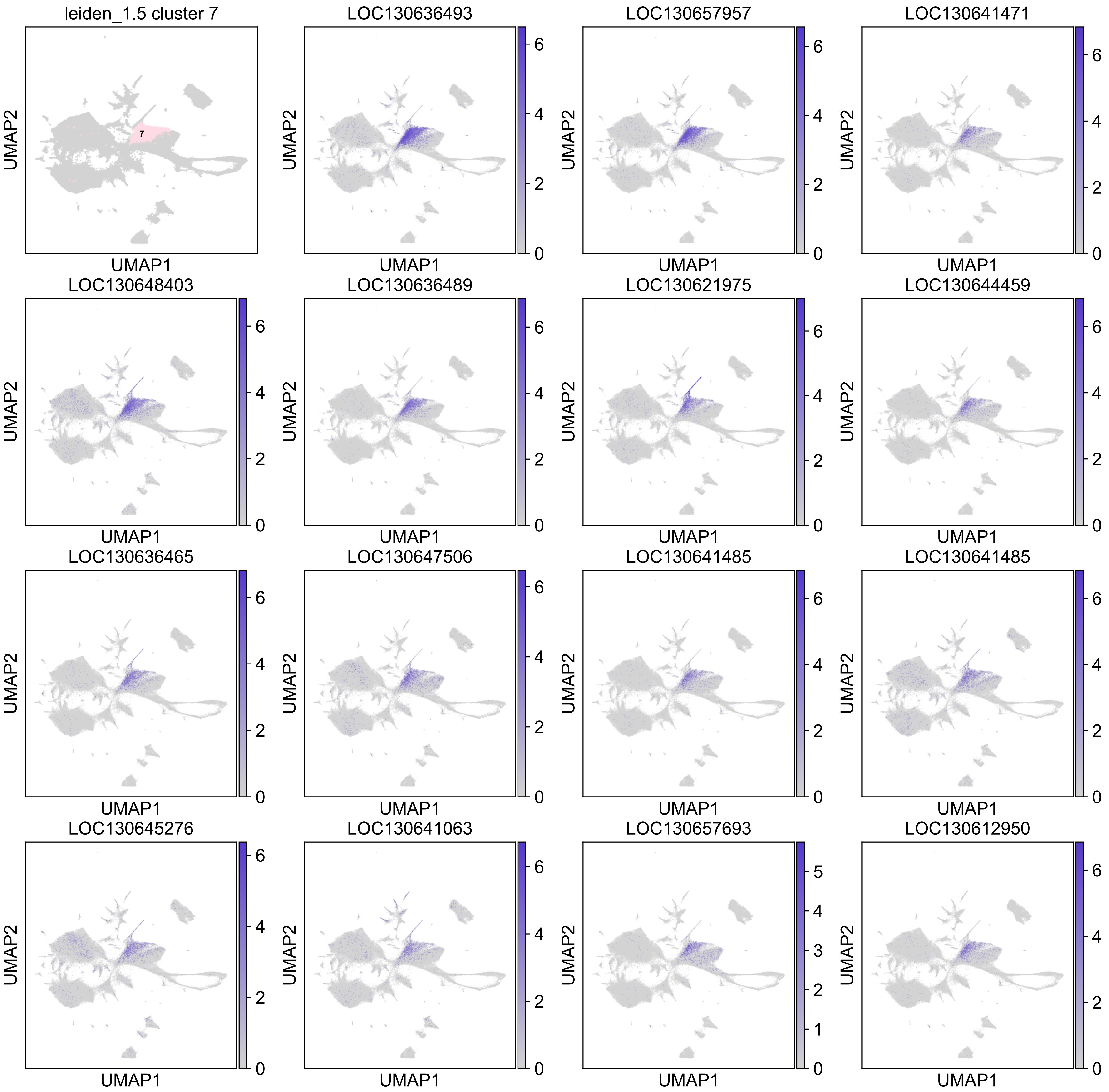
leiden_1.5 cluster 7
LOC130636493
LOC130657957
LOC130641471
LOC130648403
LOC130636489
LOC130621975
LOC130644459
LOC130636465
LOC130647506
LOC130641485
LOC130641485
LOC130645276
LOC130641063
LOC130657693
LOC130612950
UMAP1
UMAP2

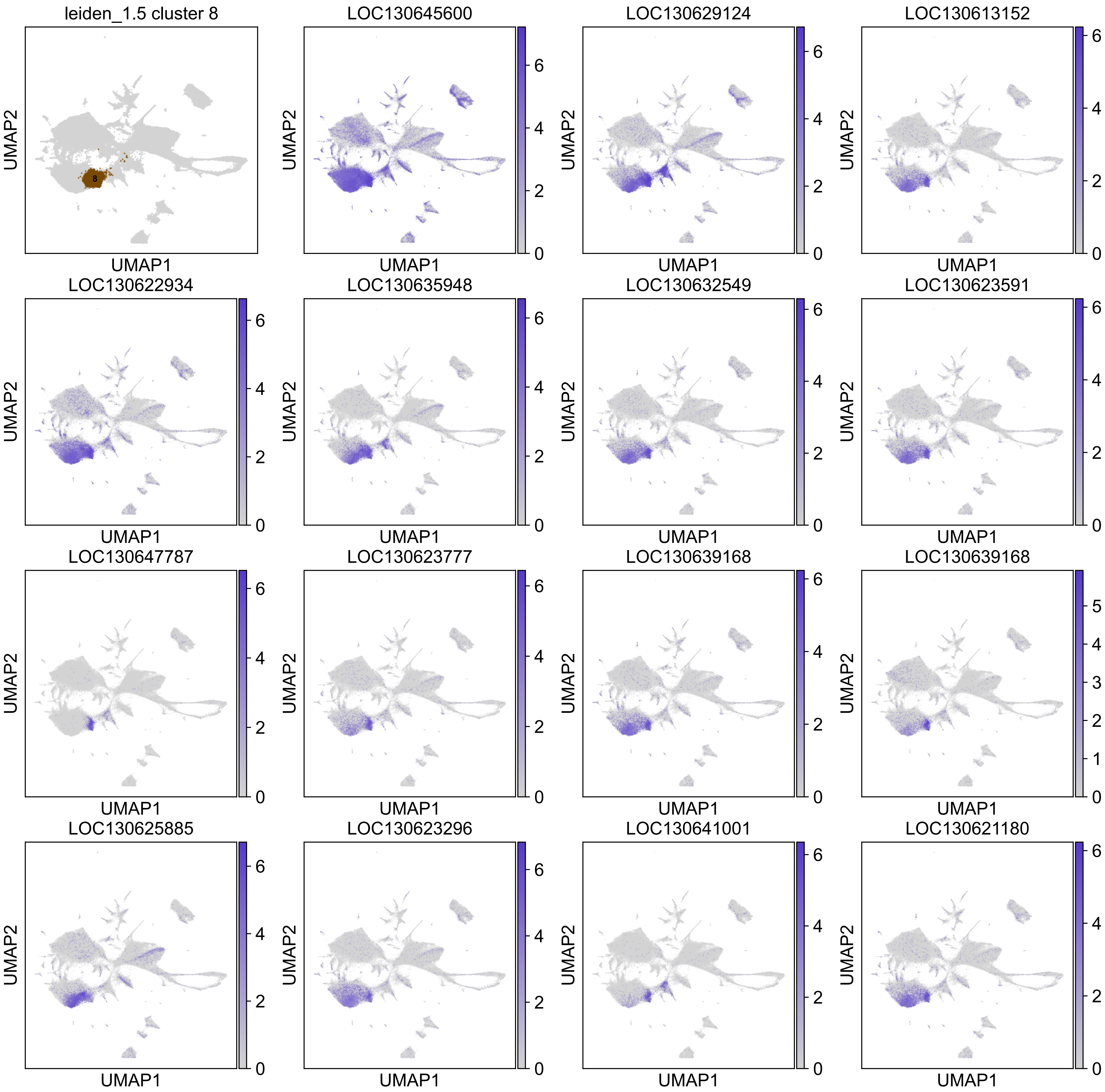

leiden_1.5 cluster 8
8
LOC130645600
LOC130629124
LOC130613152
LOC130622934
LOC130635948
LOC130632549
LOC130623591
LOC130647787
LOC130623777
LOC130639168
LOC130639168
LOC130625885
LOC130623296
LOC130641001
LOC130621180
UMAP1
UMAP2

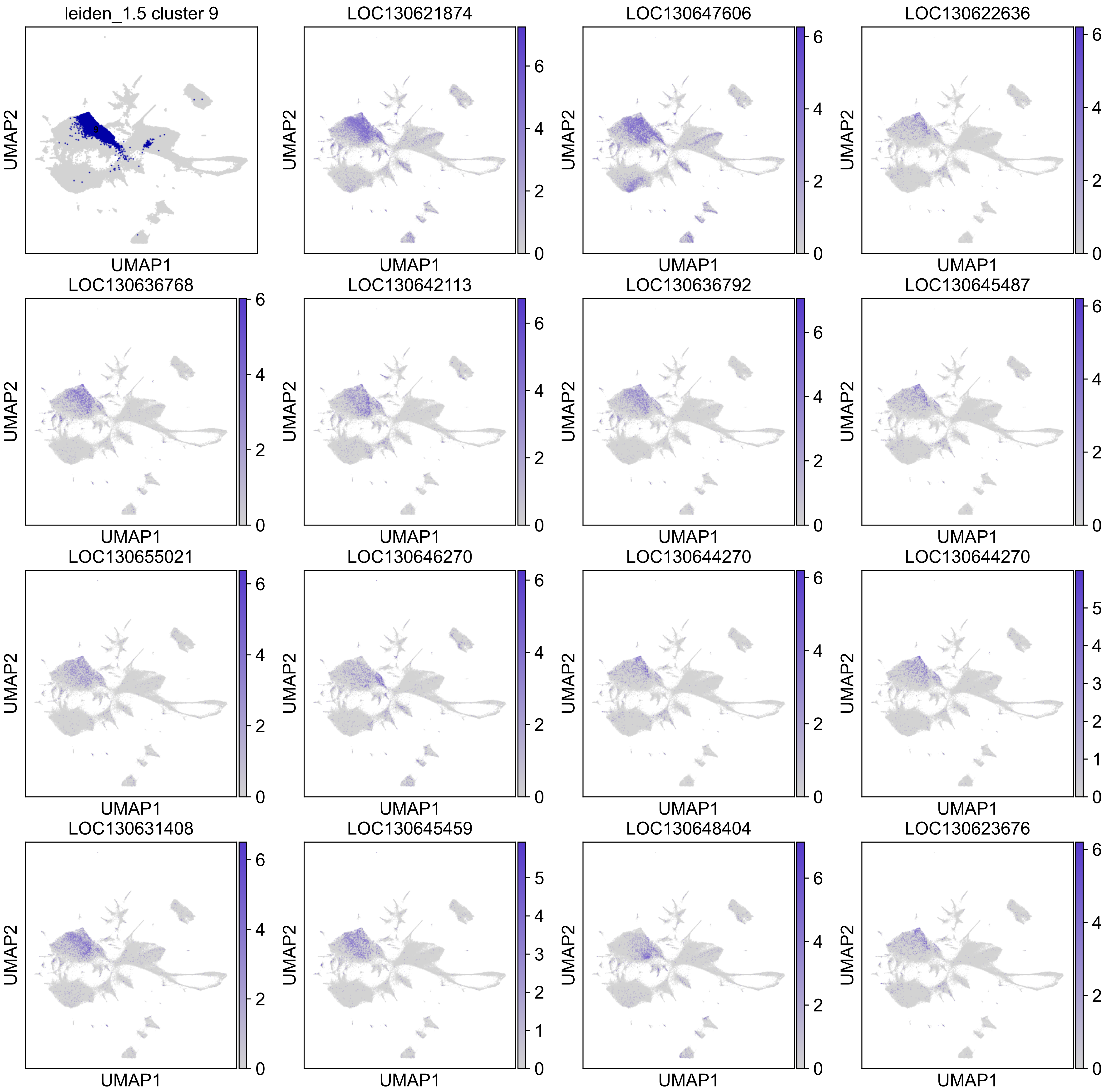

leiden_1.5 cluster 9
LOC130621874
LOC130647606
LOC130622636
LOC130636768
LOC130642113
LOC130636792
LOC130645487
LOC130655021
LOC130646270
LOC130644270
LOC130644270
LOC130631408
LOC130645459
LOC130648404
LOC130623676
UMAP1
UMAP2

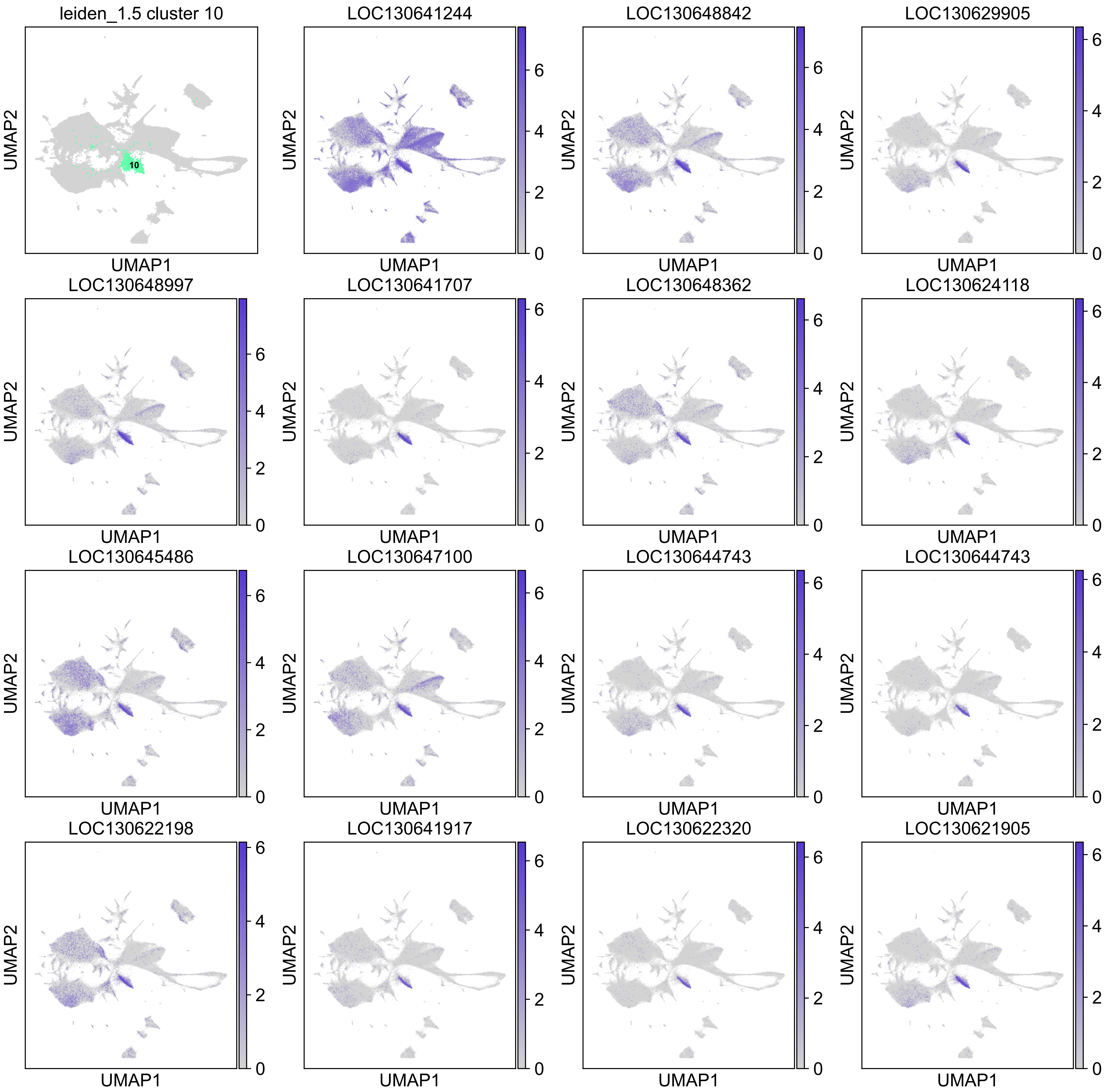
leiden_1.5 cluster 10
10
LOC130641244
LOC130648842
LOC130629905
LOC130648997
LOC130641707
LOC130648362
LOC130624118
LOC130645486
LOC130647100
LOC130644743
LOC130644743
LOC130622198
LOC130641917
LOC130622320
LOC130621905
UMAP1
UMAP2
0
2
4
6

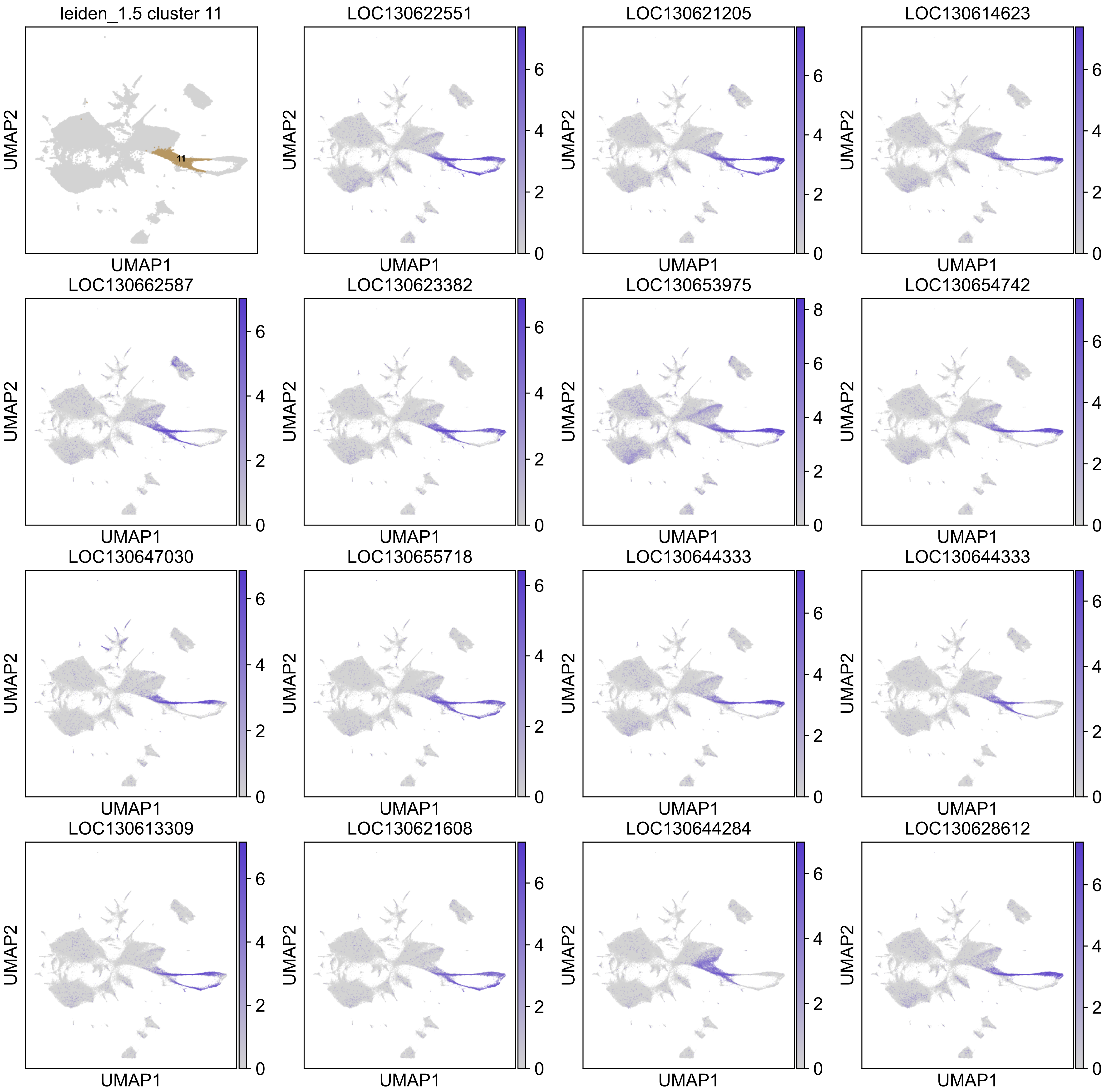
leiden_1.5 cluster 11
11
LOC130622551
LOC130621205
LOC130614623
LOC130662587
LOC130623382
LOC130653975
LOC130654742
LOC130647030
LOC130655718
LOC130644333
LOC130644333
LOC130613309
LOC130621608
LOC130644284
LOC130628612
UMAP1
UMAP2

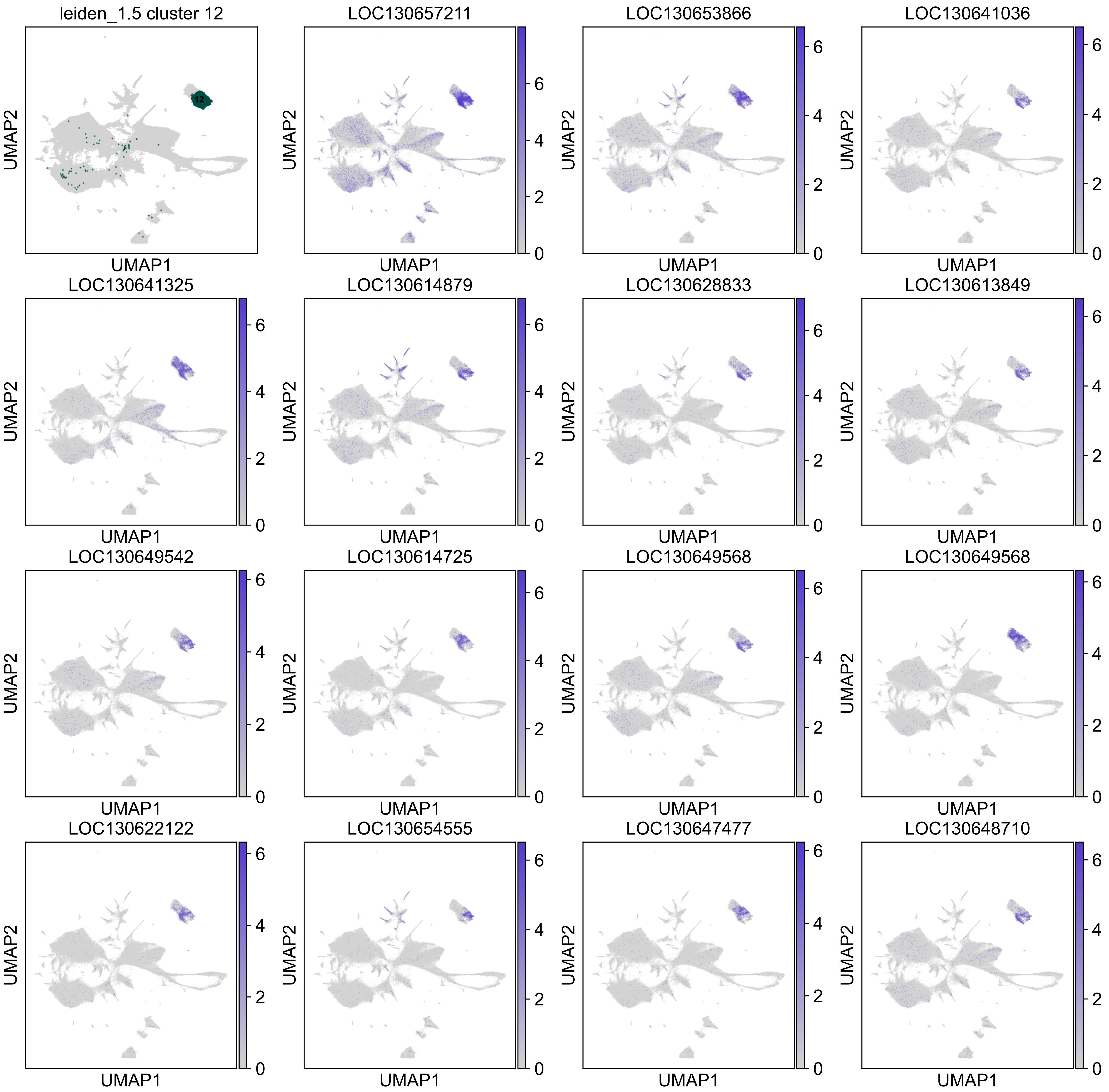

leiden_1.5 cluster 12
12
LOC130657211
LOC130653866
LOC130641036
LOC130641325
LOC130614879
LOC130628833
LOC130613849
LOC130649542
LOC130614725
LOC130649568
LOC130649568
LOC130622122
LOC130654555
LOC130647477
LOC130648710
UMAP1
UMAP2
0
2
4
6

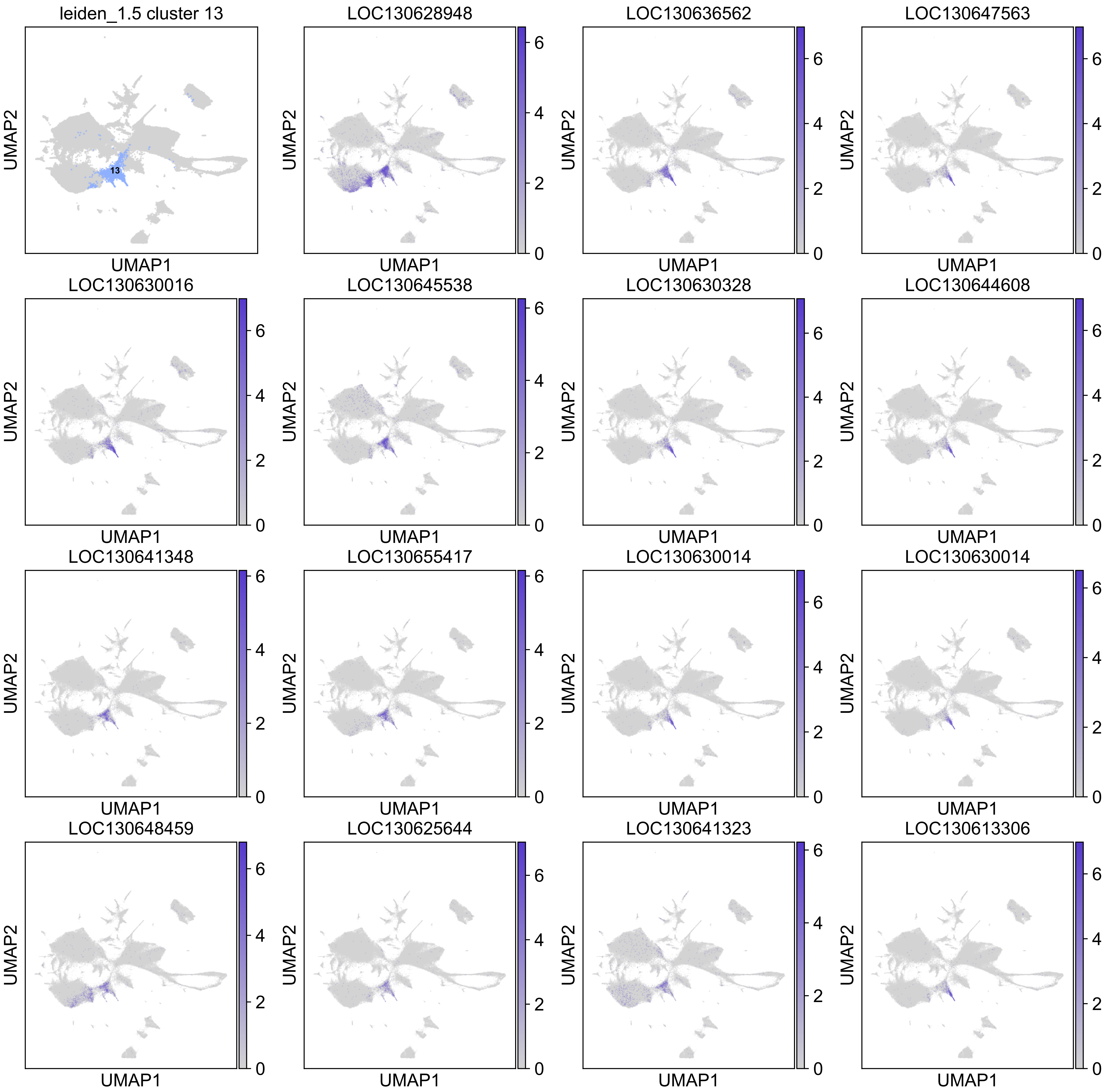
leiden_1.5 cluster 13
13
LOC130628948
LOC130636562
LOC130647563
LOC130630016
LOC130645538
LOC130630328
LOC130644608
LOC130641348
LOC130655417
LOC130630014
LOC130630014
LOC130648459
LOC130625644
LOC130641323
LOC130613306
UMAP1
UMAP2
0
2
4
6

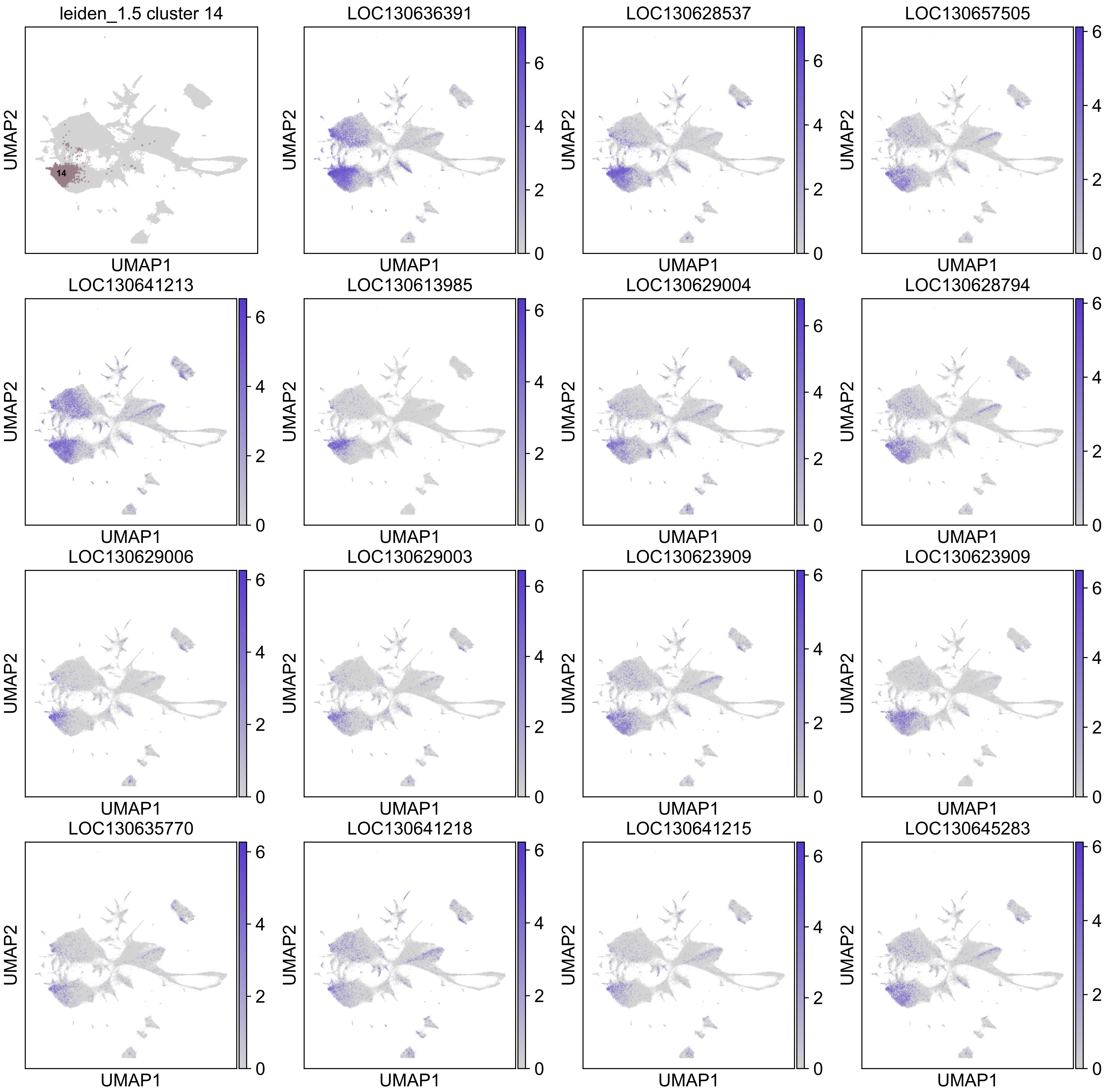
leiden_1.5 cluster 14
14
LOC130636391
LOC130628537
LOC130657505
LOC130641213
LOC130613985
LOC130629004
LOC130628794
LOC130629006
LOC130629003
LOC130623909
LOC130623909
LOC130635770
LOC130641218
LOC130641215
LOC130645283
UMAP1
UMAP2

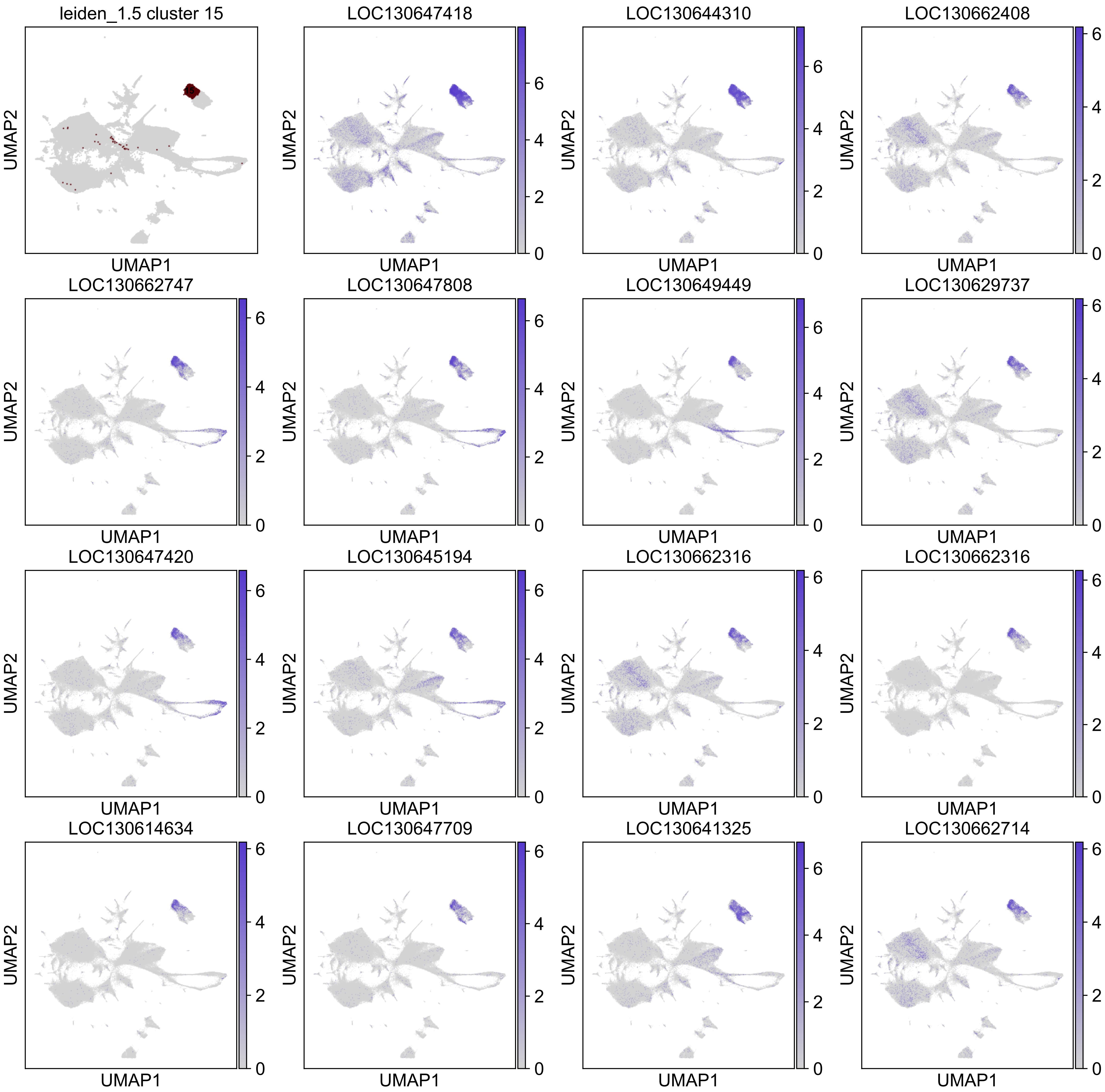
leiden_1.5 cluster 15
LOC130647418
LOC130644310
LOC130662408
LOC130662747
LOC130647808
LOC130649449
LOC130629737
LOC130647420
LOC130645194
LOC130662316
LOC130662316
LOC130614634
LOC130647709
LOC130641325
LOC130662714
UMAP1
UMAP2

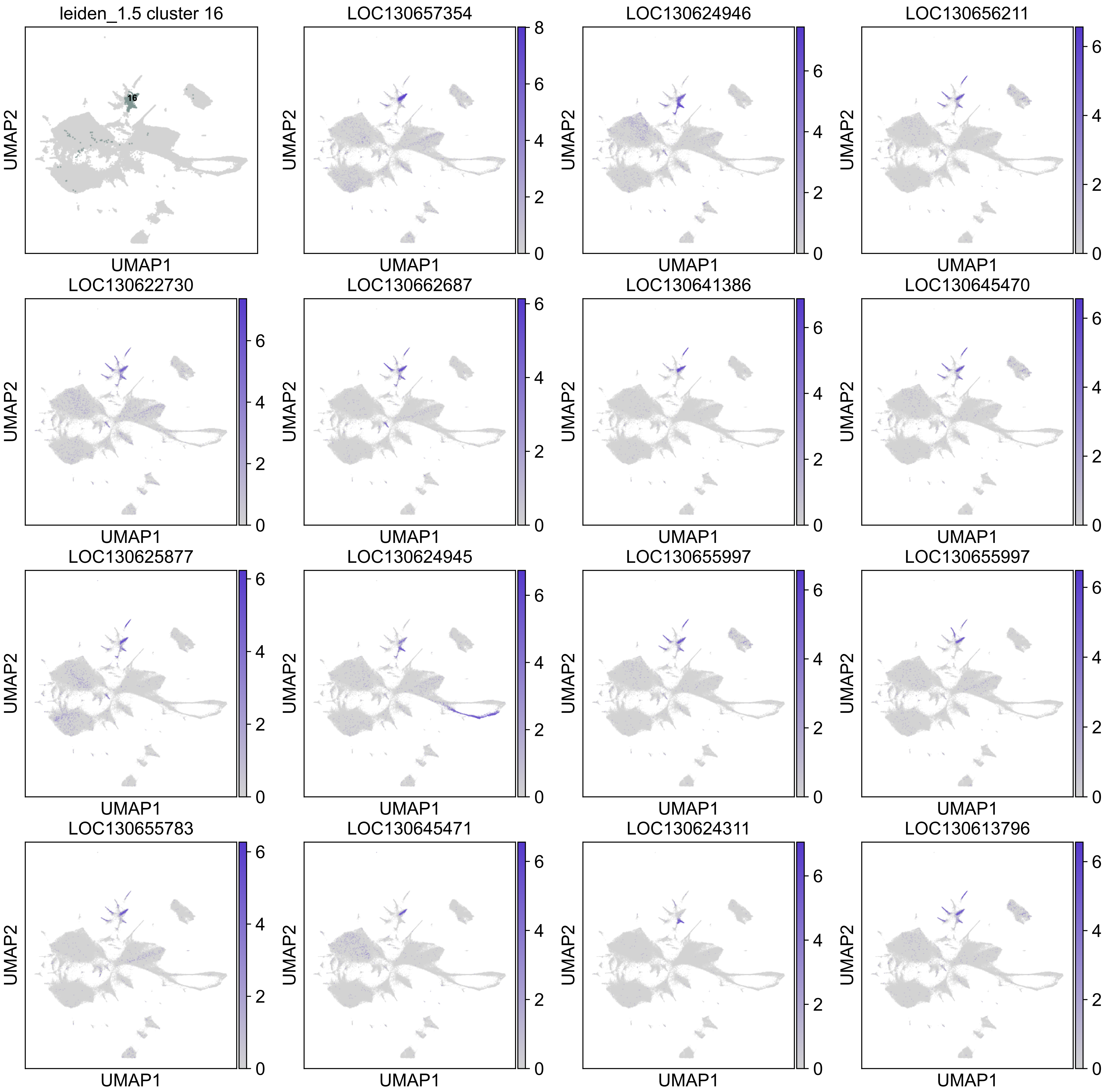
leiden_1.5 cluster 16
16
LOC130657354
LOC130624946
LOC130656211
LOC130622730
LOC130662687
LOC130641386
LOC130645470
LOC130625877
LOC130624945
LOC130655997
LOC130655997
LOC130655783
LOC130645471
LOC130624311
LOC130613796
UMAP1
UMAP2

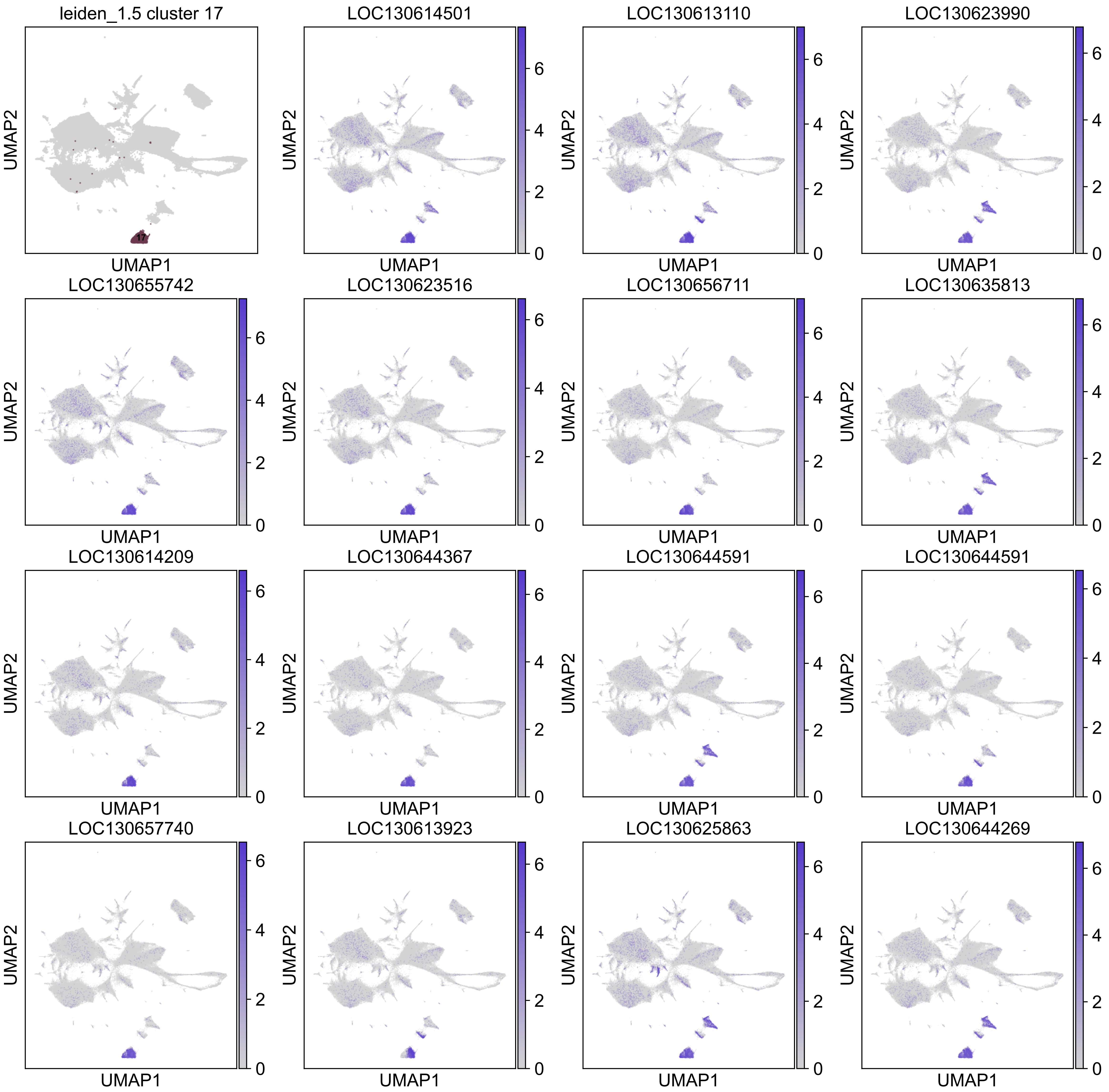
leiden_1.5 cluster 17
LOC130614501
LOC130613110
LOC130623990
LOC130655742
LOC130623516
LOC130656711
LOC130635813
LOC130614209
LOC130644367
LOC130644591
LOC130644591
LOC130657740
LOC130613923
LOC130625863
LOC130644269
UMAP1
UMAP2
17
0
2
4
6

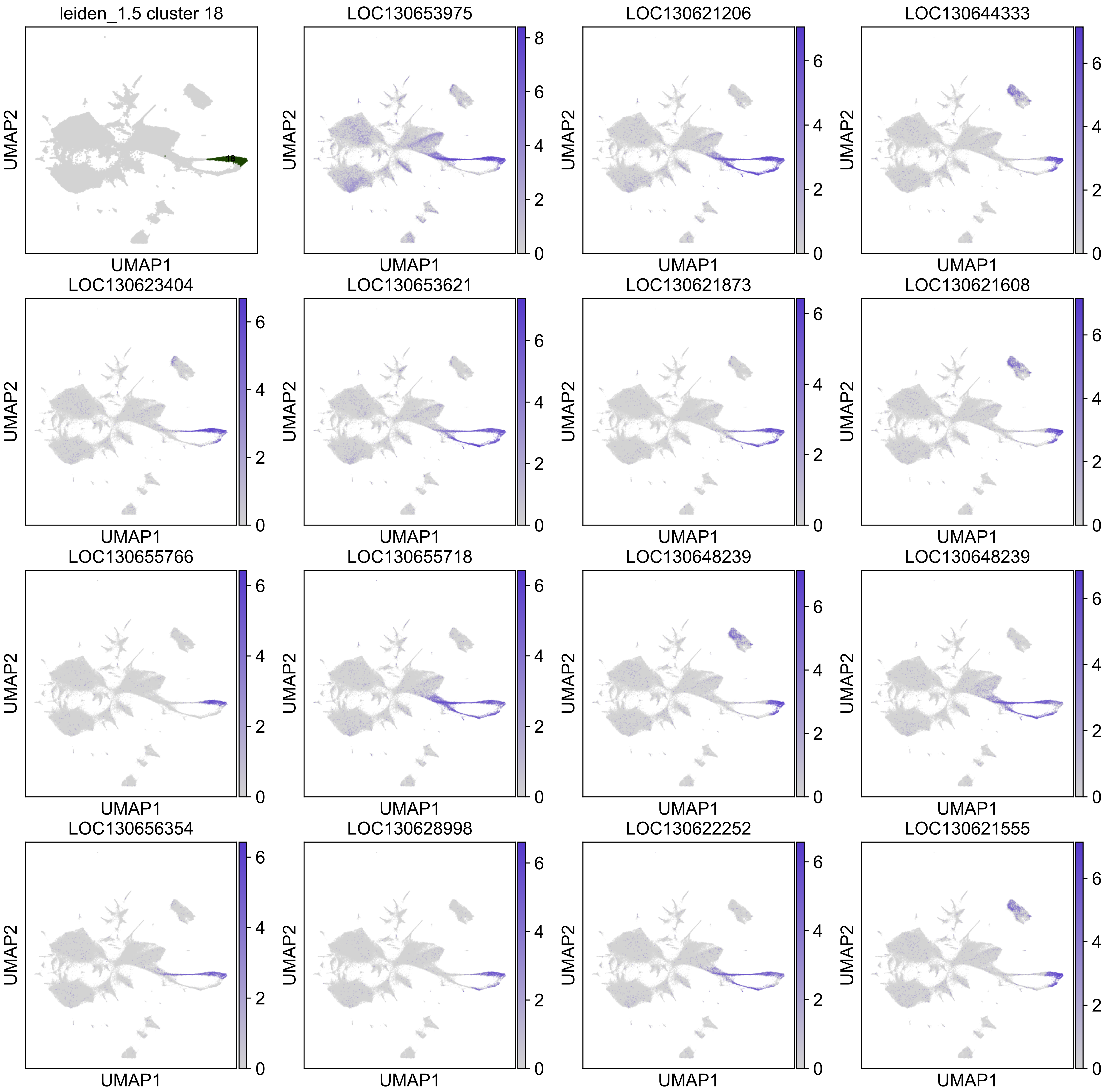
leiden_1.5 cluster 18
LOC130653975
LOC130621206
LOC130644333
LOC130623404
LOC130653621
LOC130621873
LOC130621608
LOC130655766
LOC130655718
LOC130648239
LOC130648239
LOC130656354
LOC130628998
LOC130622252
LOC130621555
UMAP1
UMAP2

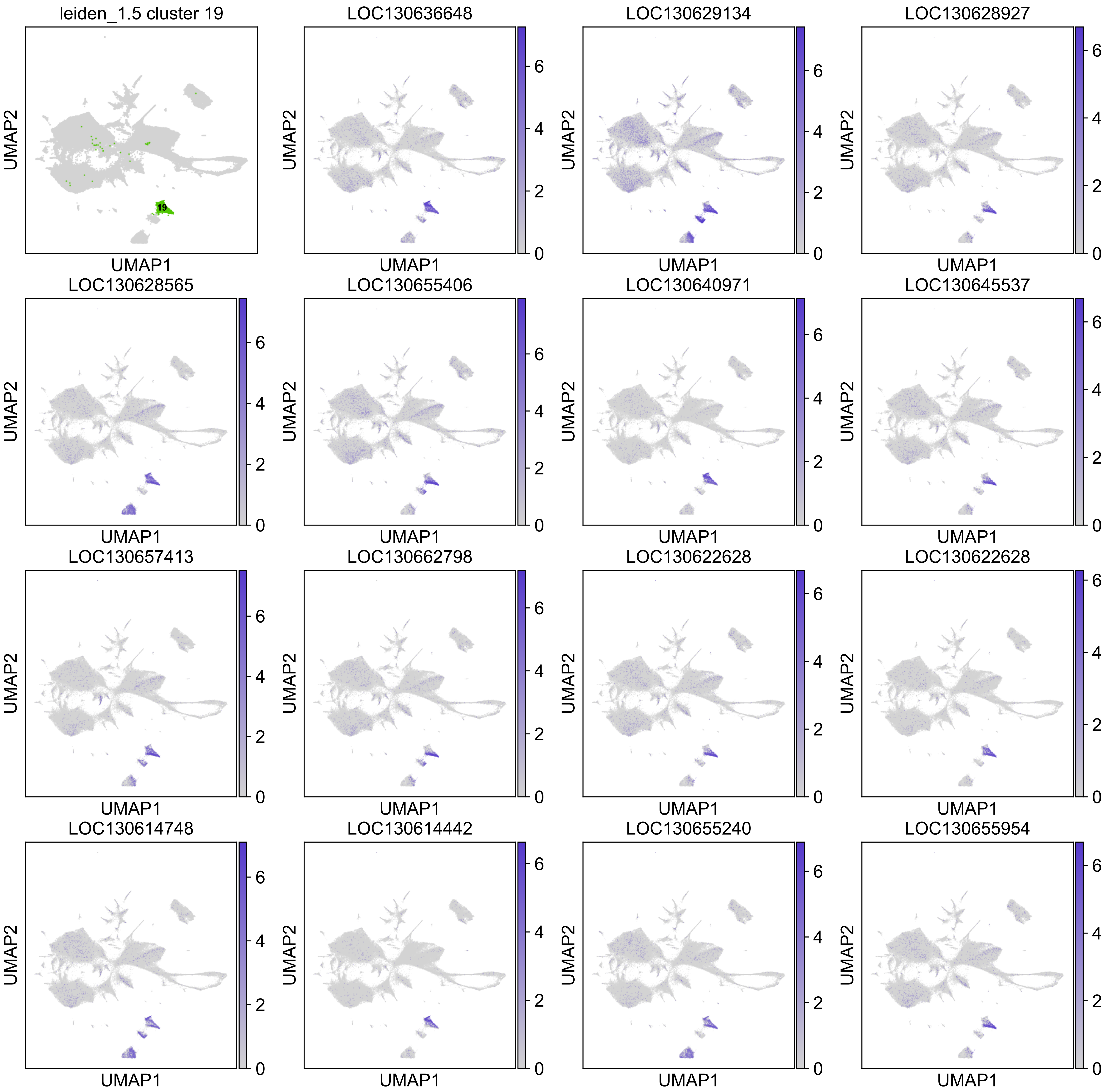
leiden_1.5 cluster 19
LOC130636648
LOC130629134
LOC130628927
LOC130628565
LOC130655406
LOC130640971
LOC130645537
LOC130657413
LOC130662798
LOC130622628
LOC130622628
LOC130614748
LOC130614442
LOC130655240
LOC130655954
UMAP1
UMAP2
19
0
2
4
6

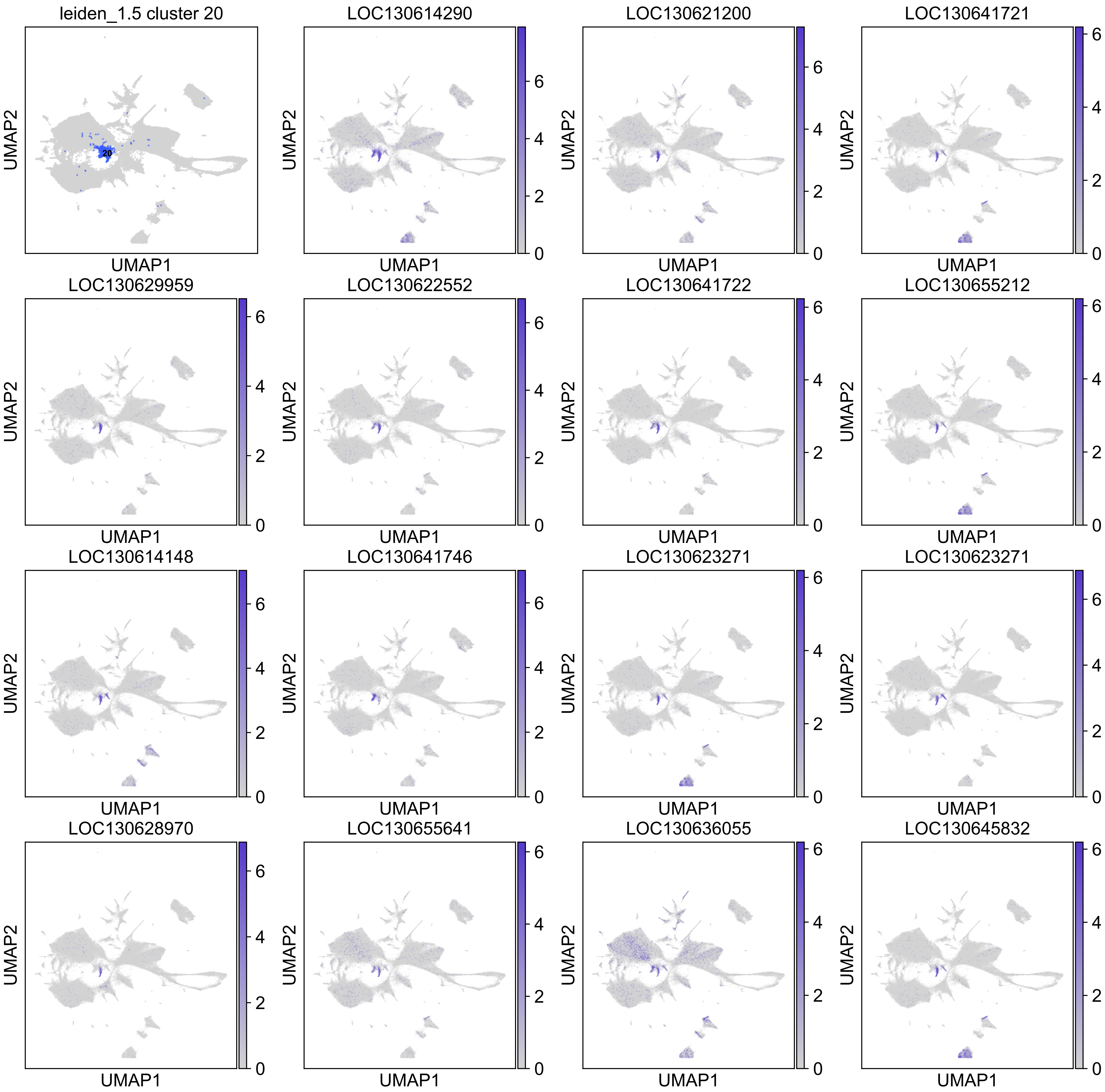
leiden_1.5 cluster 20
20
LOC130614290
LOC130621200
LOC130641721
LOC130629959
LOC130622552
LOC130641722
LOC130655212
LOC130614148
LOC130641746
LOC130623271
LOC130623271
LOC130628970
LOC130655641
LOC130636055
LOC130645832
UMAP1
UMAP2
0
2
4
6

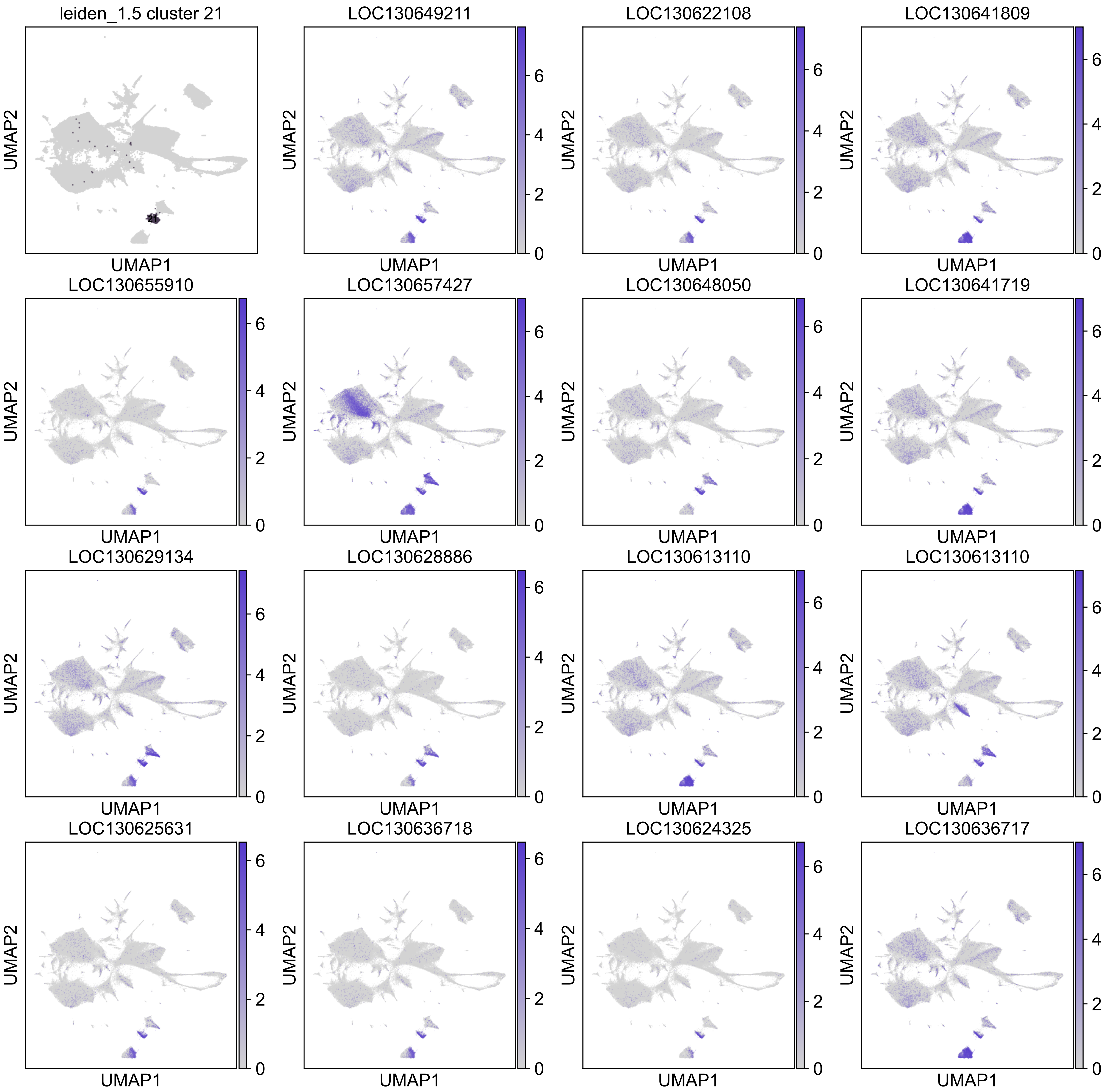
leiden_1.5 cluster 21
LOC130649211
LOC130622108
LOC130641809
LOC130655910
LOC130657427
LOC130648050
LOC130641719
LOC130629134
LOC130628886
LOC130613110
LOC130613110
LOC130625631
LOC130636718
LOC130624325
LOC130636717
UMAP1
UMAP2

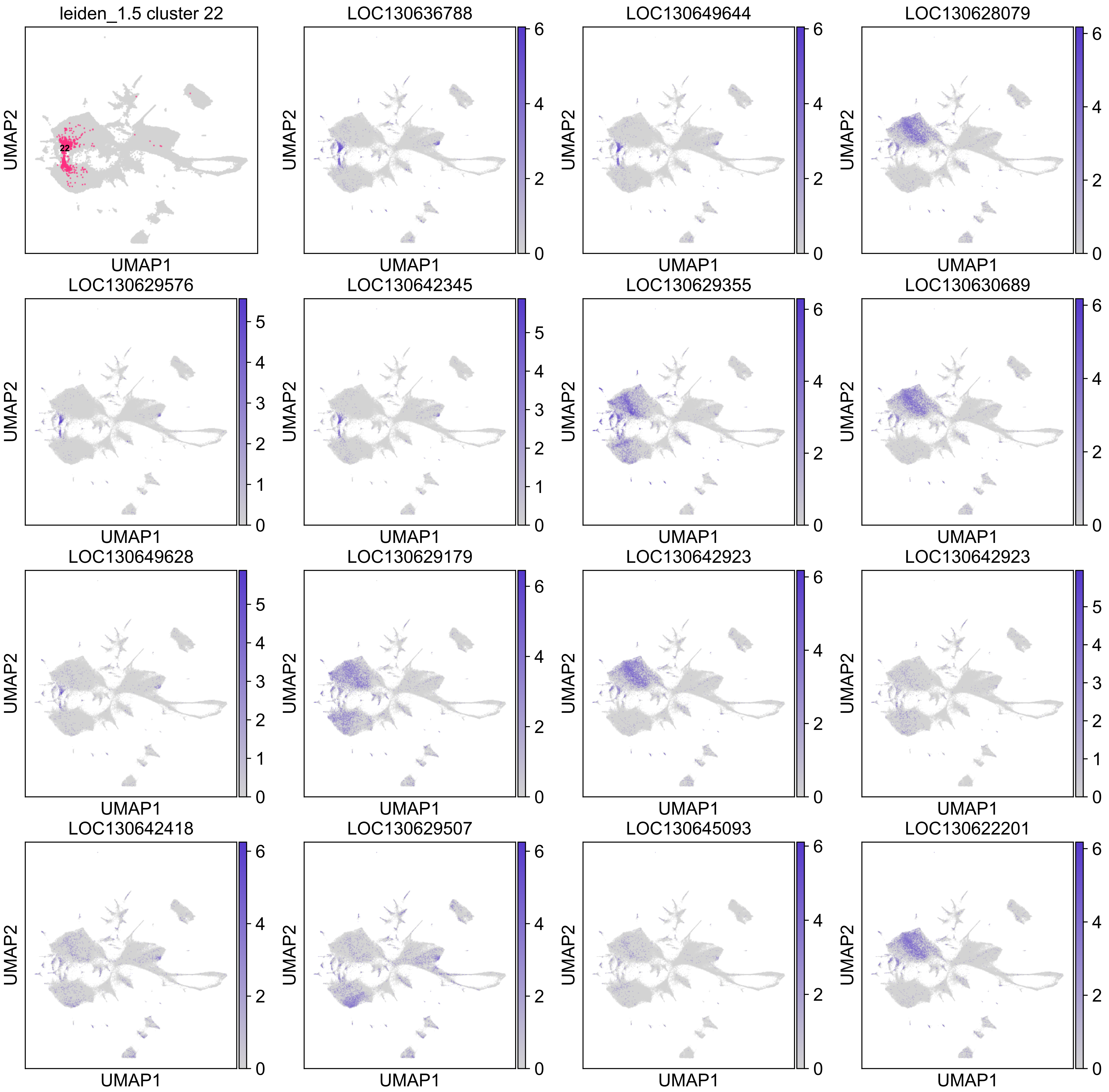
leiden_1.5 cluster 22
22
LOC130636788
LOC130649644
LOC130628079
LOC130629576
LOC130642345
LOC130629355
LOC130630689
LOC130649628
LOC130629179
LOC130642923
LOC130642923
LOC130642418
LOC130629507
LOC130645093
LOC130622201
UMAP1
UMAP2

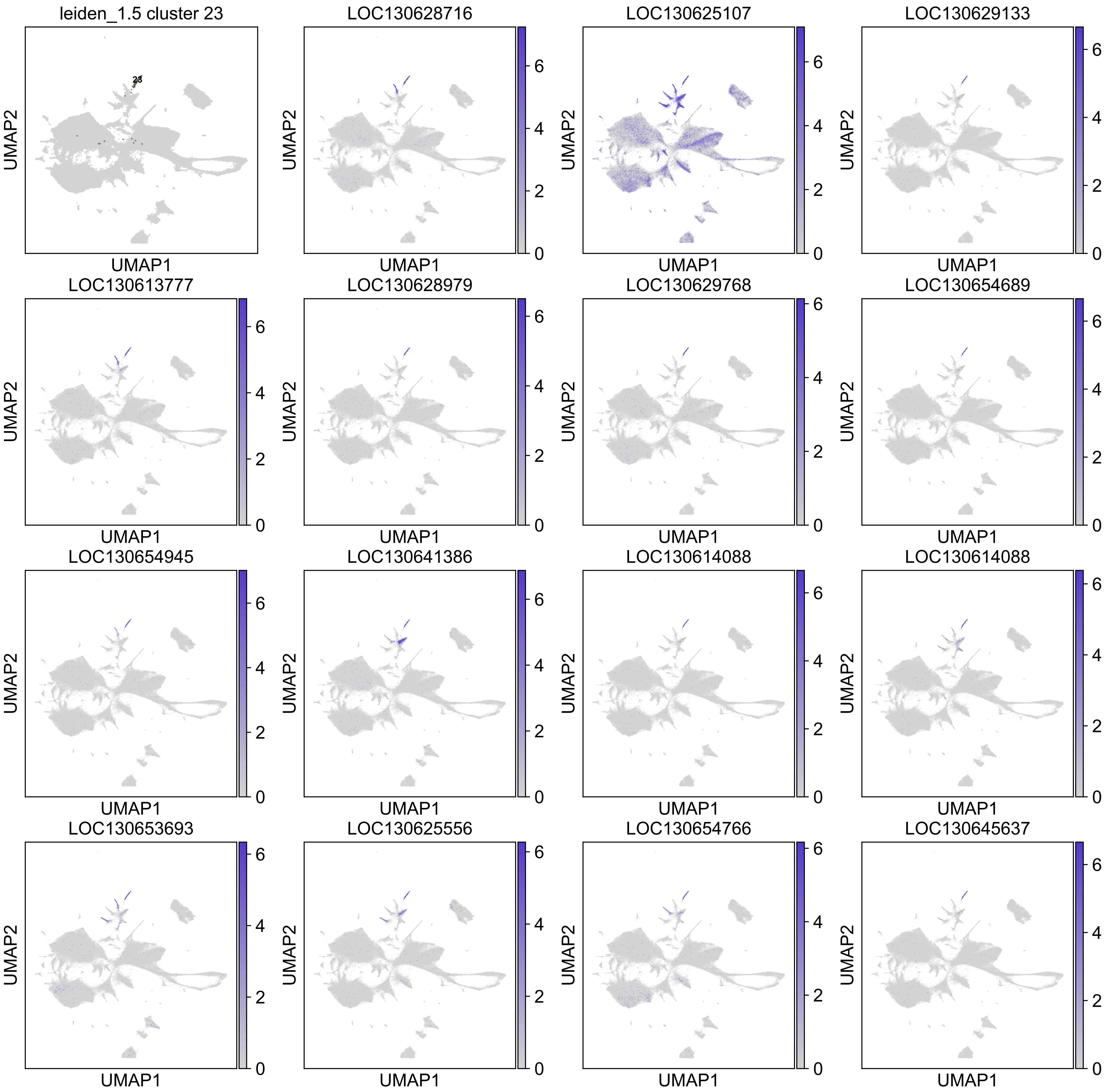

leiden_1.5 cluster 23
23
LOC130628716
LOC130625107
LOC130629133
LOC130613777
LOC130628979
LOC130629768
LOC130654689
LOC130654945
LOC130641386
LOC130614088
LOC130614088
LOC130653693
LOC130625556
LOC130654766
LOC130645637
UMAP1
UMAP2
0
2
4
6

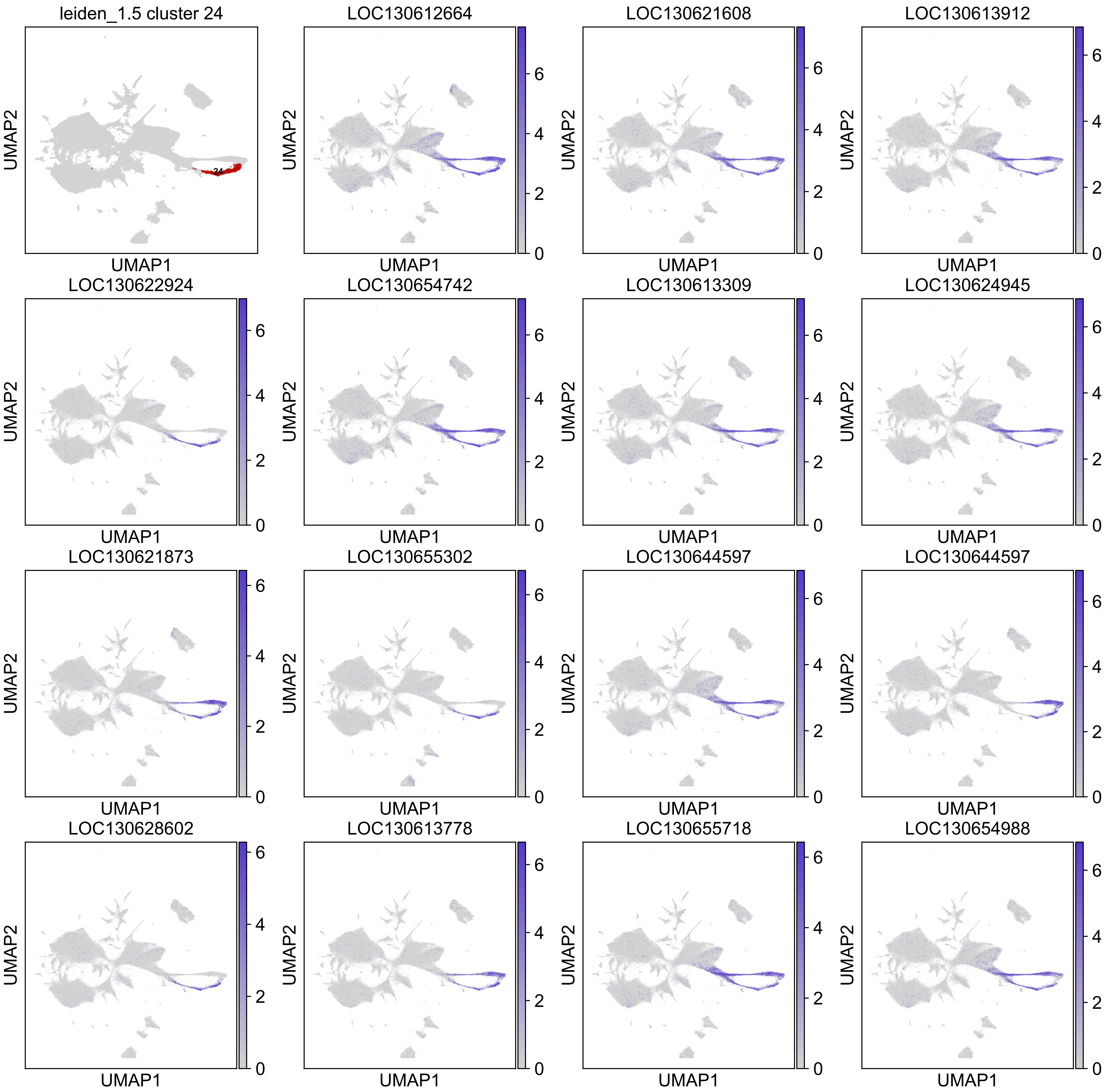
leiden_1.5 cluster 24
24
LOC130612664
LOC130621608
LOC130613912
LOC130622924
LOC130654742
LOC130613309
LOC130624945
LOC130621873
LOC130655302
LOC130644597
LOC130644597
LOC130628602
LOC130613778
LOC130655718
LOC130654988
UMAP1
UMAP2
0
2
4
6

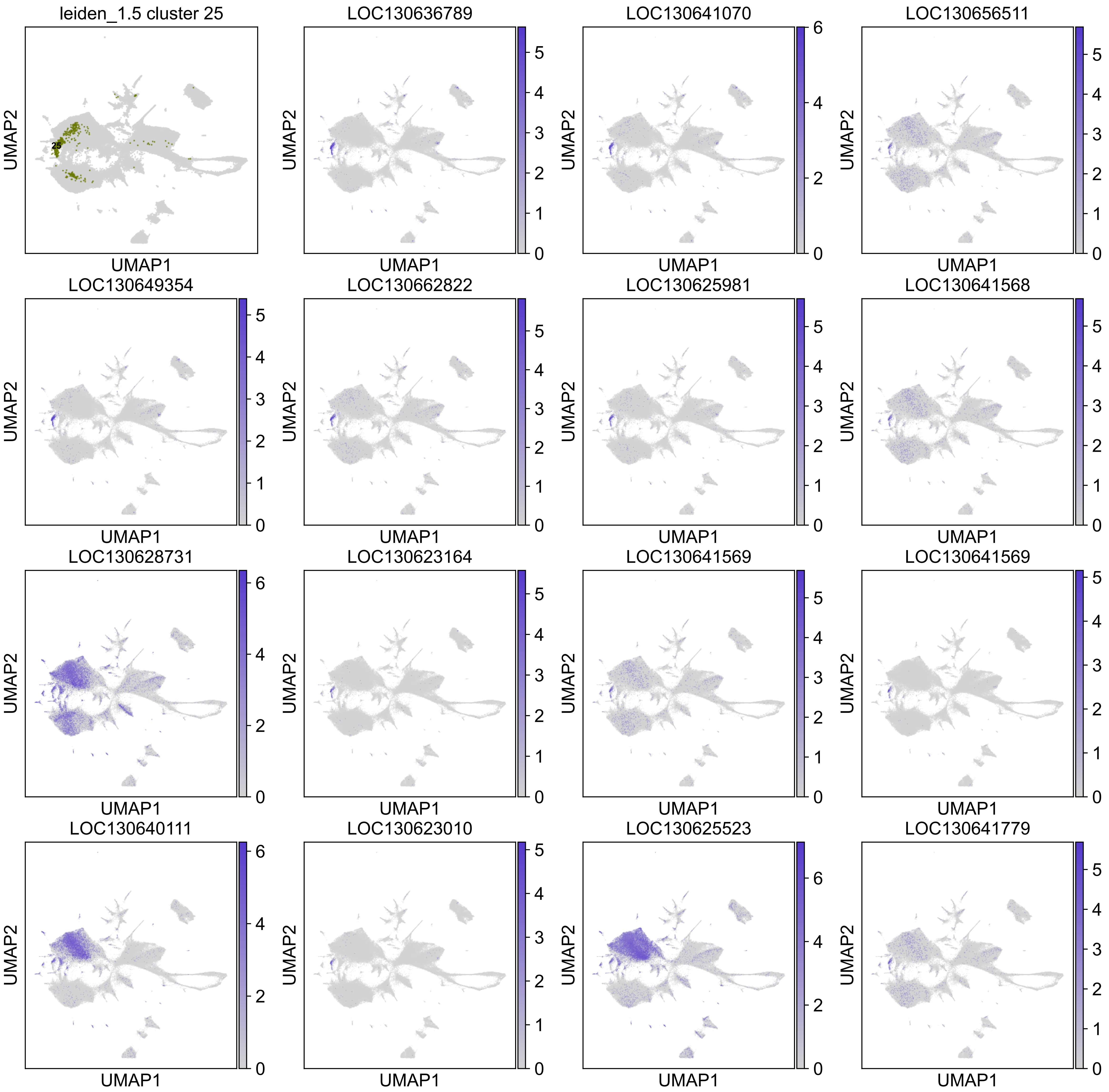
leiden_1.5 cluster 25
LOC130636789
LOC130641070
LOC130656511
LOC130649354
LOC130662822
LOC130625981
LOC130641568
LOC130628731
LOC130623164
LOC130641569
LOC130641569
LOC130640111
LOC130623010
LOC130625523
LOC130641779
UMAP1
UMAP2

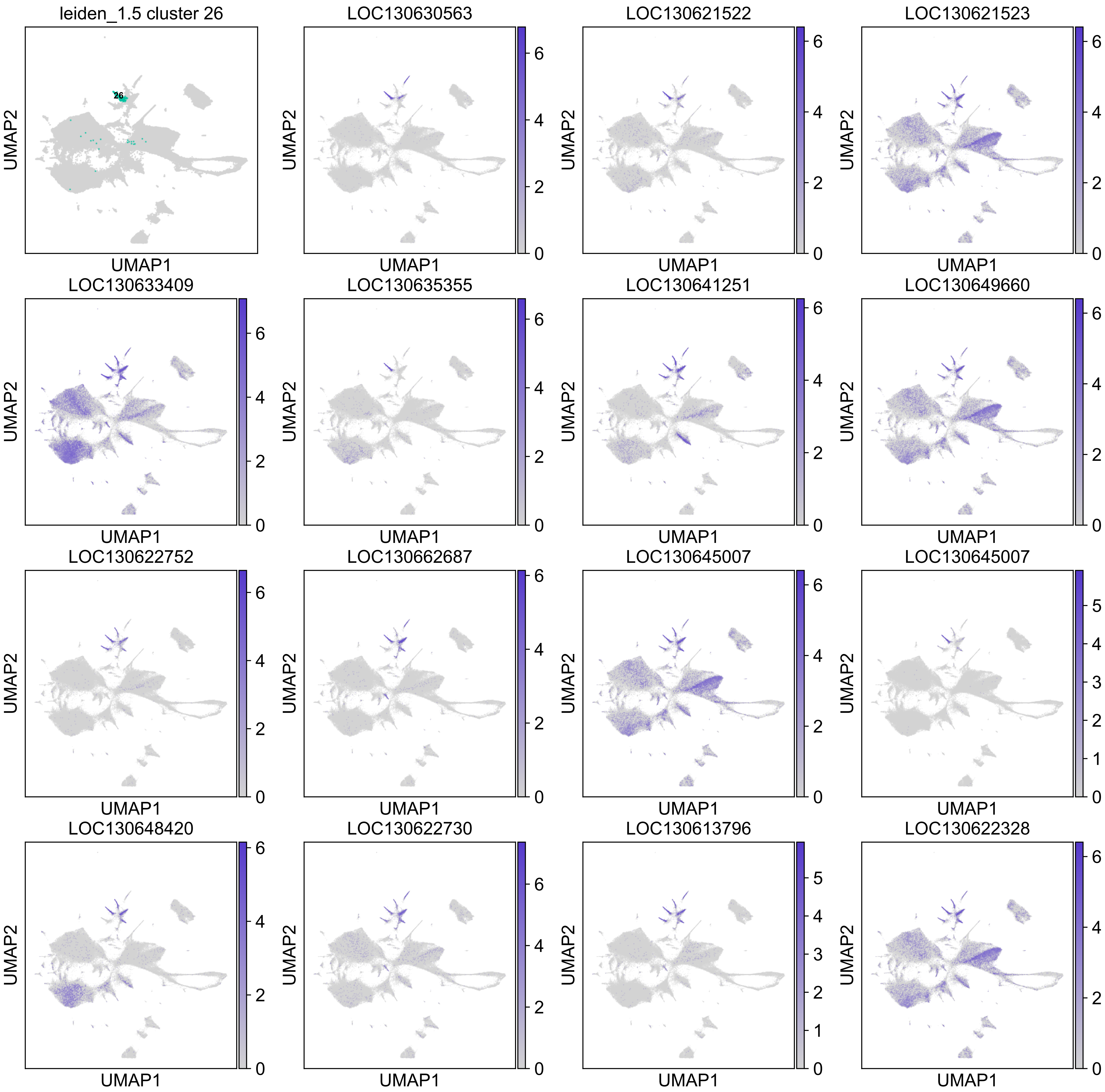

leiden_1.5 cluster 26
LOC130630563
LOC130621522
LOC130621523
LOC130633409
LOC130635355
LOC130641251
LOC130649660
LOC130622752
LOC130662687
LOC130645007
LOC130645007
LOC130648420
LOC130622730
LOC130613796
LOC130622328
UMAP1
UMAP2

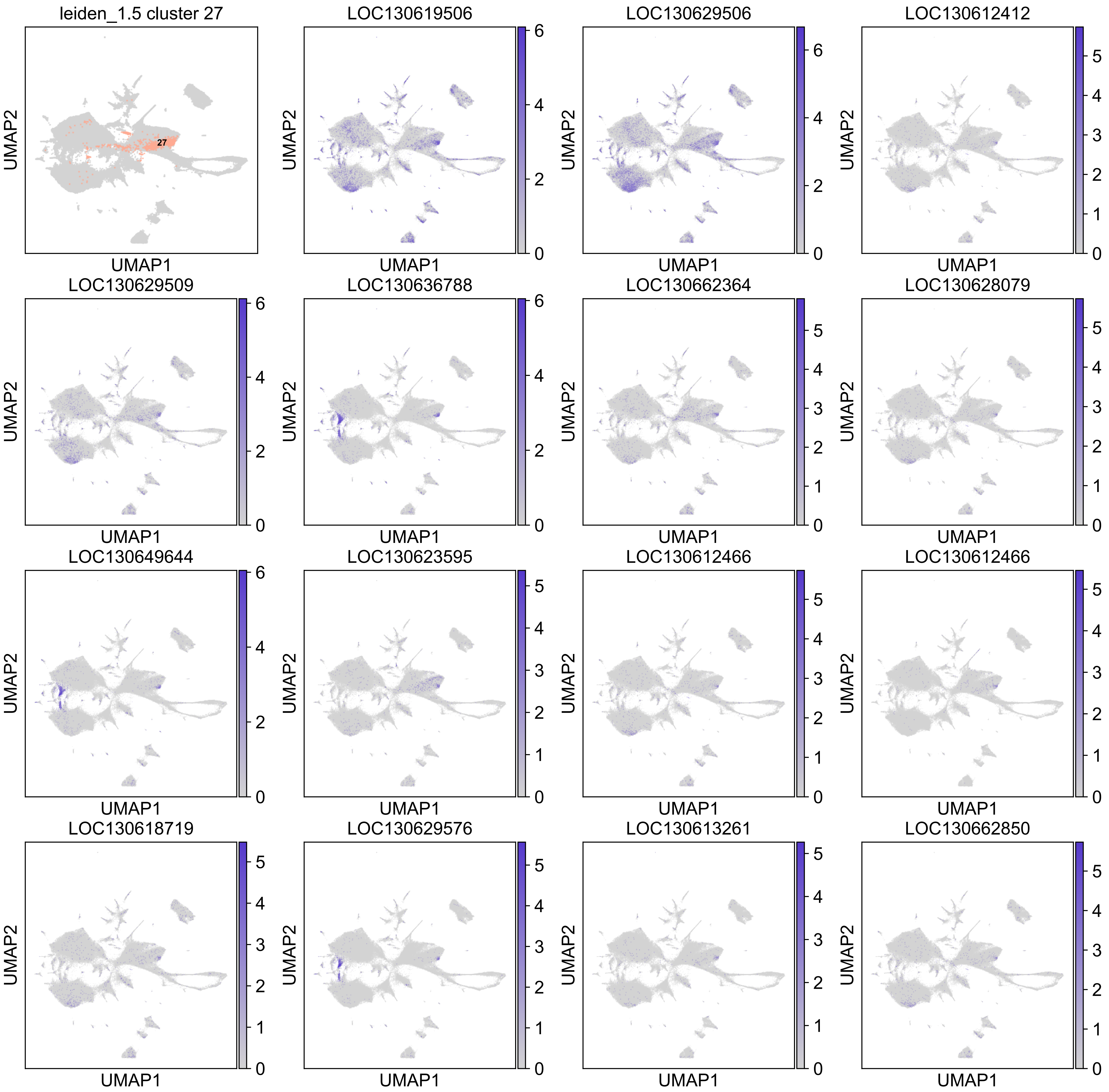
leiden_1.5 cluster 27
27
LOC130619506
LOC130629506
LOC130612412
LOC130629509
LOC130636788
LOC130662364
LOC130628079
LOC130649644
LOC130623595
LOC130612466
LOC130612466
LOC130618719
LOC130629576
LOC130613261
LOC130662850
UMAP1
UMAP2

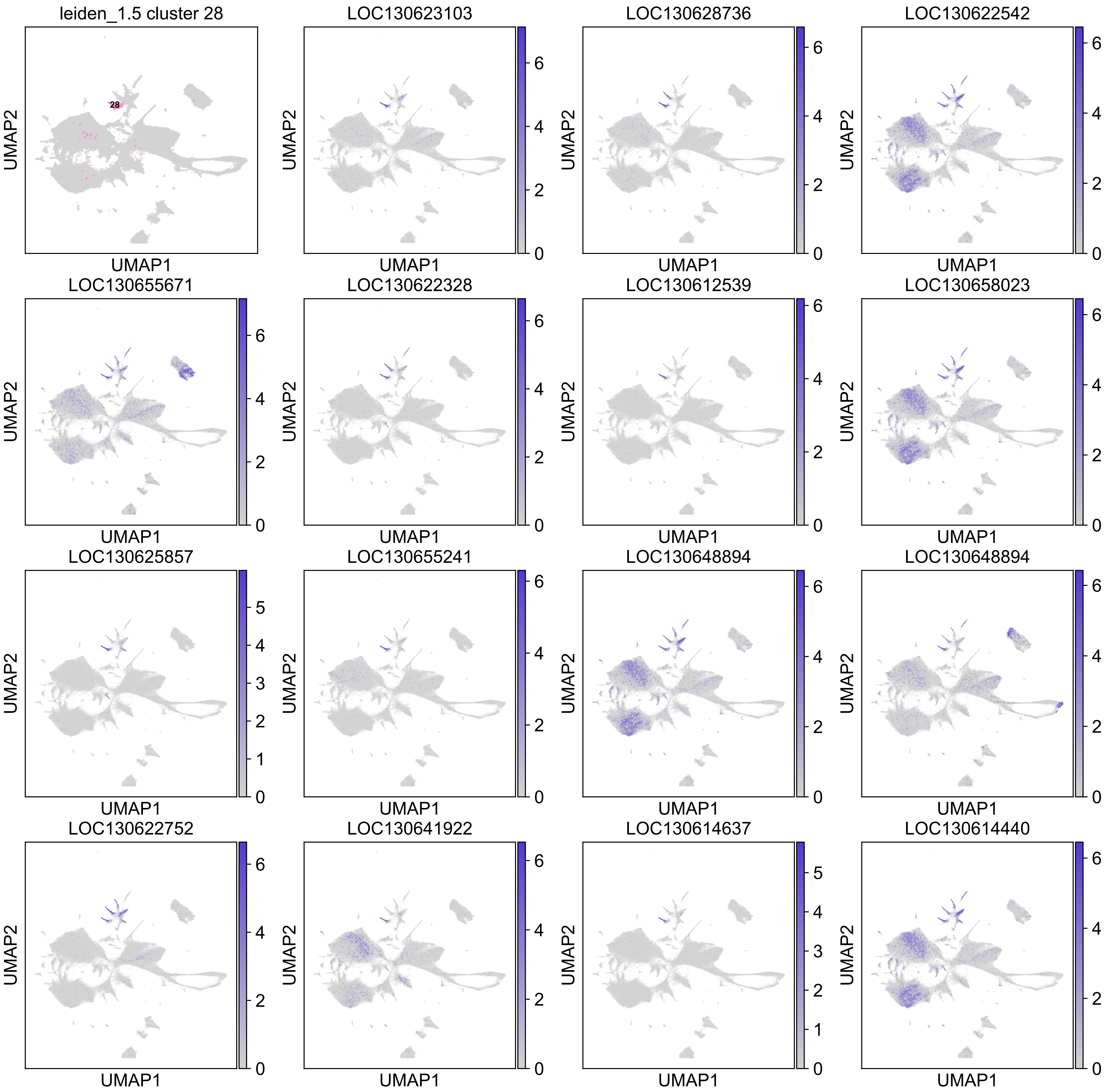

leiden_1.5 cluster 28
28
LOC130623103
LOC130628736
LOC130622542
LOC130655671
LOC130622328
LOC130612539
LOC130658023
LOC130625857
LOC130655241
LOC130648894
LOC130648894
LOC130622752
LOC130641922
LOC130614637
LOC130614440
UMAP1
UMAP2

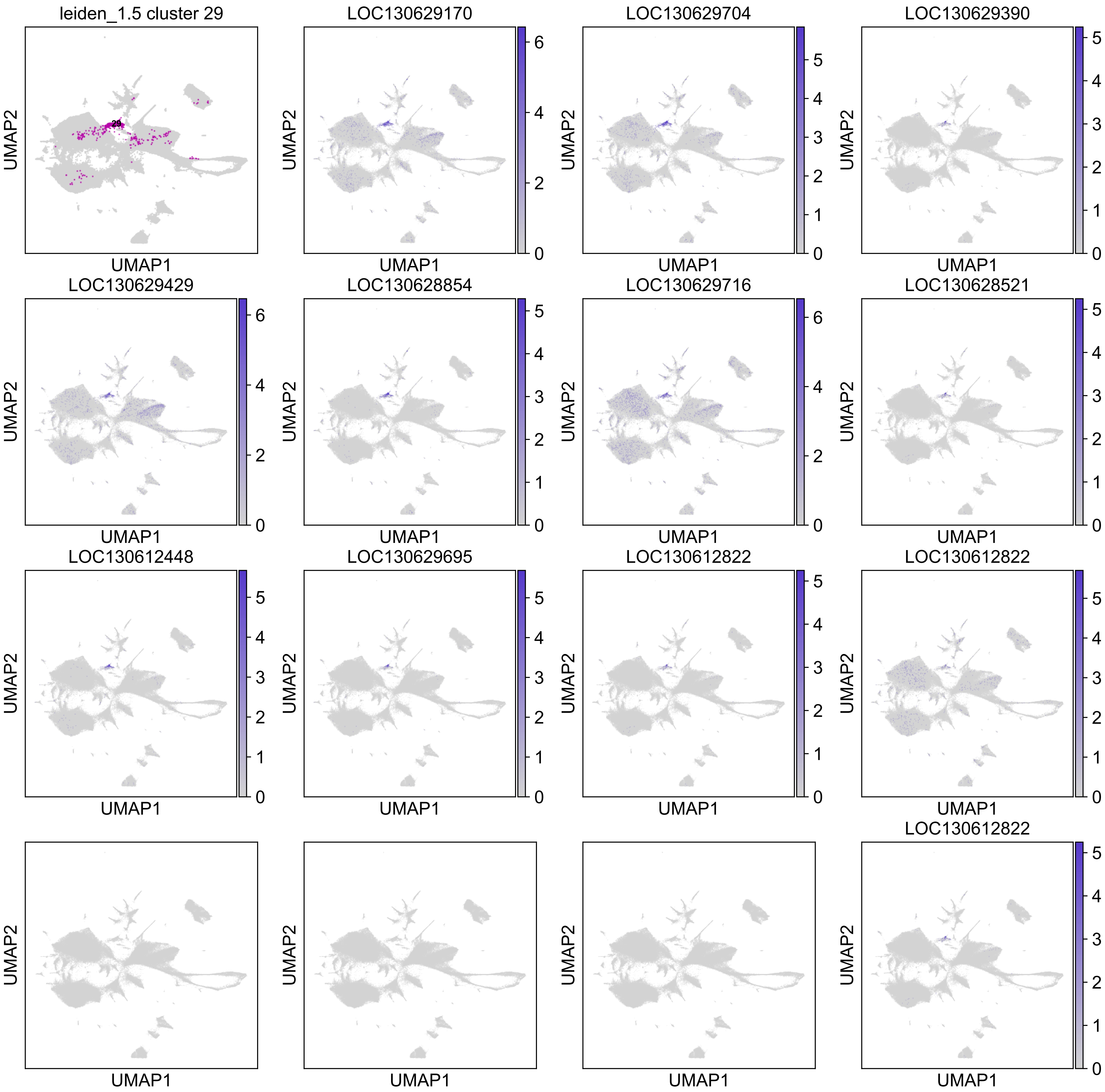
leiden_1.5 cluster 29
29
LOC130629170
LOC130629704
LOC130629390
LOC130629429
LOC130628854
LOC130629716
LOC130628521
LOC130612448
LOC130629695
LOC130612822
LOC130612822
LOC130612822
UMAP1
UMAP2

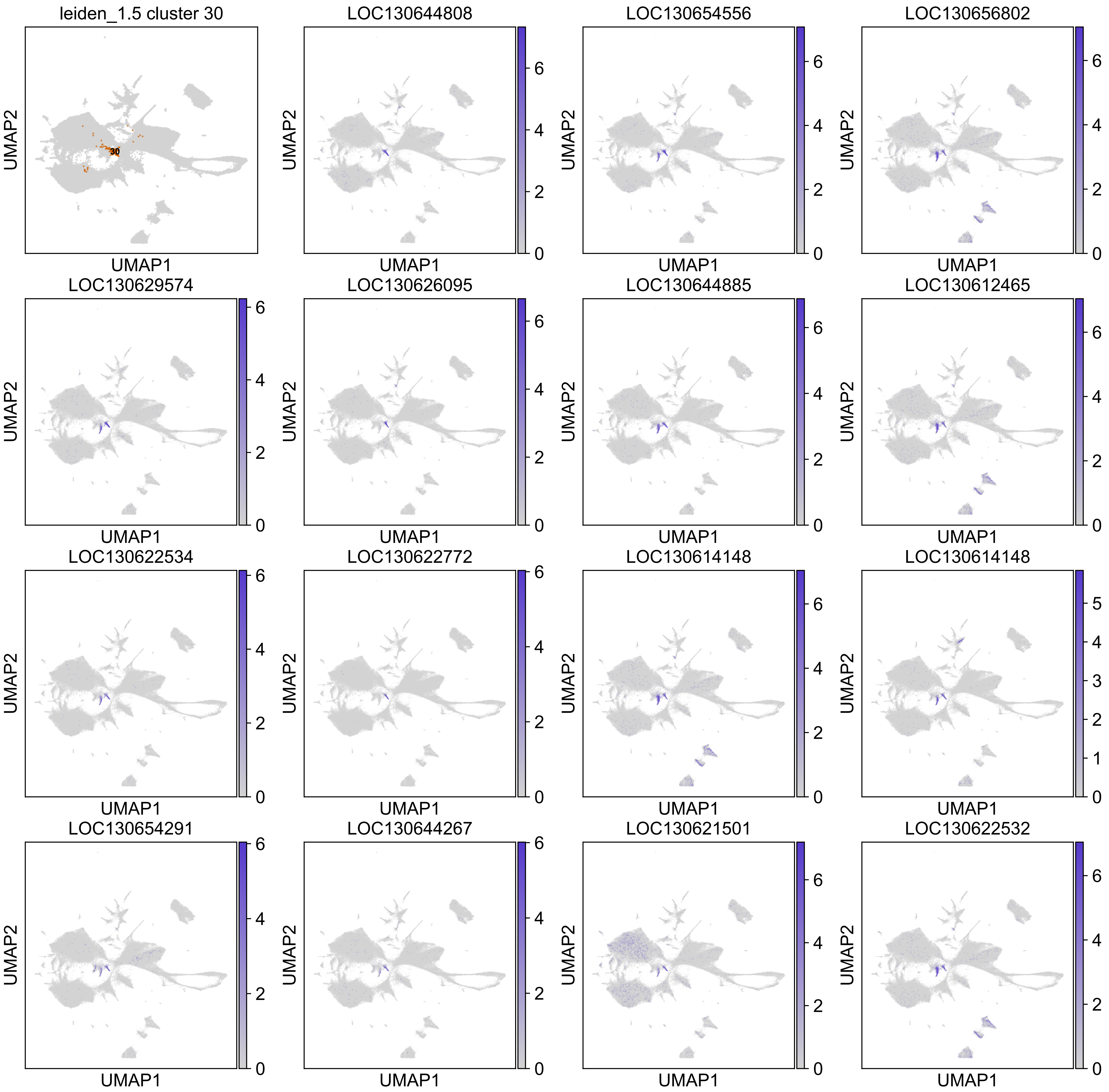
leiden_1.5 cluster 30
30
LOC130644808
LOC130654556
LOC130656802
LOC130629574
LOC130626095
LOC130644885
LOC130612465
LOC130622534
LOC130622772
LOC130614148
LOC130614148
LOC130654291
LOC130644267
LOC130621501
LOC130622532
UMAP1
UMAP2

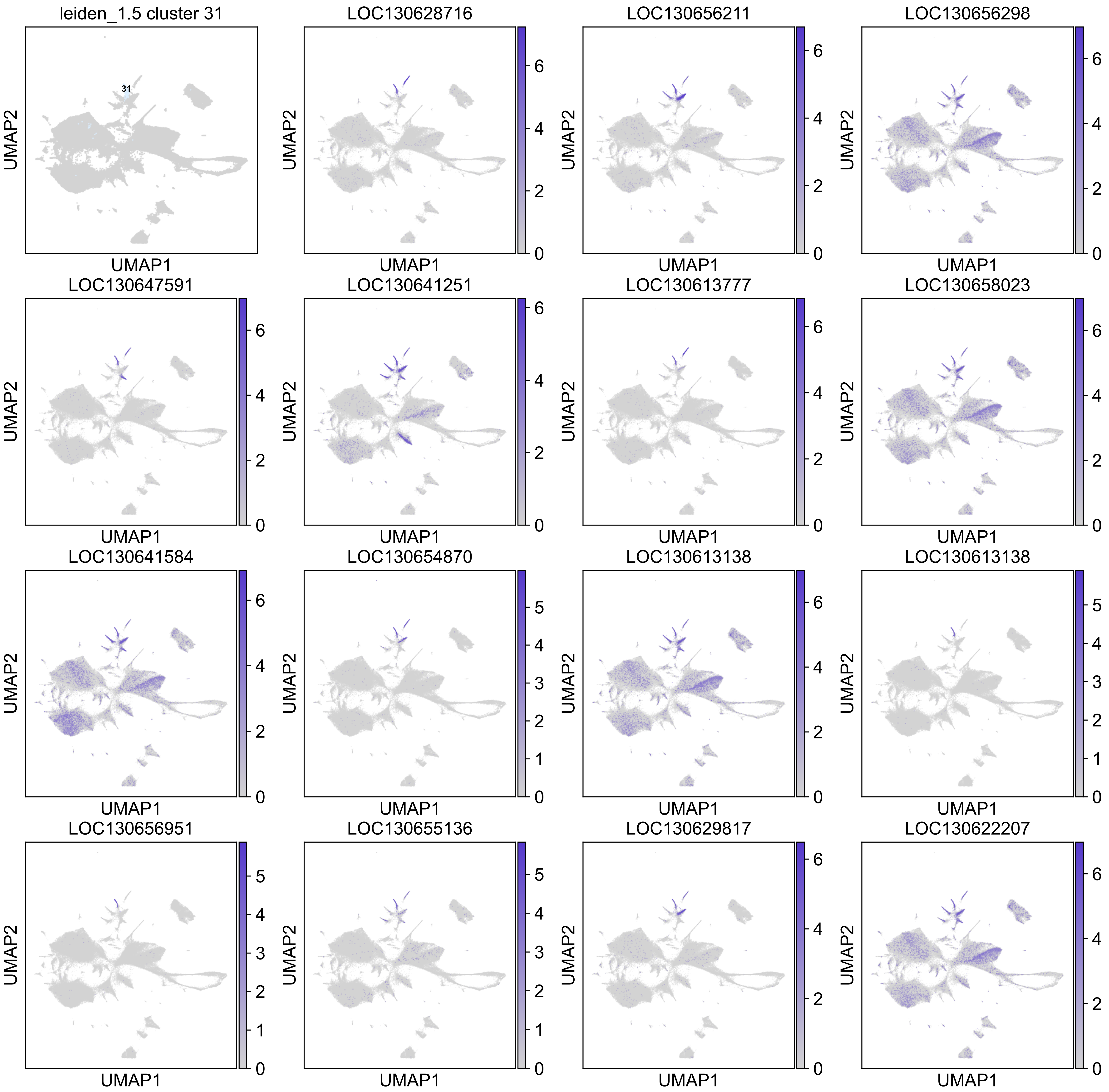
leiden_1.5 cluster 31
31
LOC130628716
LOC130656211
LOC130656298
LOC130647591
LOC130641251
LOC130613777
LOC130658023
LOC130641584
LOC130654870
LOC130613138
LOC130613138
LOC130656951
LOC130655136
LOC130629817
LOC130622207
UMAP1
UMAP2

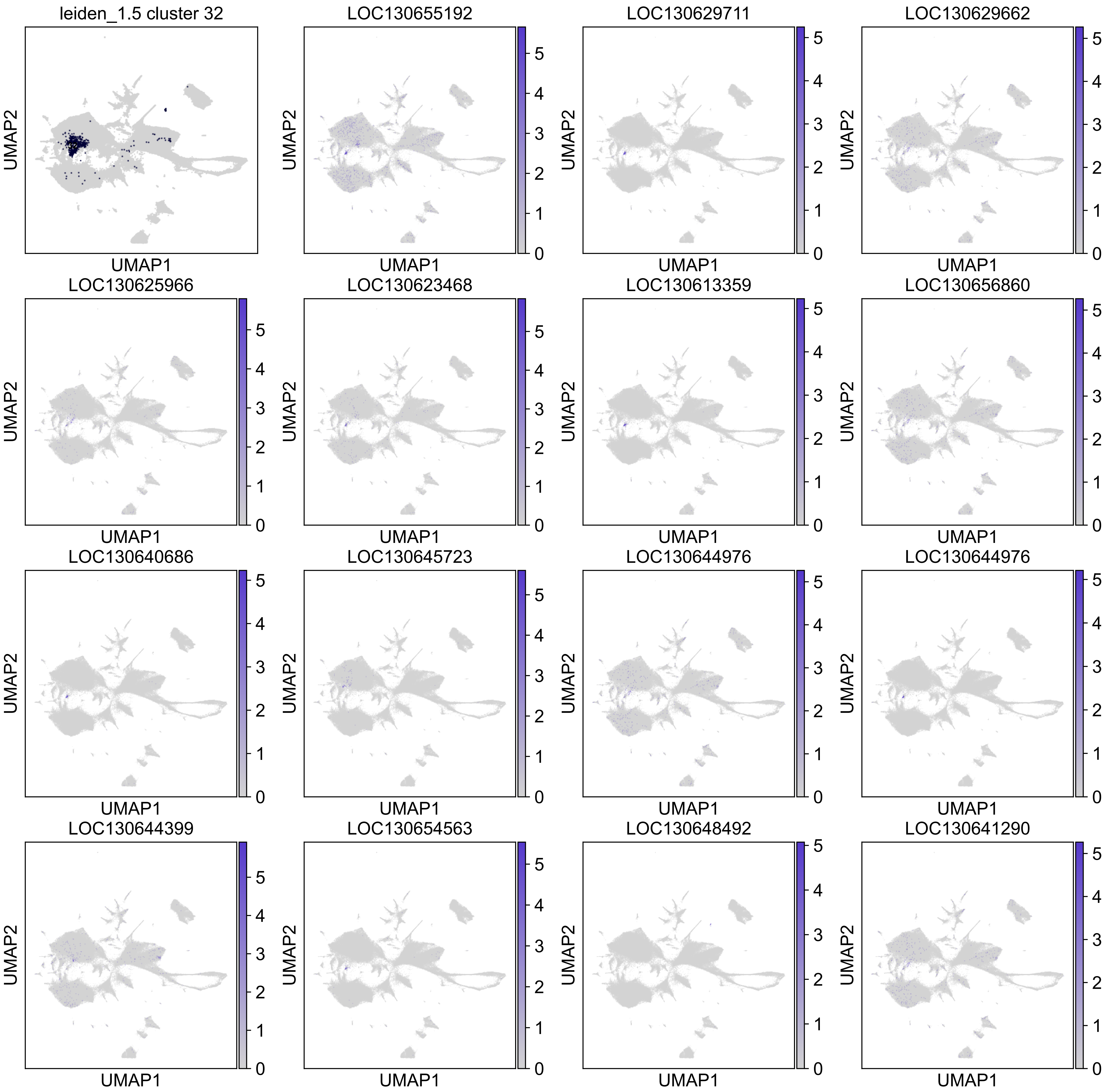
leiden_1.5 cluster 32
LOC130655192
LOC130629711
LOC130629662
LOC130625966
LOC130623468
LOC130613359
LOC130656860
LOC130640686
LOC130645723
LOC130644976
LOC130644976
LOC130644399
LOC130654563
LOC130648492
LOC130641290
UMAP1
UMAP2

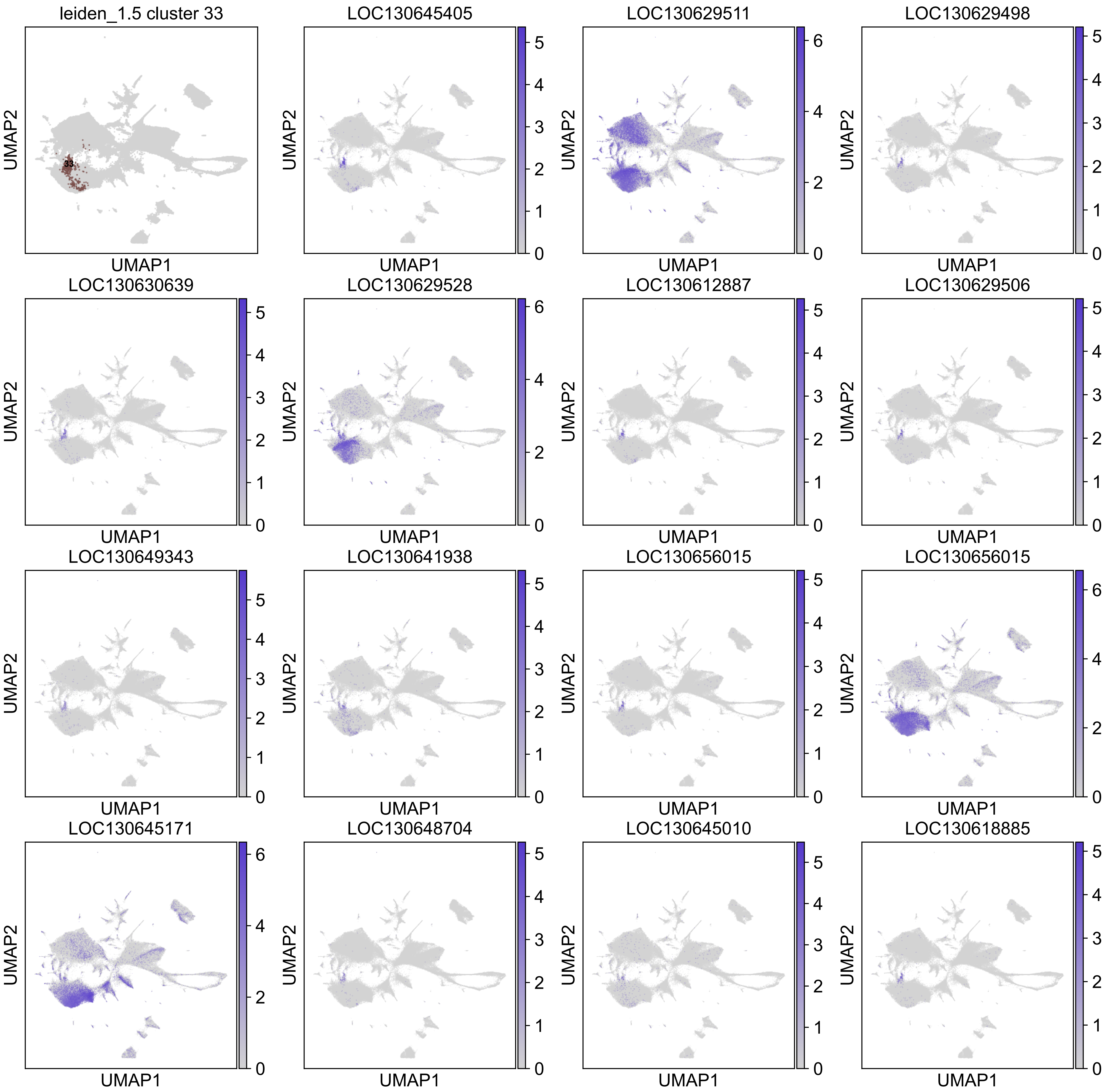

leiden_1.5 cluster 33
33
LOC130645405
LOC130629511
LOC130629498
LOC130630639
LOC130629528
LOC130612887
LOC130629506
LOC130649343
LOC130641938
LOC130656015
LOC130656015
LOC130645171
LOC130648704
LOC130645010
LOC130618885
UMAP1
UMAP2

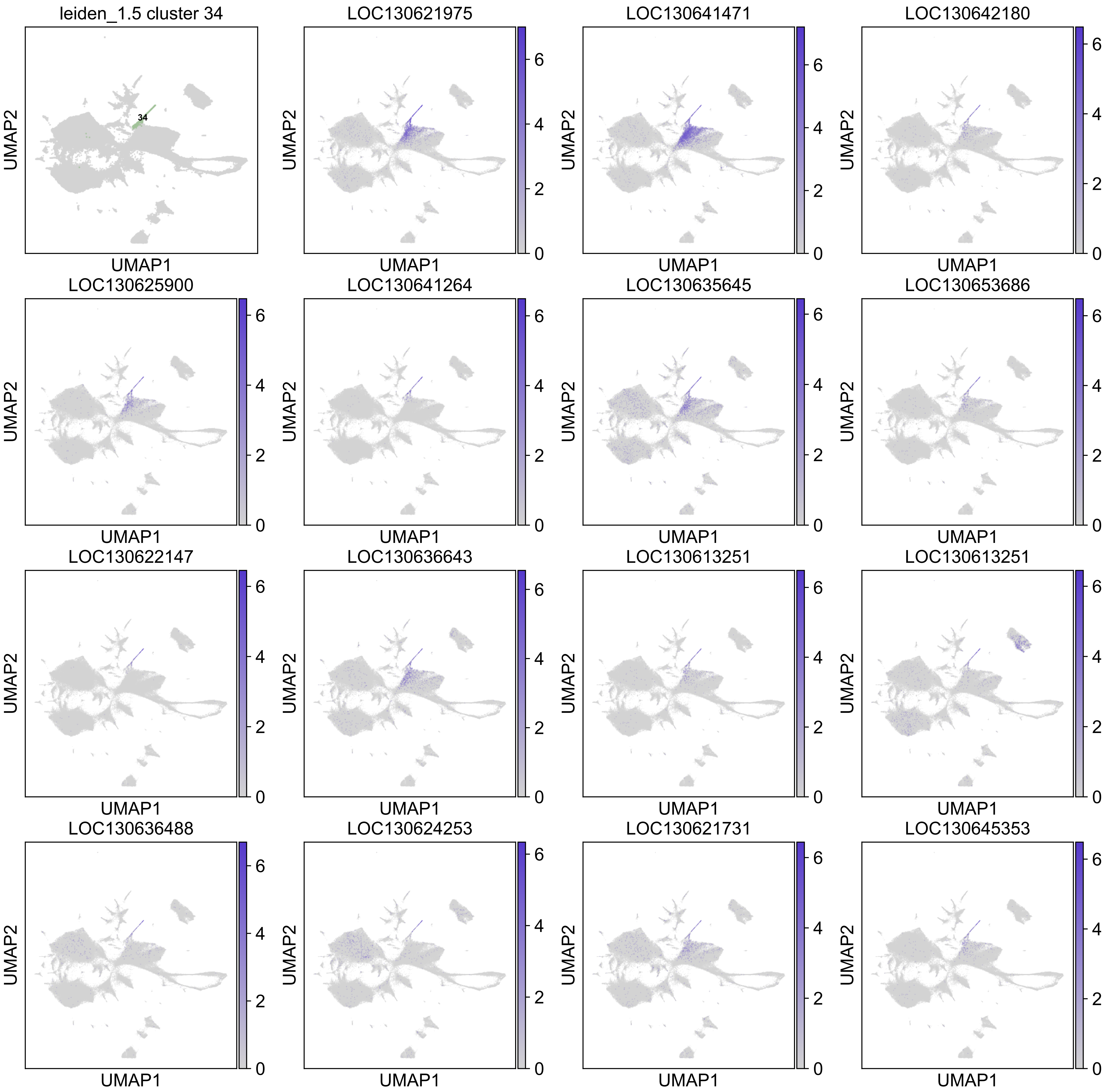

leiden_1.5 cluster 34
34
LOC130621975
LOC130641471
LOC130642180
LOC130625900
LOC130641264
LOC130635645
LOC130653686
LOC130622147
LOC130636643
LOC130613251
LOC130613251
LOC130636488
LOC130624253
LOC130621731
LOC130645353
UMAP1
UMAP2
0
2
4
6

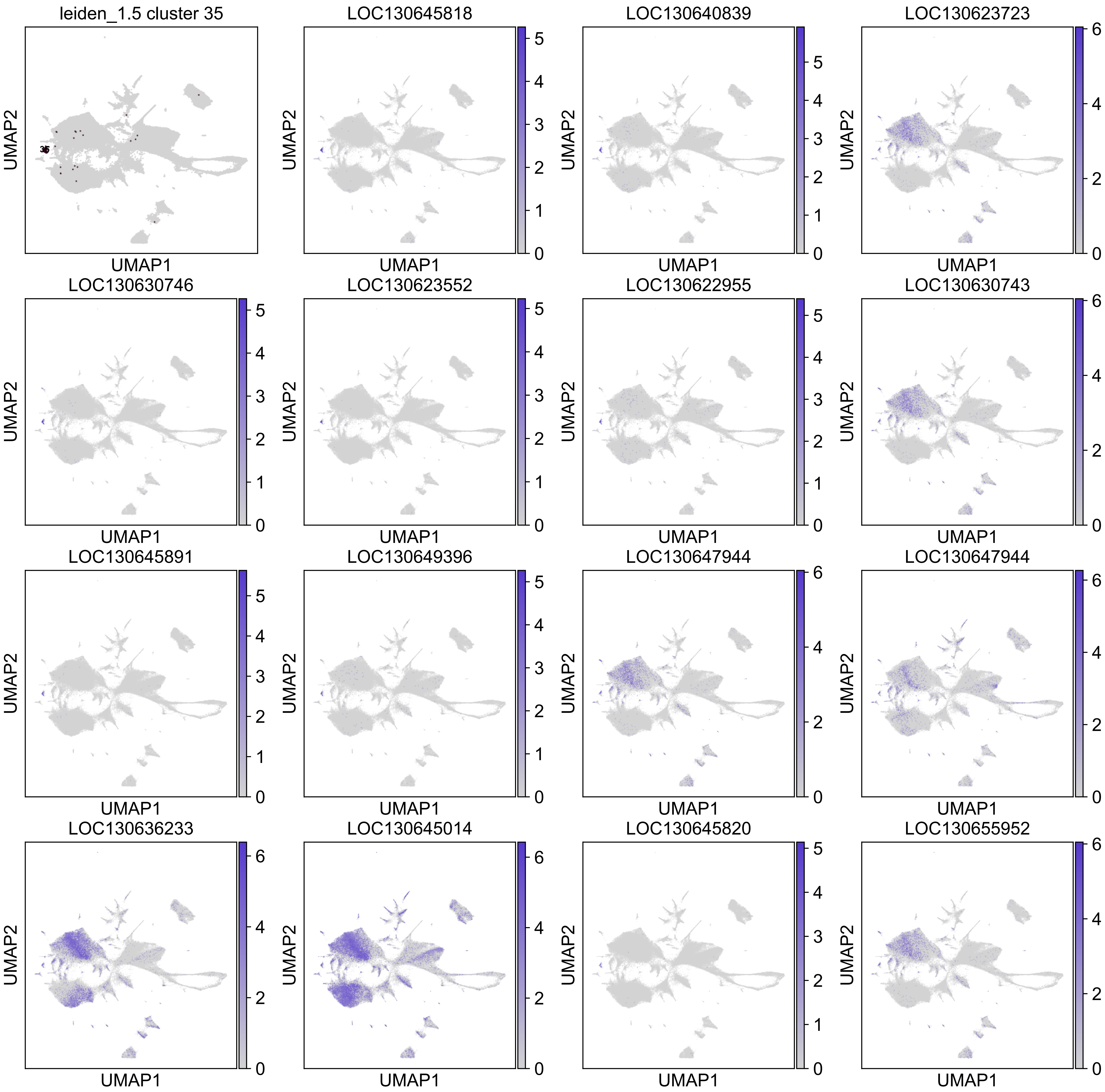
leiden_1.5 cluster 35
LOC130645818
LOC130640839
LOC130623723
LOC130630746
LOC130623552
LOC130622955
LOC130630743
LOC130645891
LOC130649396
LOC130647944
LOC130647944
LOC130636233
LOC130645014
LOC130645820
LOC130655952
UMAP1
UMAP2

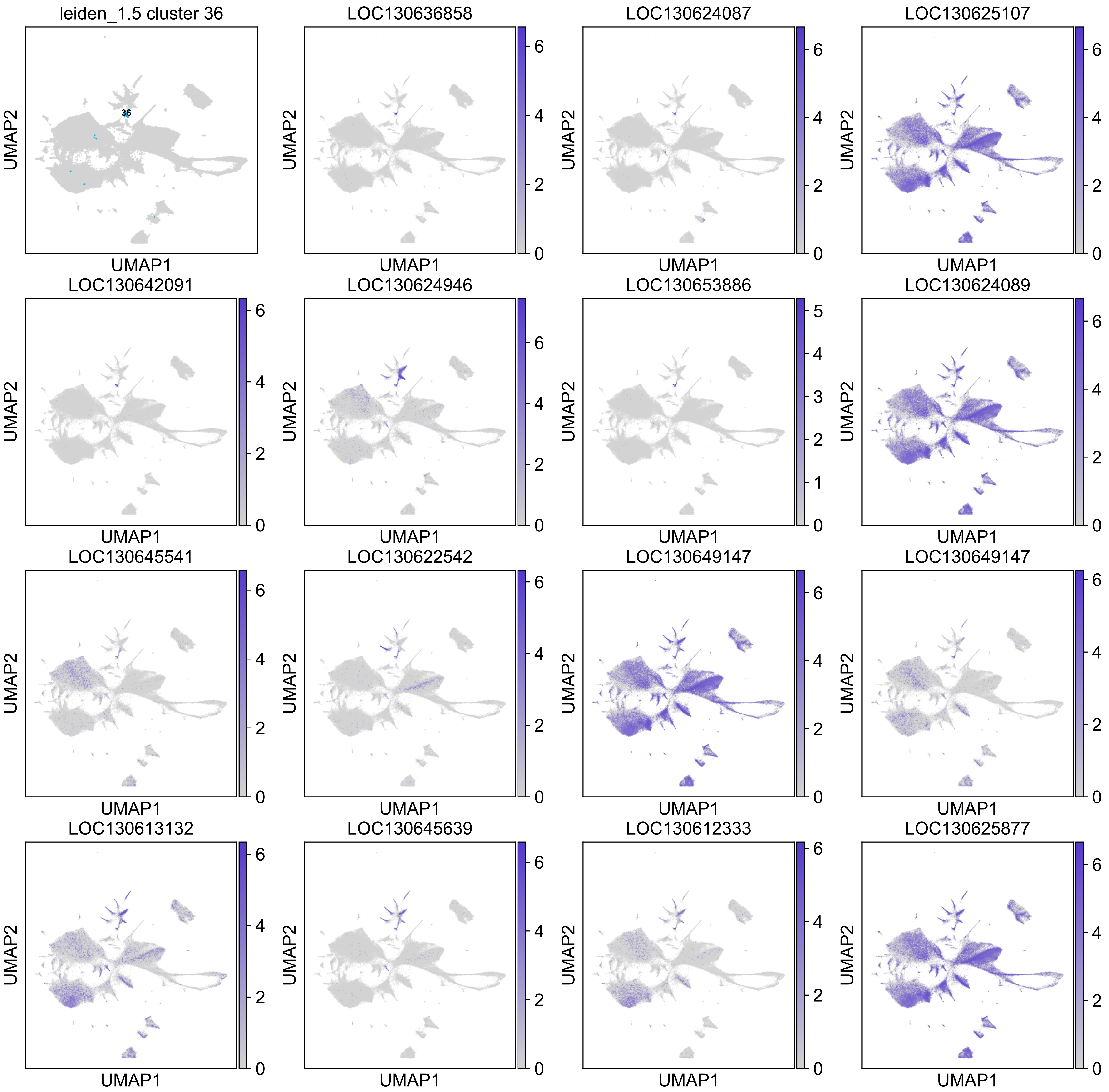
leiden_1.5 cluster 36
36
LOC130636858
LOC130624087
LOC130625107
LOC130642091
LOC130624946
LOC130653886
LOC130624089
LOC130645541
LOC130622542
LOC130649147
LOC130649147
LOC130613132
LOC130645639
LOC130612333
LOC130625877
UMAP1
UMAP2

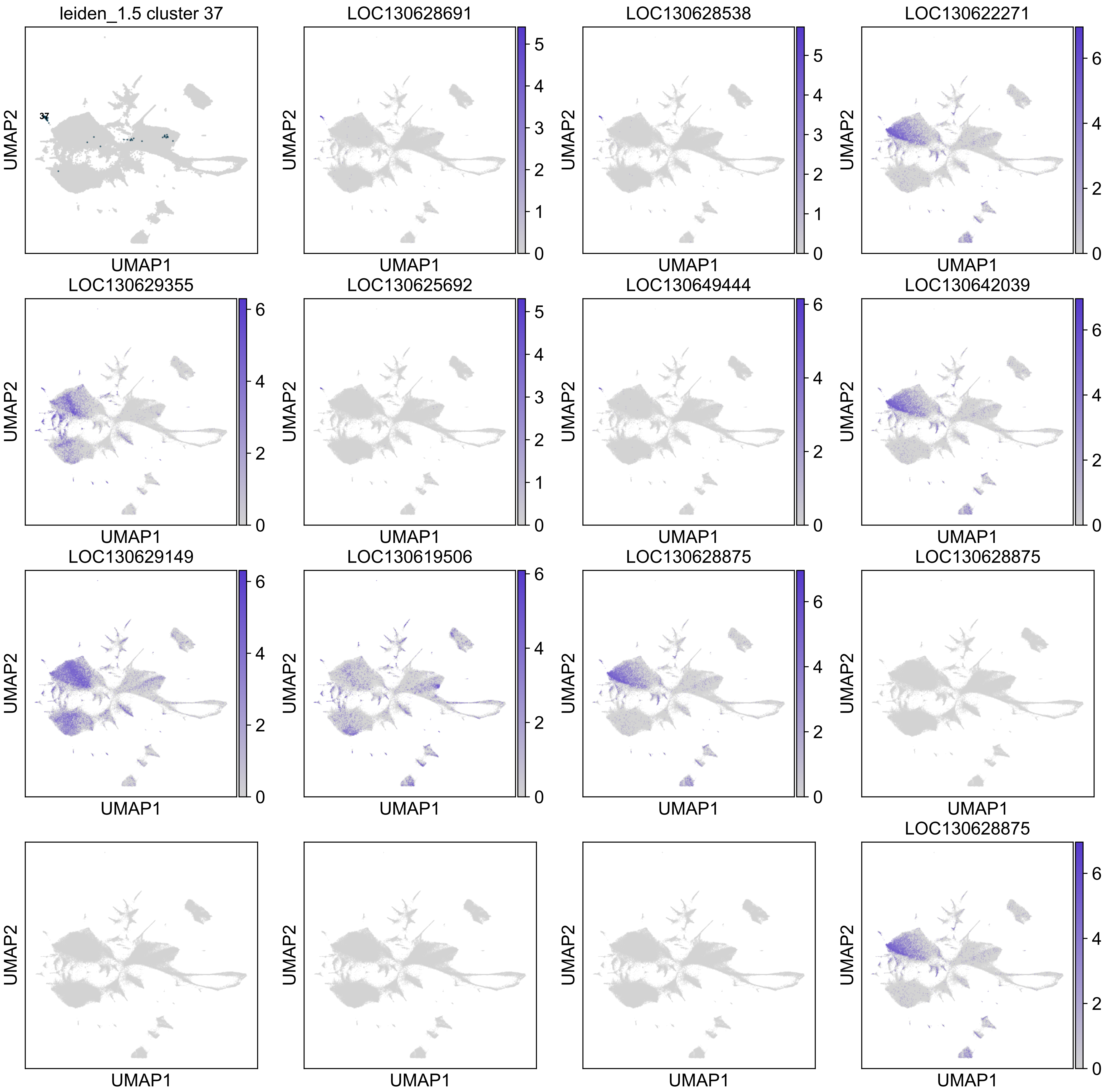

leiden_1.5 cluster 37
37
LOC130628691
LOC130628538
LOC130622271
LOC130629355
LOC130625692
LOC130649444
LOC130642039
LOC130629149
LOC130619506
LOC130628875
LOC130628875
LOC130628875
UMAP1
UMAP2

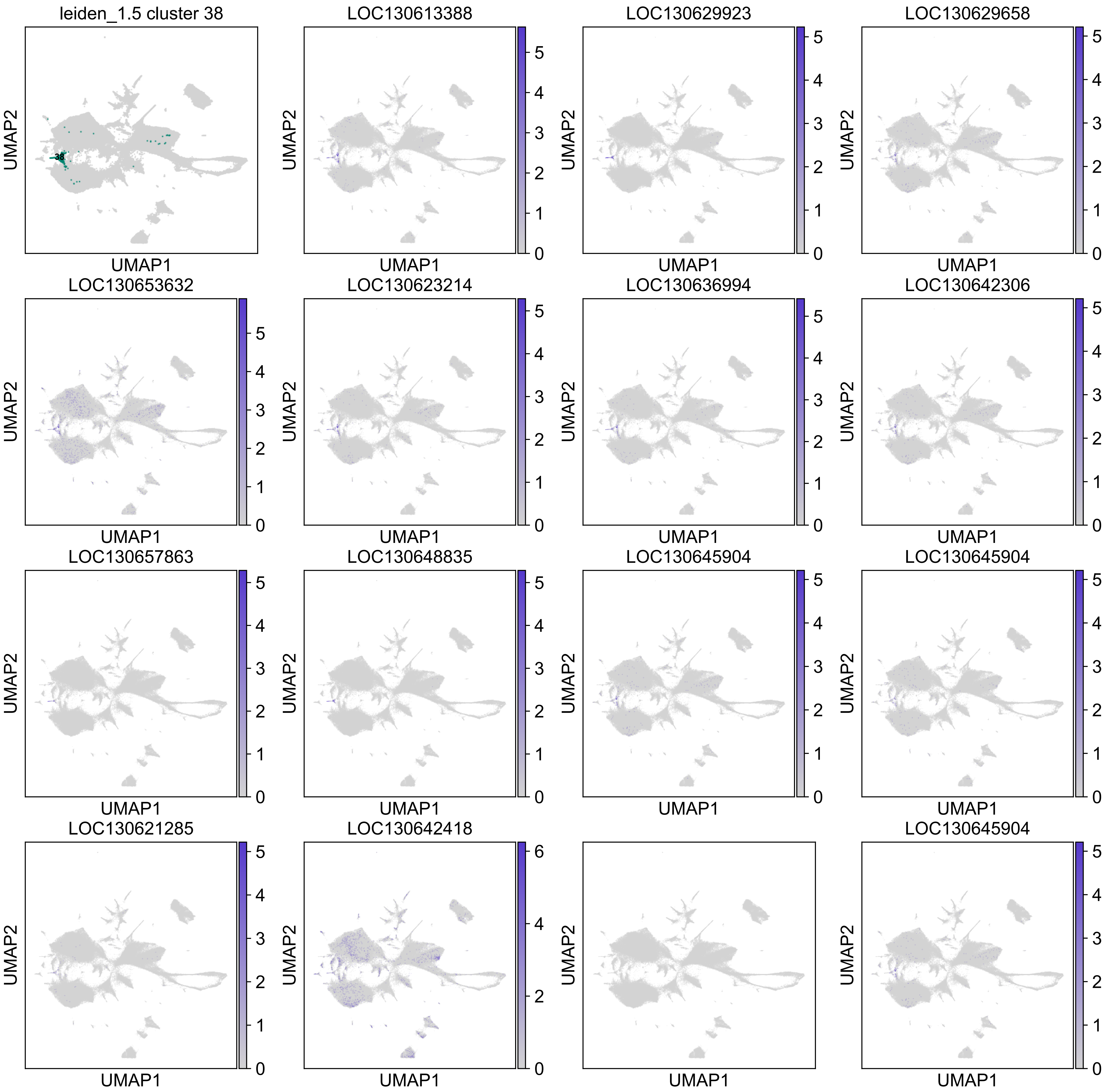

leiden_1.5 cluster 38
38
LOC130613388
LOC130629923
LOC130629658
LOC130653632
LOC130623214
LOC130636994
LOC130642306
LOC130657863
LOC130648835
LOC130645904
LOC130645904
LOC130621285
LOC130642418
LOC130645904
UMAP1
UMAP2

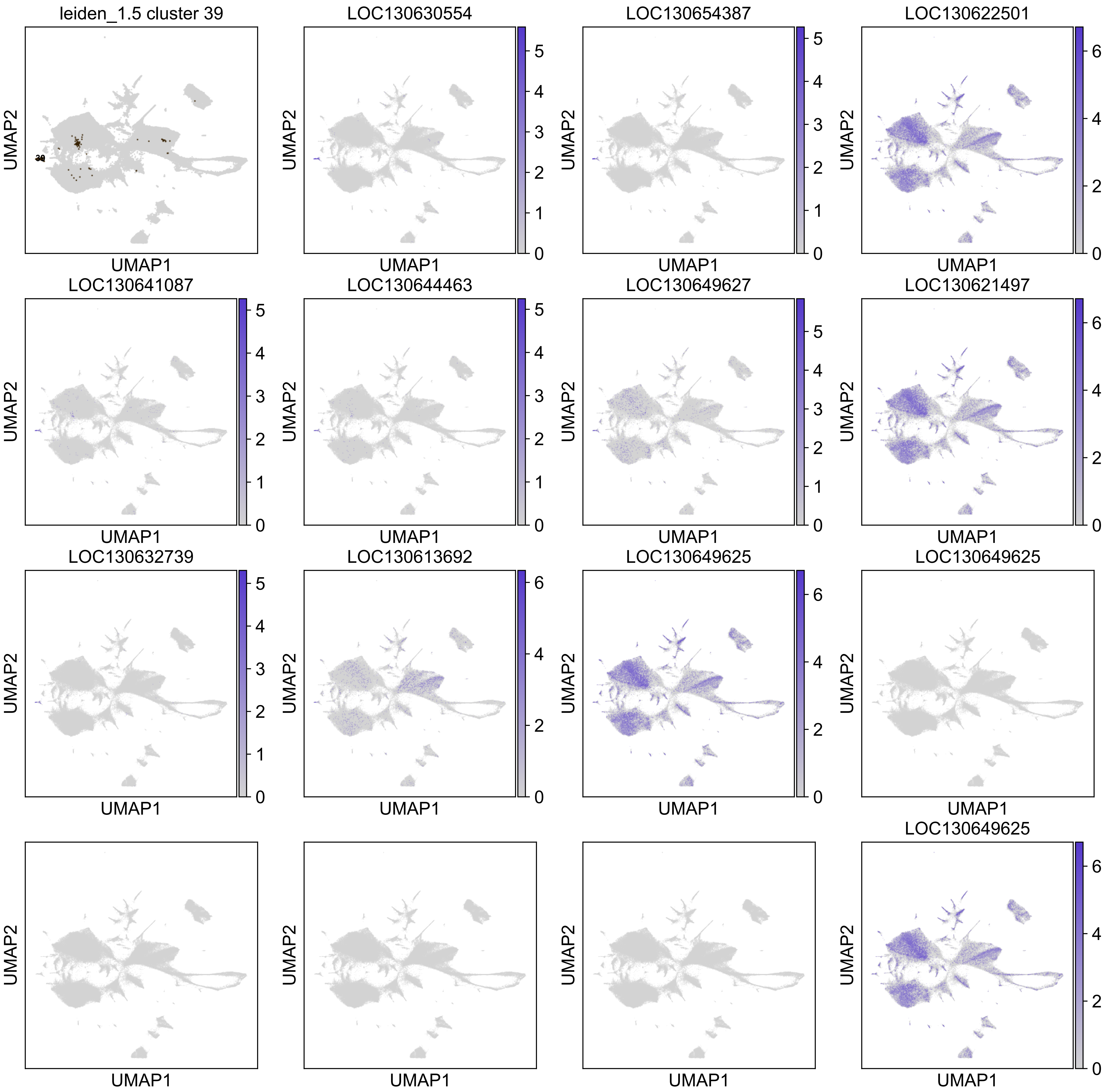

leiden_1.5 cluster 39
LOC130630554
LOC130654387
LOC130622501
LOC130641087
LOC130644463
LOC130649627
LOC130621497
LOC130632739
LOC130613692
LOC130649625
LOC130649625
LOC130649625
UMAP1
UMAP2

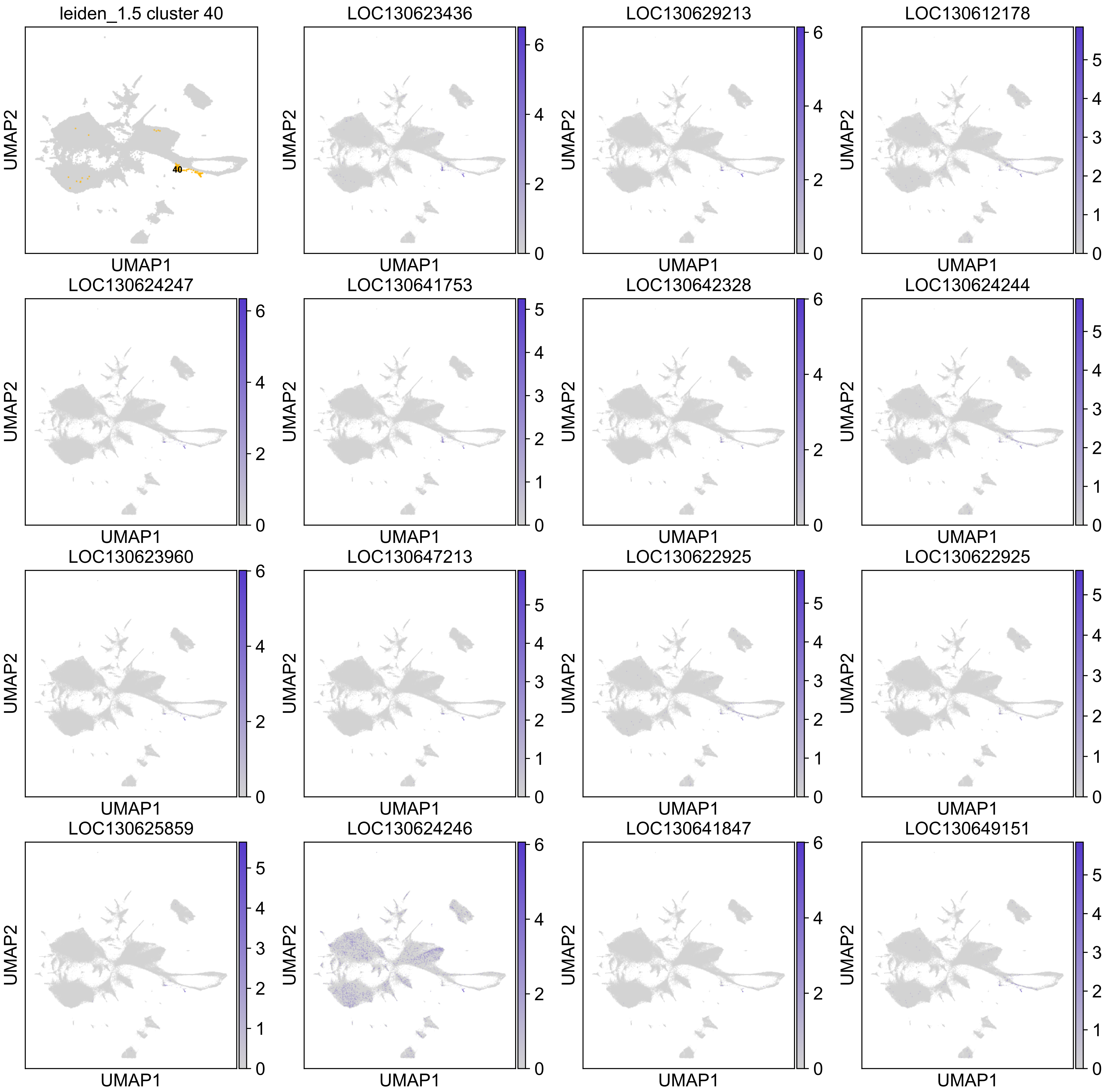

leiden_1.5 cluster 40
40
LOC130623436
LOC130629213
LOC130612178
LOC130624247
LOC130641753
LOC130642328
LOC130624244
LOC130623960
LOC130647213
LOC130622925
LOC130622925
LOC130625859
LOC130624246
LOC130641847
LOC130649151
UMAP1
UMAP2

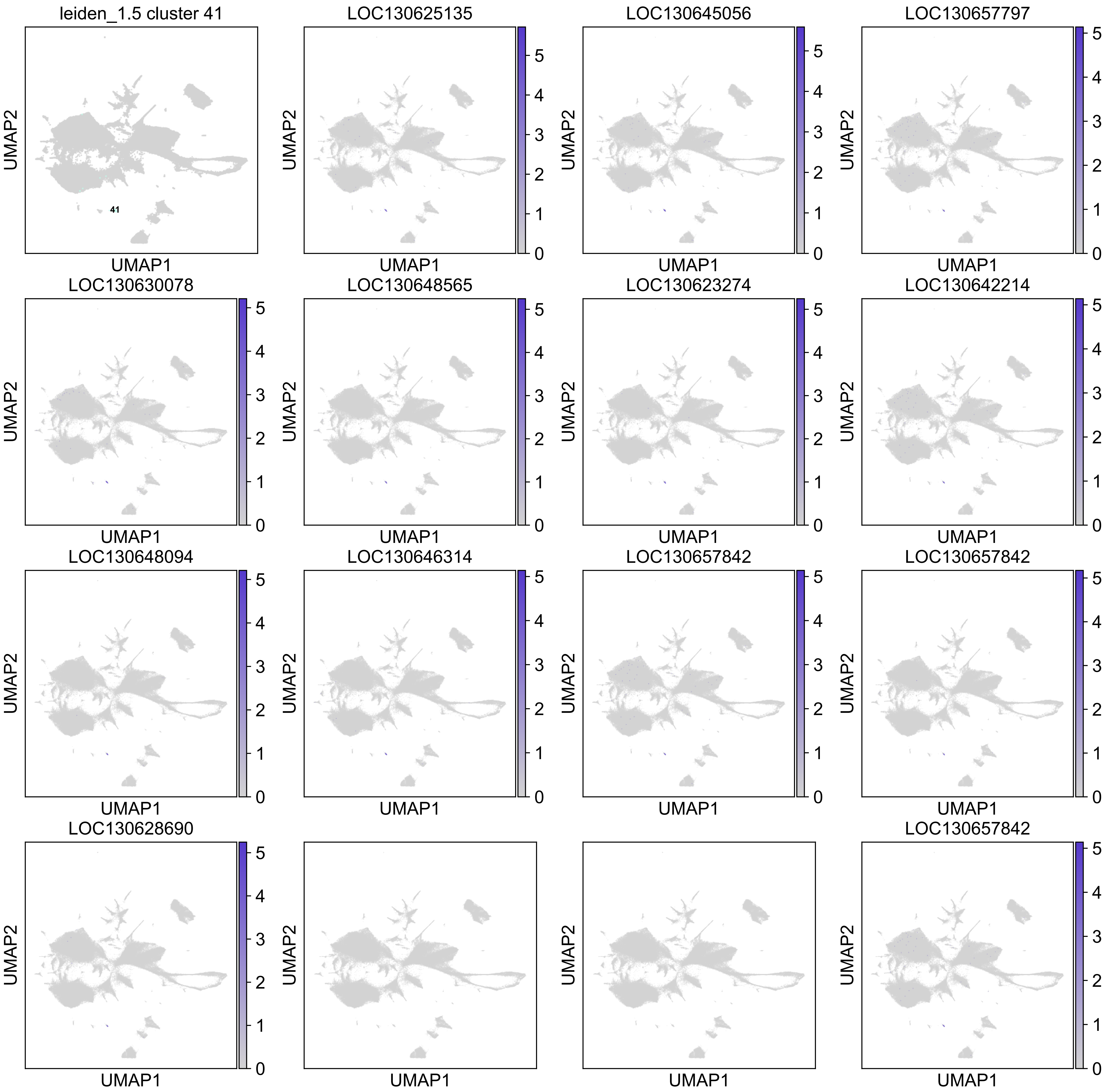
leiden_1.5 cluster 41
41
LOC130625135
LOC130645056
LOC130657797
LOC130630078
LOC130648565
LOC130623274
LOC130642214
LOC130648094
LOC130646314
LOC130657842
LOC130657842
LOC130628690
LOC130657842
UMAP1
UMAP2
0
1
2
3
4
5

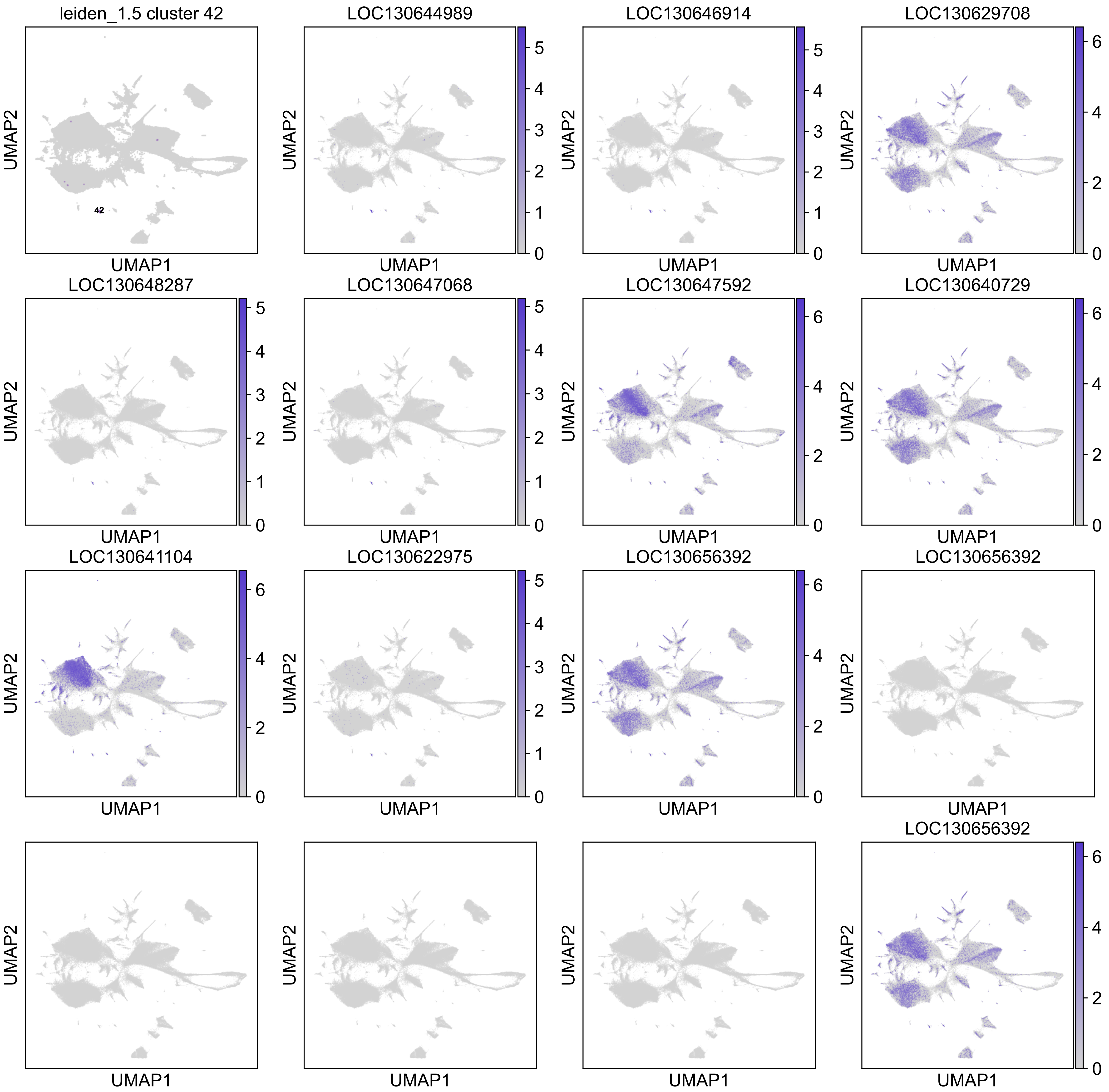

leiden_1.5 cluster 42
42
LOC130644989
LOC130646914
LOC130629708
LOC130648287
LOC130647068
LOC130647592
LOC130640729
LOC130641104
LOC130622975
LOC130656392
LOC130656392
LOC130656392
UMAP1
UMAP2

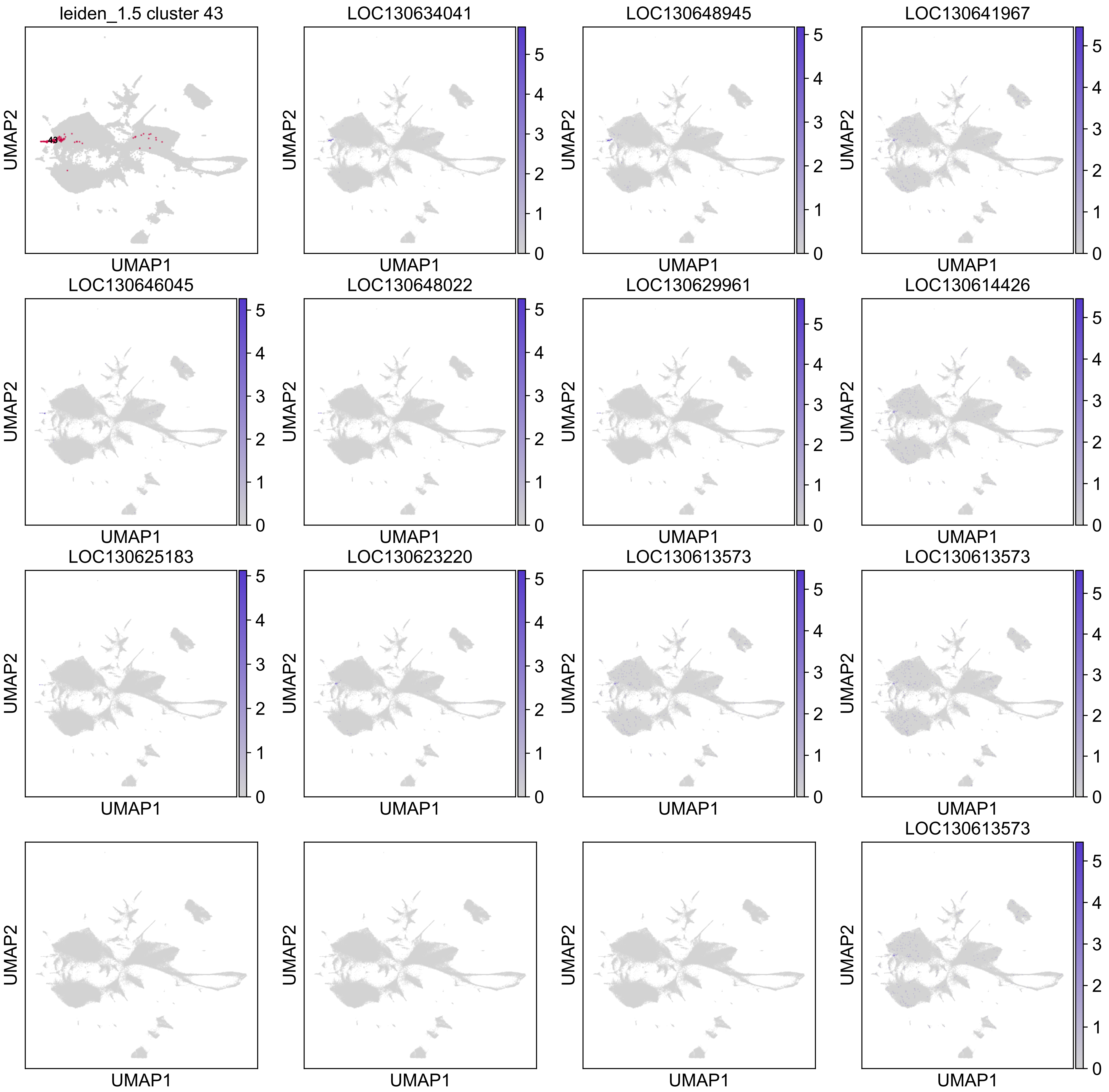
leiden_1.5 cluster 43
LOC130634041
LOC130648945
LOC130641967
LOC130646045
LOC130648022
LOC130629961
LOC130614426
LOC130625183
LOC130623220
LOC130613573
LOC130613573
LOC130613573
UMAP1
UMAP2

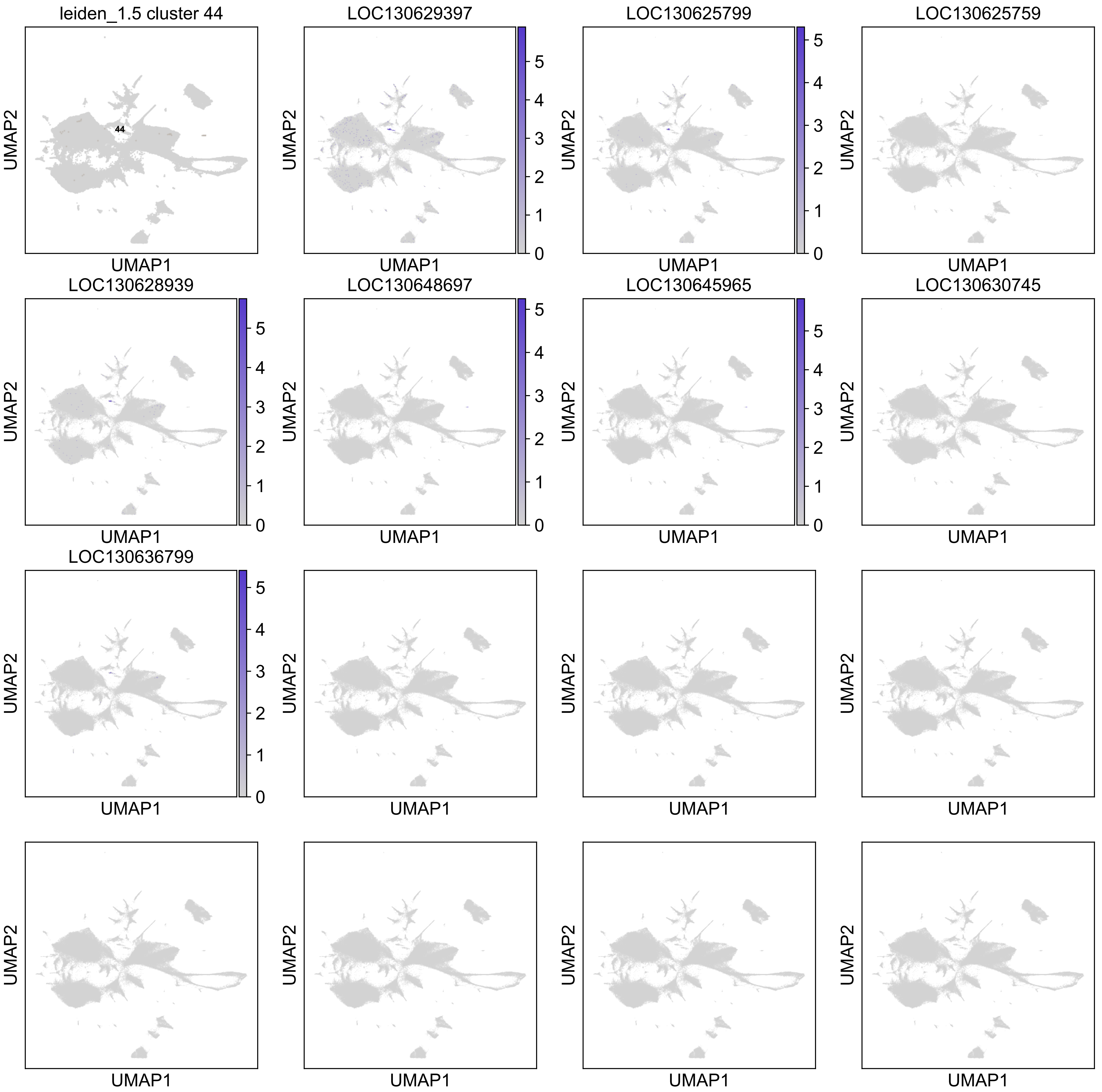
leiden_1.5 cluster 44
44
LOC130629397
LOC130625799
LOC130625759
LOC130628939
LOC130648697
LOC130645965
LOC130630745
LOC130636799
UMAP1
UMAP2

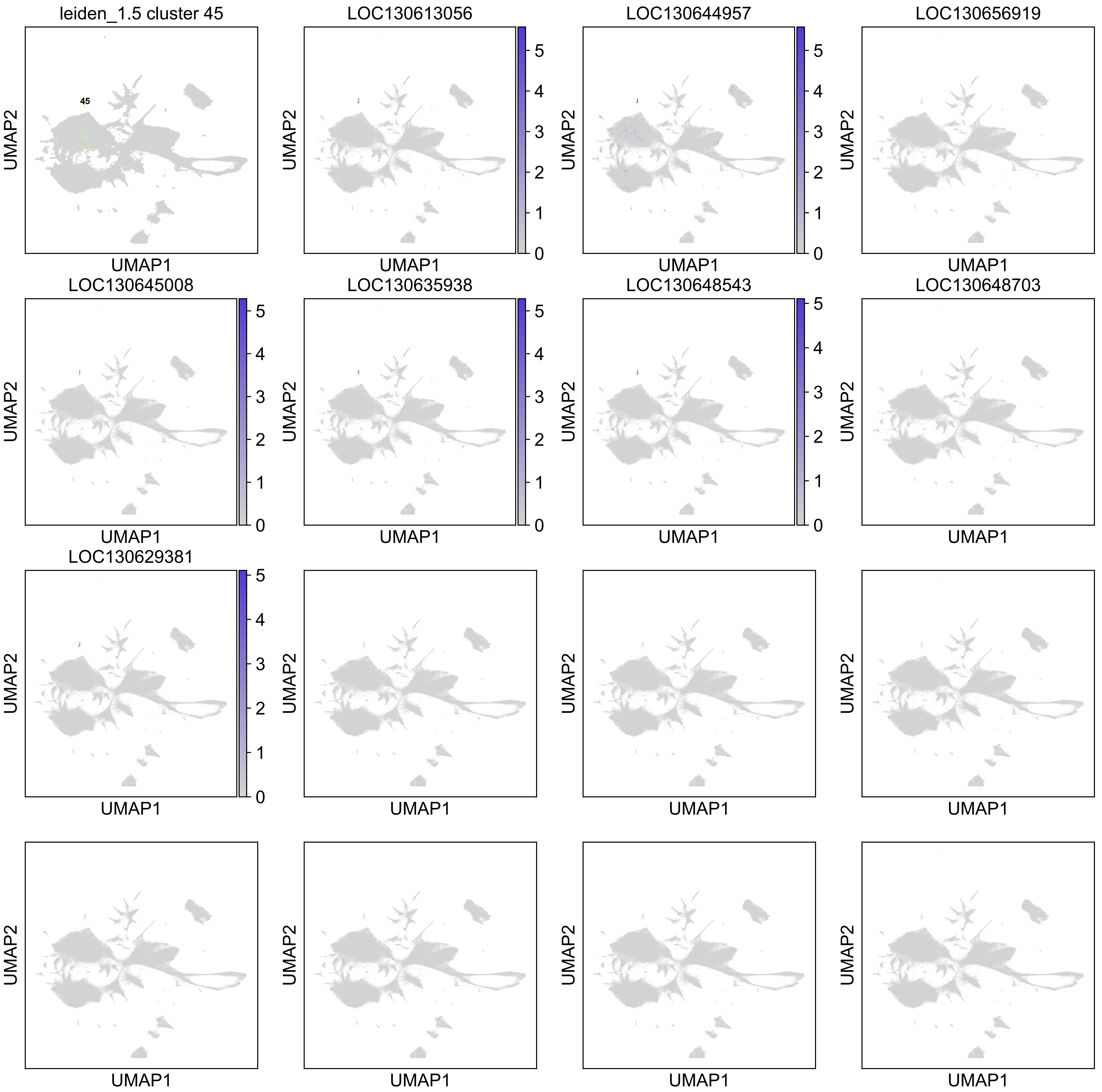
leiden_1.5 cluster 45
45
LOC130613056
LOC130644957
LOC130656919
LOC130645008
LOC130635938
LOC130648543
LOC130648703
LOC130629381
UMAP1
UMAP2
0
1
2
3
4
5

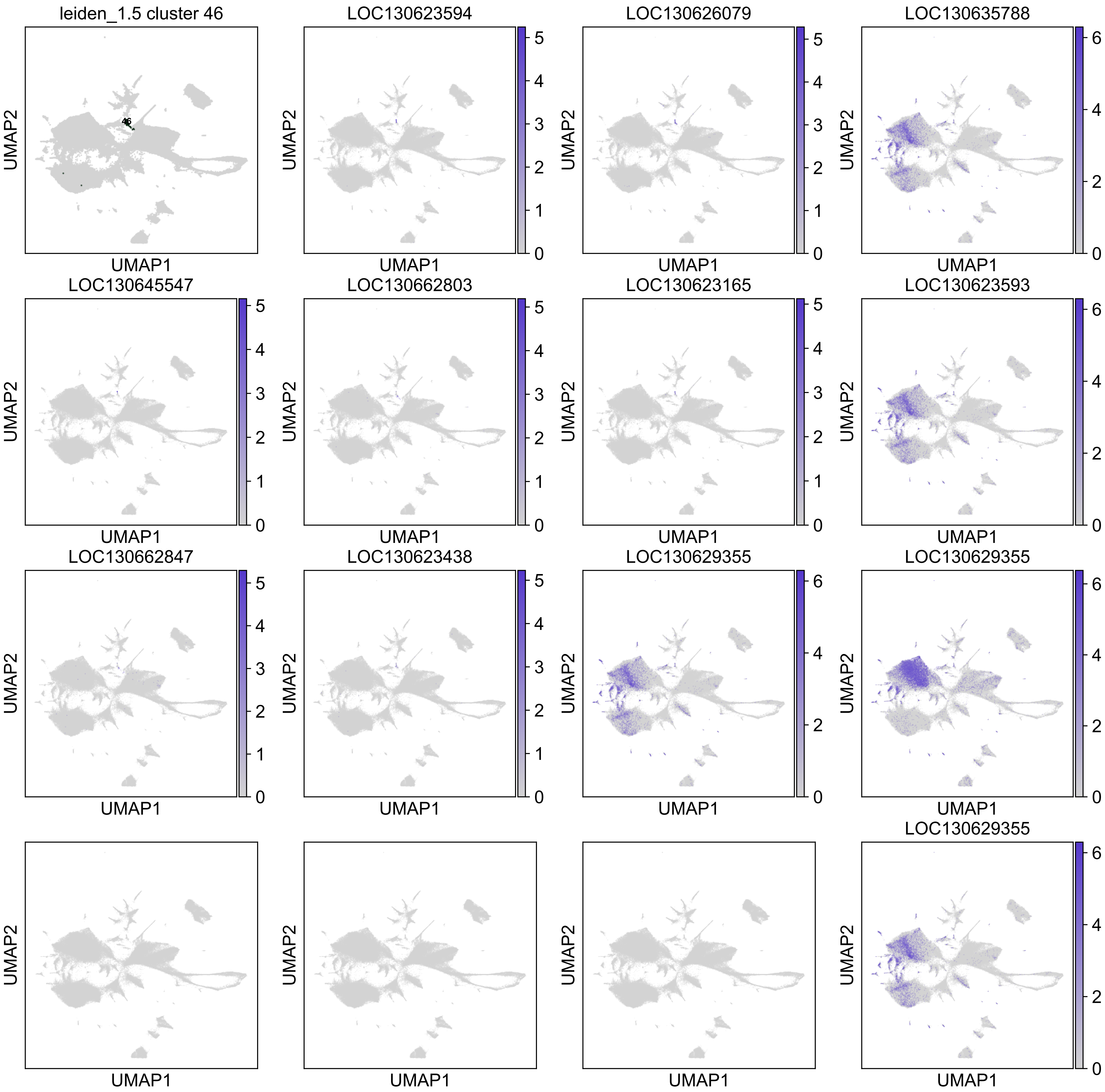
leiden_1.5 cluster 46
46
LOC130623594
LOC130626079
LOC130635788
LOC130645547
LOC130662803
LOC130623165
LOC130623593
LOC130662847
LOC130623438
LOC130629355
LOC130629355
LOC130629355
UMAP1
UMAP2

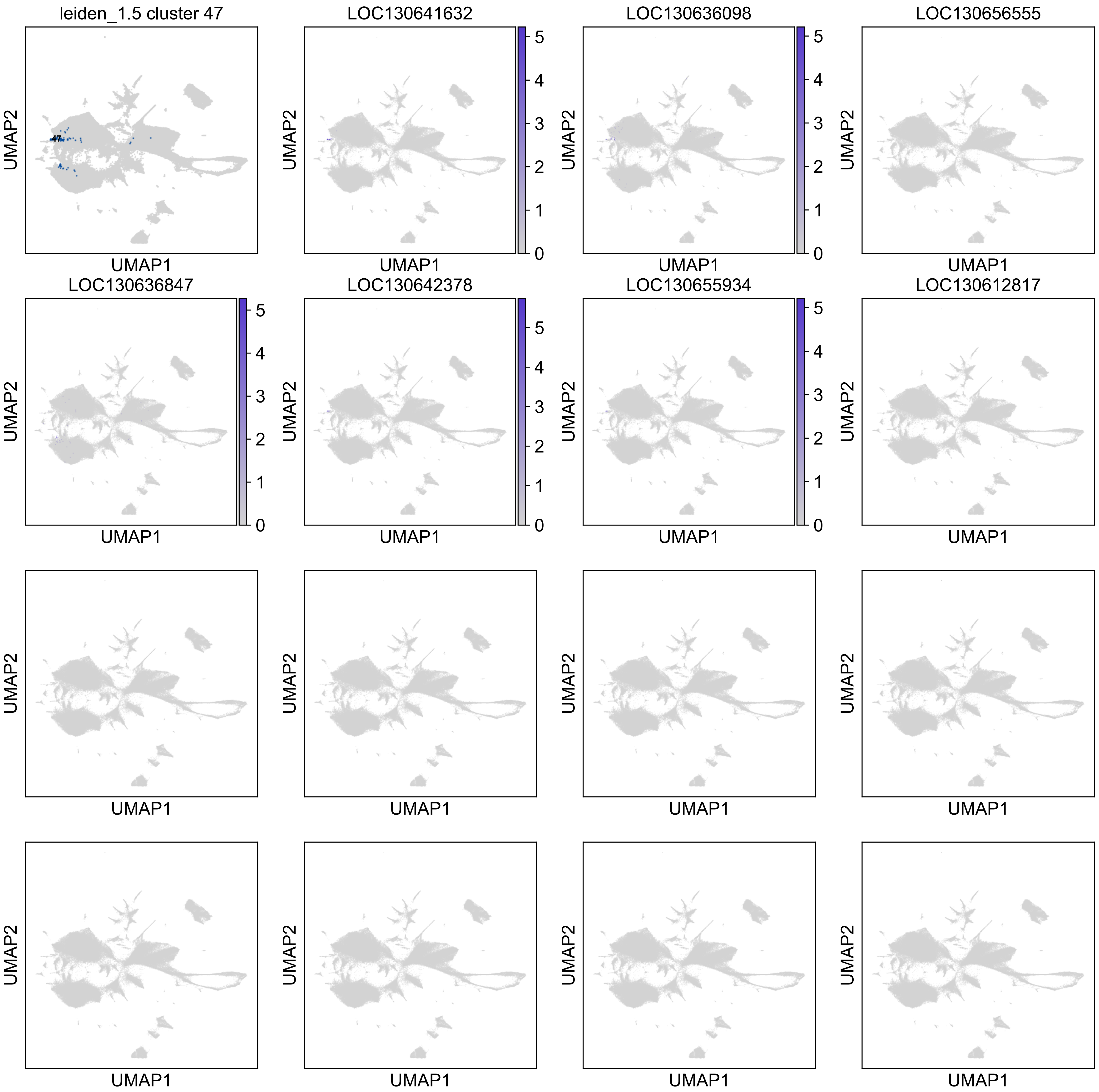
leiden_1.5 cluster 47
LOC130641632
LOC130636098
LOC130656555
LOC130636847
LOC130642378
LOC130655934
LOC130612817
UMAP1
UMAP2
47
0
1
2
3
4
5

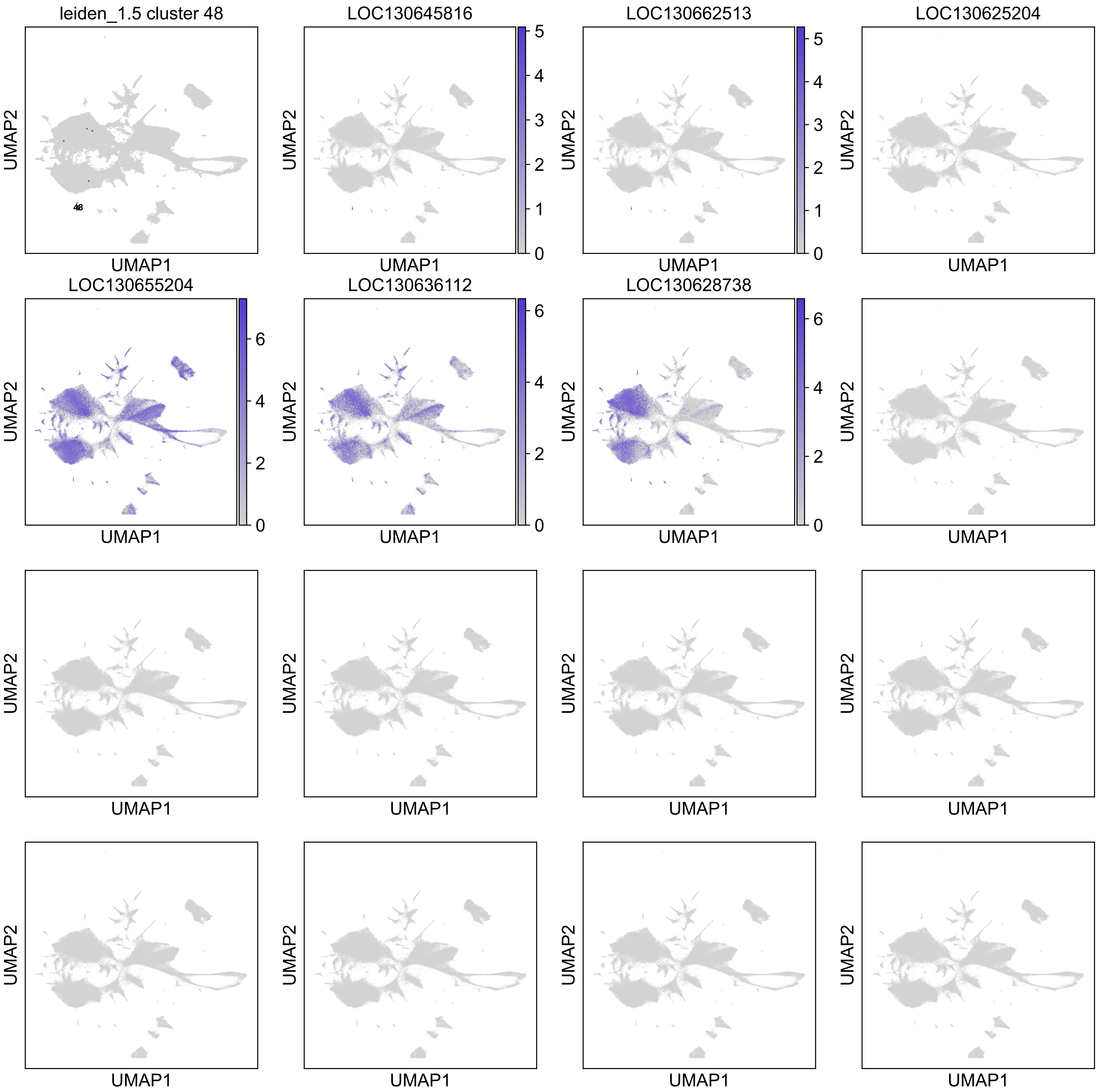
leiden_1.5 cluster 48
48
LOC130645816
LOC130662513
LOC130625204
LOC130655204
LOC130636112
LOC130628738
UMAP1
UMAP2
0
1
2
3
4
5
6

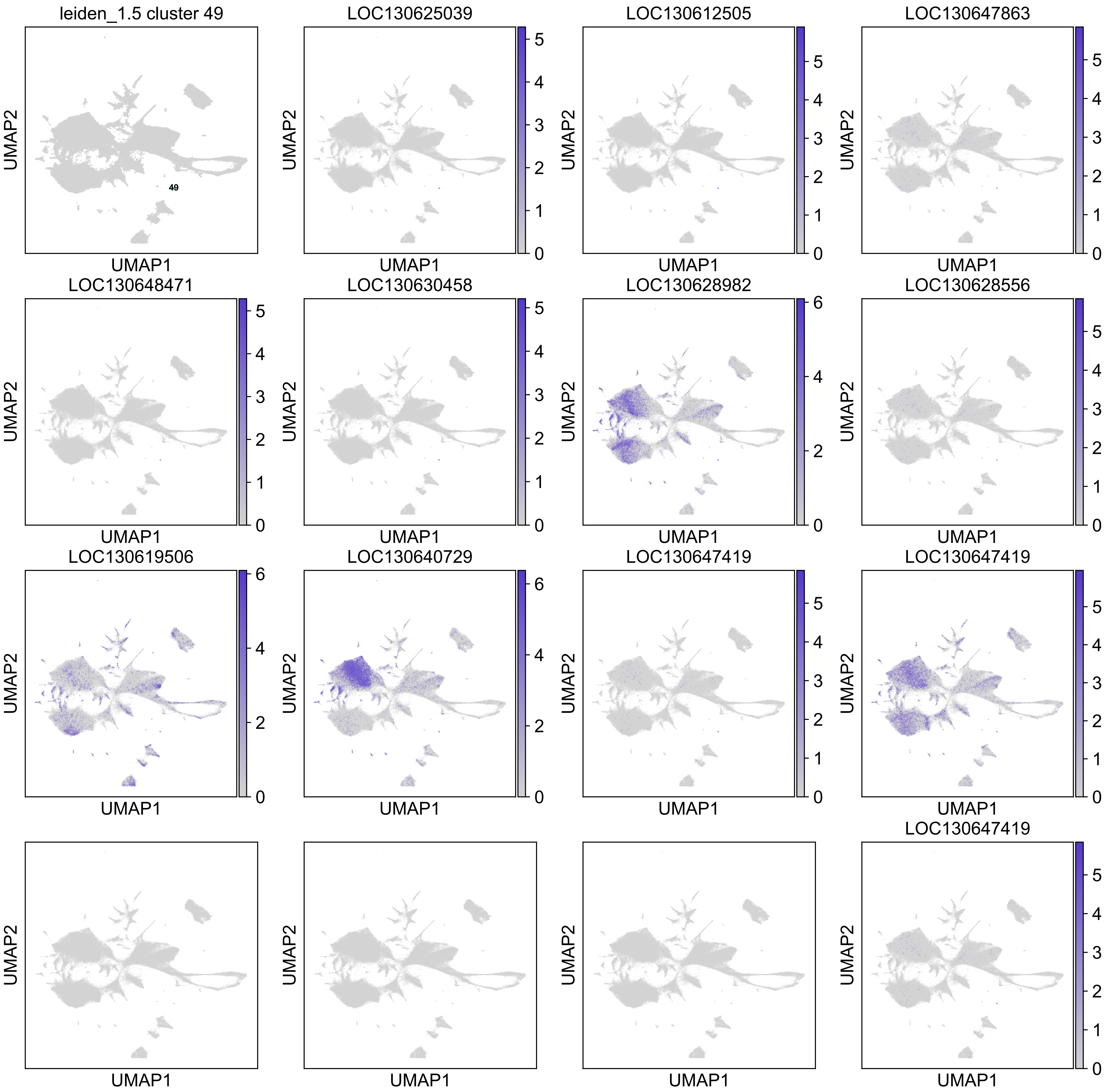

leiden_1.5 cluster 49
49
LOC130625039
LOC130612505
LOC130647863
LOC130648471
LOC130630458
LOC130628982
LOC130628556
LOC130619506
LOC130640729
LOC130647419
LOC130647419
LOC130647419
UMAP1
UMAP2

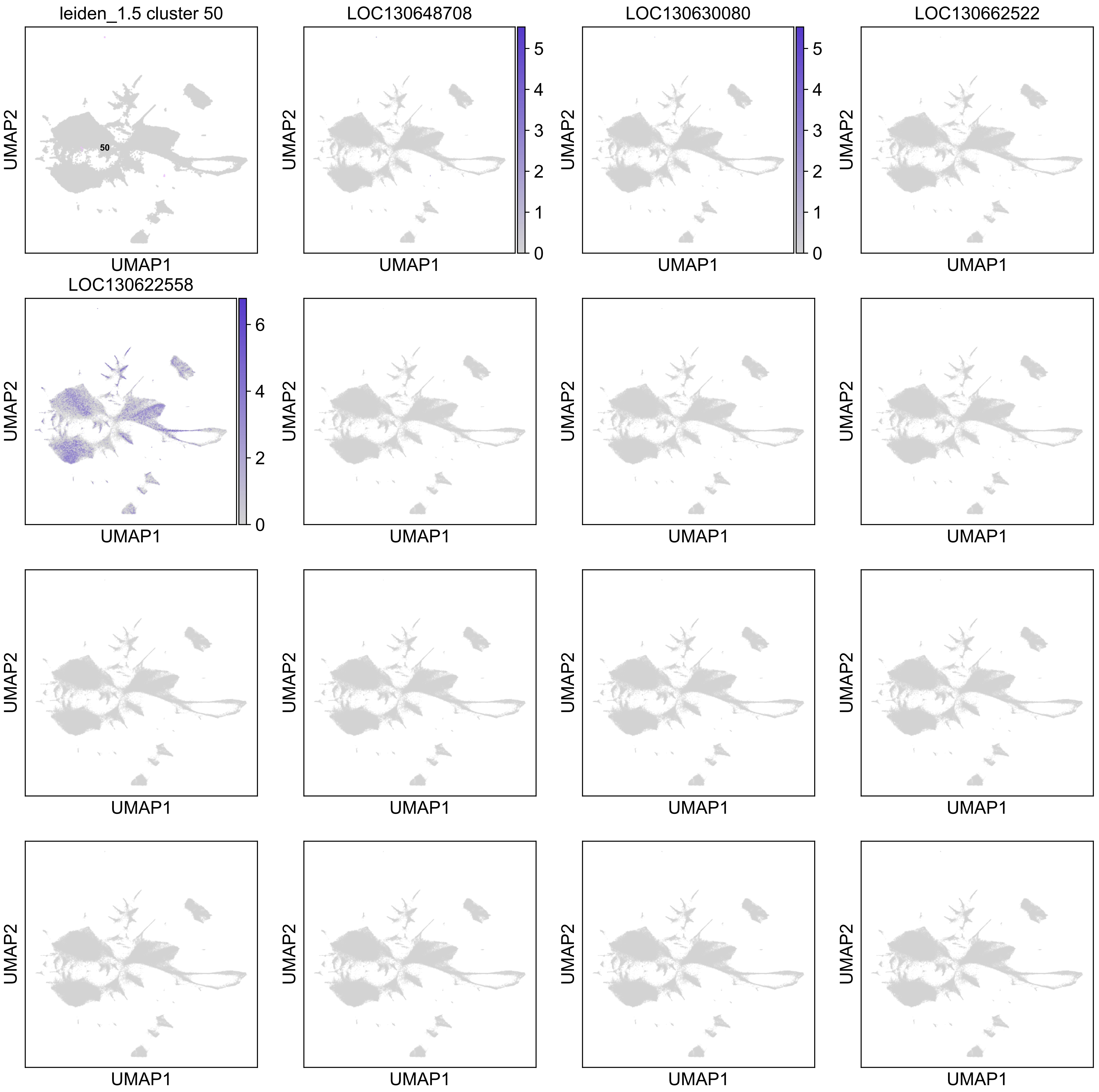
leiden_1.5 cluster 50
50
LOC130648708
LOC130630080
LOC130662522
LOC130622558
UMAP1
UMAP2
0
1
2
3
4
5
6

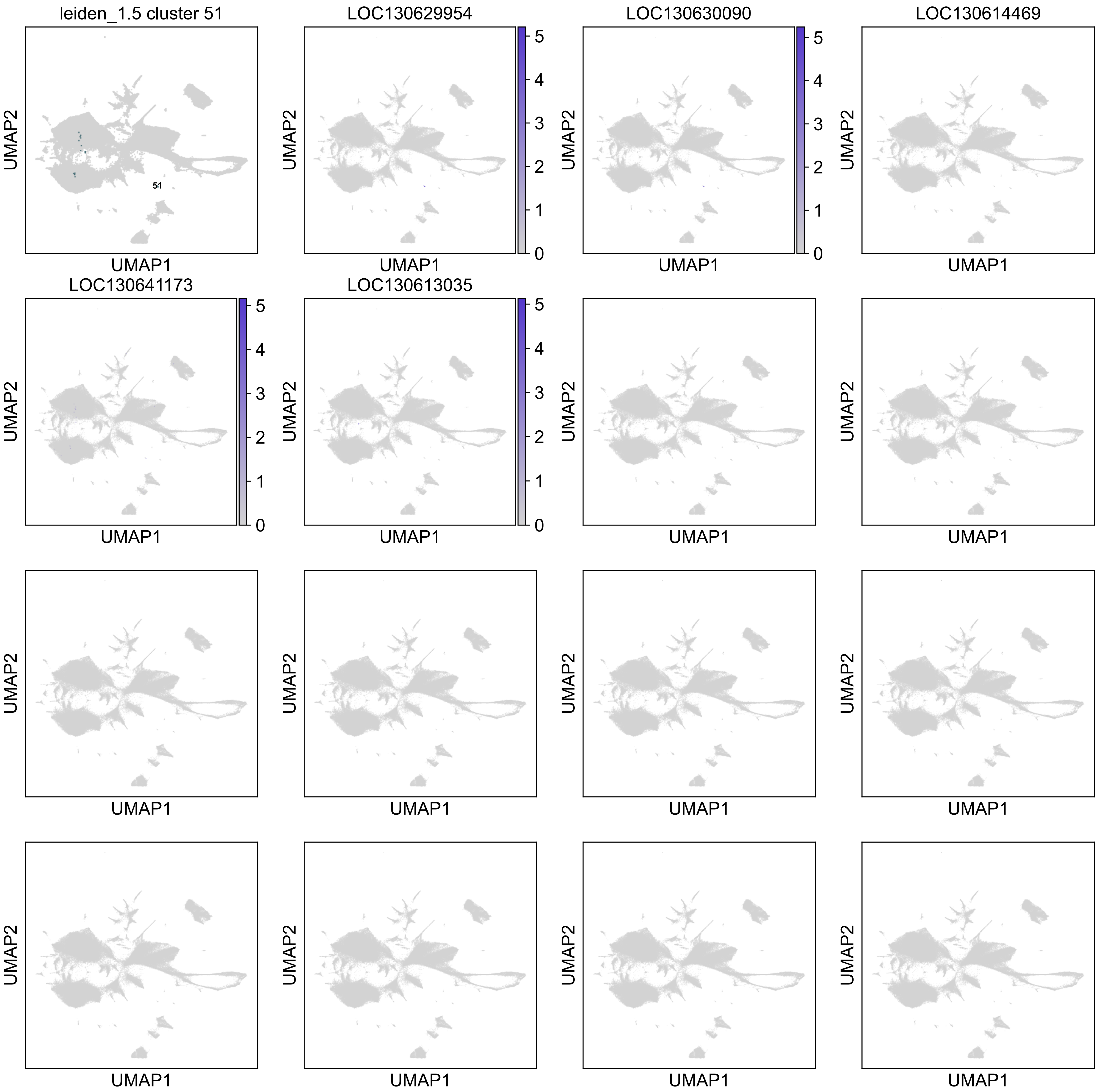
leiden_1.5 cluster 51
51
LOC130629954
LOC130630090
LOC130614469
LOC130641173
LOC130613035
UMAP1
UMAP2
0
1
2
3
4
5
